# *Aedes albopictus* and dengue transmission risk in France over the 21^st^ century

**DOI:** 10.64898/2026.03.06.710080

**Authors:** Andrea Radici, Pachka Hammami, Florence Fournet, Didier Fontenille, Cyril Caminade

**Affiliations:** MIVEGEC (Université de Montpellier, IRD, CNRS), F-34090 Montpellier, France; ASTRE (CIRAD, INRAE, Université de Montpellier), F-34398 Montferrier-sur-Lez, France; ICTP, I-34151 Trieste, Italy

**Keywords:** *Aedes albopictus*, dengue, climate change, vector ecology, epidemiological modelling

## Abstract

Climate change is dramatically affecting species distribution and phenology worldwide. Its impact on arthropod vectors, such as the *Aedes albopictus* mosquito, has important consequences for biting nuisance and arbovirus transmission risk. Here, we assess the impact of climate change on the presence and abundance of *Ae. albopictus*, as well as the risk of dengue transmission over France during the 21^st^ century. We use a mechanistic model that we adjusted against records of recent autochthonous cases of dengue in France. We simulate climatic suitability indicators, such as the adult abundance during the activity period, epidemic risk and secondary cases of dengue under different climatic and demographic scenarios at different periods up to 2085. Future simulations are based on a high-pressure scenario (high greenhouse gas emissions, high demographic growth) and a median-pressure scenario (median greenhouse gas emissions, demographic stagnation). To account for climate model uncertainty, we repeat the simulations for three different regional climate models. By 2085, in the high-pressure scenario, most of France (89-96%) will be climatically suitable for the establishment of *Ae. albopictus*, with the exception of mountain ranges. Similarly, autochthonous transmission of dengue will be theoretically possible in all colonized areas except over northern lowlands (71-95%). In the median pressure scenario, both climatic suitability for establishment (49-89%) and autochthonous dengue transmission risk (31-82%) exhibit large variation. Low population density areas show moderate suitability for vector establishment but exhibit the highest potential for dengue transmission. Overwintering mechanisms, such as egg diapause, indispensable for survival in temperate climates, may not be necessary along the Mediterranean and Basque coasts, allowing activity of the vector all year-round in the future.

## Introduction

Recent arboviral outbreaks in Western Europe, such as the Fano dengue outbreak that occurred in Italy in 2024 and the widespread Chikungunya epidemic that affected France in 2025, have brought the attention of the Asian tiger mosquito, *Aedes albopictus,* to citizens and public health institutions (Sacco et al., 2024; Santé Publique France, 2025). In the last decades, entomologists, epidemiologists and public health officers have been monitoring the concomitant progression of *Ae. albopictus* and arboviral cases in European countries(Farooq et al., 2025; Gossner et al., 2022; Wint et al., 2023). This mosquito, native to Asian tropical rainforest, has been colonizing Europe since the end of the 20^th^ century. According to the latest 2025 update by the European Centre for Disease Prevention and Control (ECDC), *Ae. albopictus* is currently established in 30 out of 45 countries in Europe. The Asian tiger mosquito is spreading northward in Belgium and Germany, and Italy, France, Spain and Croatia have been affected by autochthonous cases of dengue transmission in the last 15 years (ECDC, 2024a, 2024b).

Several factors contribute to such a rapid spread. Despite its short active flight range, this species has taken advantage of its anthropophilic behavior to being transported on both long and short distances (Marini et al., 2019). Thanks to globalization, dormant eggs have been moved in shipping containers from endemic countries to continents except Antarctica. This is how *Ae. albopictus* arrived in Albania first in 1979, and later in Italy (Sherpa et al., 2019). At shorter spatial scales, adult female mosquitoes have been hitchhiking on motorized vehicles (Eritja et al., 2017). Several studies showed how vehicular traffic contributed to the spread of this invasive species at subnational scale (Roche et al., 2015; Roques & Bonnefon, 2016).

Once a sufficiently large population of *Ae. albopictus* has reached a new site, its establishment depends on a number of environmental factors (Waldock et al., 2013). Among all factors favoring its establishment, climatic factors attracted the interest of the scientific community (Couper et al., 2025; Gizaw et al., 2024; Lafferty, 2009; Morin et al., 2013; Souza & Weaver, 2024). Because *Ae. albopictus* is a species native to tropical countries, characterized by warm and rather stable temperatures, and frequent rain, it underwent behavioral and physiological adaptations to establish in the European temperate climate, with cold winter and reduced water availability. Behavioral adaptations include the increased supply of human irrigation through man-made containers, which replaced natural water (Paupy et al., 2009); while physiological adaptation includes diapause, *i.e.* the process by which, when the daylight shortens below a critical photoperiod, differentiated eggs are laid. These diapausing eggs endure harsher winter conditions and will hatch subsequently in spring (Lacour et al., 2015). Diapause permits the persistence of *Ae. albopictus* population in temperate conditions, but warm seasons allow this population to grow and strengthen. As most ectotherms, these mosquitoes depend on external thermal conditions to sustain their life cycle (Delatte et al., 2009; Mordecai et al., 2019). At warm temperatures, mosquitoes develop faster, have longer life expectancy, shorten their gonotrophic cycles therefore laying more eggs in each period. Furthermore, viral transmission also benefits from warm climates: the higher the temperatures, the shorter the viral external incubation period of arboviruses (EIP; Morin et al., 2013).

For these reasons, increasing temperatures are reasonably regarded as one of the main global drivers affecting future dengue spread (Dorigatti et al., 2025; Mordecai et al., 2019). Several modelling studies have tried to estimate the impact of different climate scenarios on the distribution of *Ae. albopictus* (Brady et al., 2014; Colón-González et al., 2021; Kraemer et al., 2019; Pardo-Araujo et al., 2024) and the related risk of dengue transmission (Liu-Helmersson et al., 2016). For example, in cold temperate areas, milder winters may favor overwintering, and warmer summers the growth of the mosquito population and the transmission of arboviruses. It is the case for the British Isles, where large southern regions will likely become suitable for this vector by 2060 (Metelmann et al., 2019). Radici et al. (2025a) suggested that the recent temperature increase sustained its colonization in Hexagonal France. In warm temperate regions, higher temperatures in spring and autumn may extend the activity season of the vector in future, up to overlap to the season when arboviruses circulate in the southern Hemisphere. Pardo-Araujo et al. (2024) suggested that in some regions of Spain, mosquitoes will be active in every season in a few decades. In Southern Europe, mild winter may allow an homodynamic (year-round) activity of the Asian tiger mosquito (Del Lesto et al., 2022).

On the other hand, climate change is a complex phenomena, not limited to global warming (Morin et al., 2013; Souza & Weaver, 2024). Change in precipitation may impact water availability for the immature stages of the mosquito (Gizaw et al., 2024). Summer temperature extremes may exceed the heat tolerance of mosquitoes in the warmest places, causing population decline (Garrido Zornoza et al., 2024) and resulting in local decrease in the burden of dengue (Kraemer et al., 2019). The same climate extremes also affect host behavior, for example by modifying their daily exposition to the vector (Reiter et al., 2003). Moreover, anthropic changes are further influencing mosquito population, by altering urban infrastructures affecting the availability of breeding or rest sites (Clauzon et al., 2025; Mercat et al., 2025), and dengue transmission risk by modifying host availability.

Metropolitan France (the mainland and the island of Corsica, referred to as “France” in the following), where the colonization of *Ae. albopictus* is rapidly expanding since its first establishment in 2004, is a particularly interesting case study. Due to its broad climatic gradient including mediterranean, oceanic, continental and alpine climates, this country acts as a bridge between warm southern and cold northern Europe. In addition, given the movement of goods and persons with its overseas territories, the French population is exposed to a relatively high number of introductions of dengue cases, closely monitored by the national Health institutions. So far, other studies have already investigated the future of dengue and *Ae. albopictus* in Europe (Colón-González et al., 2021; Kraemer et al., 2019; Pardo-Araujo et al., 2024), but no study has focused on France using a multi-model, multi-scenarios approach based on high-resolution projections.

Here, we use high-resolution projections (about 8km) to dynamically simulate the impact of climate and demographic change on *Ae. albopictus* presence/abundance and the associated risk of dengue transmission in France over the period 2025-2085. We focus on dengue instead of Zika and chikungunya since it is the most widespread arbovirus worldwide and it has been causing local cases in France every year since 2010. Simulations are carried out using a dynamic demographic-epidemiological model that was adjusted based on recent local dengue data. We define two climatic and demographic scenarios: a high-pressure scenario (high greenhouse gas emissions, high demographic growth) and a median-pressure scenario (median greenhouse gas emissions, demographic stagnation). Ecological-entomological indicators (suitability for establishment, density of adult females during the activity period) and epidemiological ones (duration of the dengue transmission season, dengue secondary cases) are derived for each scenario. To account for climate model uncertainty and to standardize climate change simulations at national scale, we carry out simulations based on three different climate models with different climate sensitivities (coldest, median and warmest models for a given scenario). These simulations will guide local, regional and national public health agencies to take actions against this mosquito nuisance and the risk of dengue transmission in order to initiate timely and targeted prevention and control measures.

## Material and methods

### Climatic and demographic data

We retrieved climatic projections for France from the DRIAS repository (Lémond et al., 2011; Soubeyroux et al., 2020). Calibrated Regional Climate Model (RCM) projections, available at a resolution of 8 km (the SAFRAN grid; Bertuzzi et al., 2024) for the period 1950-2100, included average, maximum and minimum daily temperature (°C), as well as cumulated daily precipitation (converted to mm/d) for different climatic scenarios, the so-called Reference Concentration Pathways (RCP). These RCM simulations were carried out within the EURO-CORDEX project framework (Jacob et al., 2020). To account for uncertainties related to different climate sensitivities, we considered outputs from three different RCMs among those indicated in the TRACC project by the French weather services (“Trajectoire de Réchauffement de référence pour l’Adaptation au Changement Climatique”; MétéoFrance, 2024). These three RCMs are driven by observed (for the historical baseline) and projected (for the future) greenhouse gas emissions, and correspond to a rapid, median and slow warming under a high emission scenario (RCP8.5). These simulations are based on an ensemble of global circulation models driving high-resolution regional circulation models for Europe, which we refer to as “warm”, “intermediate” and “cold” models (Table 1). To deal with multi-model outputs, simulated values will be reported as: “median (lowest – highest value)” in the Results section.

**Table 1.**
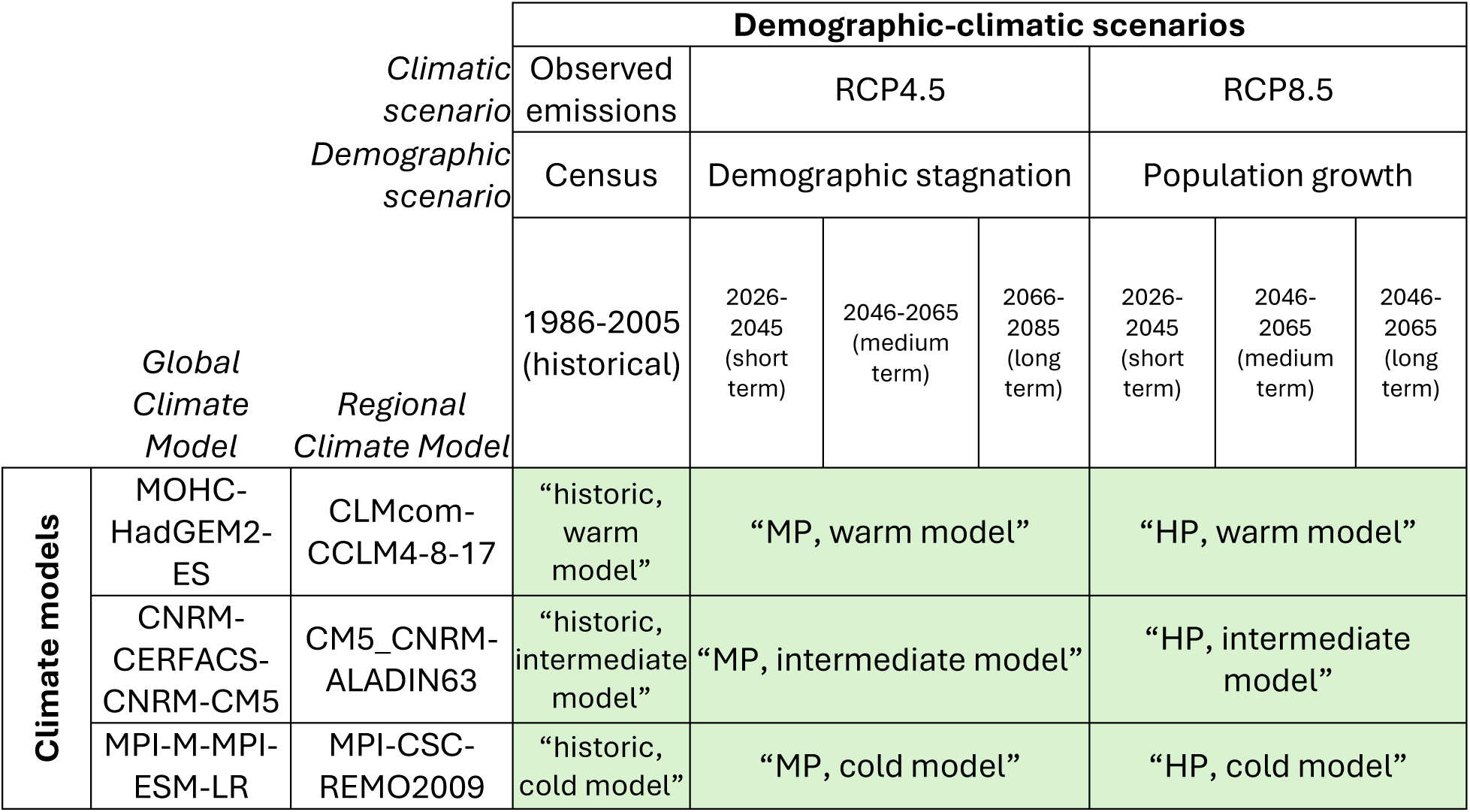
Summary of the demographic-climatic scenarios (in green) by crossing demographic scenarios, climatic scenarios, climatic models at different time horizon.

Because of the lack of demographic projections available at high spatial resolution to fit the most recent IPCC scenarios (Gao, 2017; Olén & Lehsten, 2022; Wang et al., 2022), we used estimates from the Omphale model, provided by the French national institute of statistics (INSEE). Omphale provides department-level (corresponding to EU NUTS3 administrative levels) population projections for each age class in the period 2018-2070 for different demographic hypothesis (INSEE, 2022). We considered the 2018 census, which is used by Omphale as the baseline, to redistribute the department-level future projections at the municipality level, and successively computed population densities (inhab./km²) on the same grid as the climatic projections (INSEE, 2025). In the Results, when mentioning a site in France, we will refer to the ∼64 km² grid cell containing/contained by that site. Population densities of the 1999 census were used as the historical baseline (Table 1).

We selected and aggregated climatic (two out of the four RCP-IPCC scenarios) and demographic (two out of eleven scenarios by Omphale) projections to find a compromise between the level of details and computational intensity. We therefore focused on two contrasted demographic-climatic scenarios, *i.e.* the High Pressure (HP) scenario, which combines the highest emission scenario (RCP8.5) with the highest migration and fertility scenario (“hypothèse haute” of the Omphale model) and the Median Pressure (MP) scenario, which consider median emissions (RCP4.5) and demographic stagnation (“hypothèse centrale”). We compared these scenarios with the historical pre-*Aedes albopictus* establishment period (1986-2005, with 1999 population data). Future time slices include the near future (2026-2045, with 2035 population projections), middle of century (2046-2065, with 2055 population projections) and end of century (2066-2085, with 2070 population projections; see Table 1).

### Dynamic vector-host model overview

We based our modeling approach on a simplified version of the mechanistic framework presented by Metelmann et al. (2019, 2021) which combines a mosquito population model (eggs *E*, diapausing eggs *E*_*D*_, larvae and pupae as juveniles *J*, unfed immature female adults *I*, susceptible female adults *A*_*s*_) with an epidemiological model (exposed female adults *A*_*E*_, infected female adults *A*_*I*_, susceptible humans *H*_*S*_) to estimate the number of secondary cases produced by an introduction of an imported case of dengue (Figure 1) in a given location *x*:

**Figure 1.**
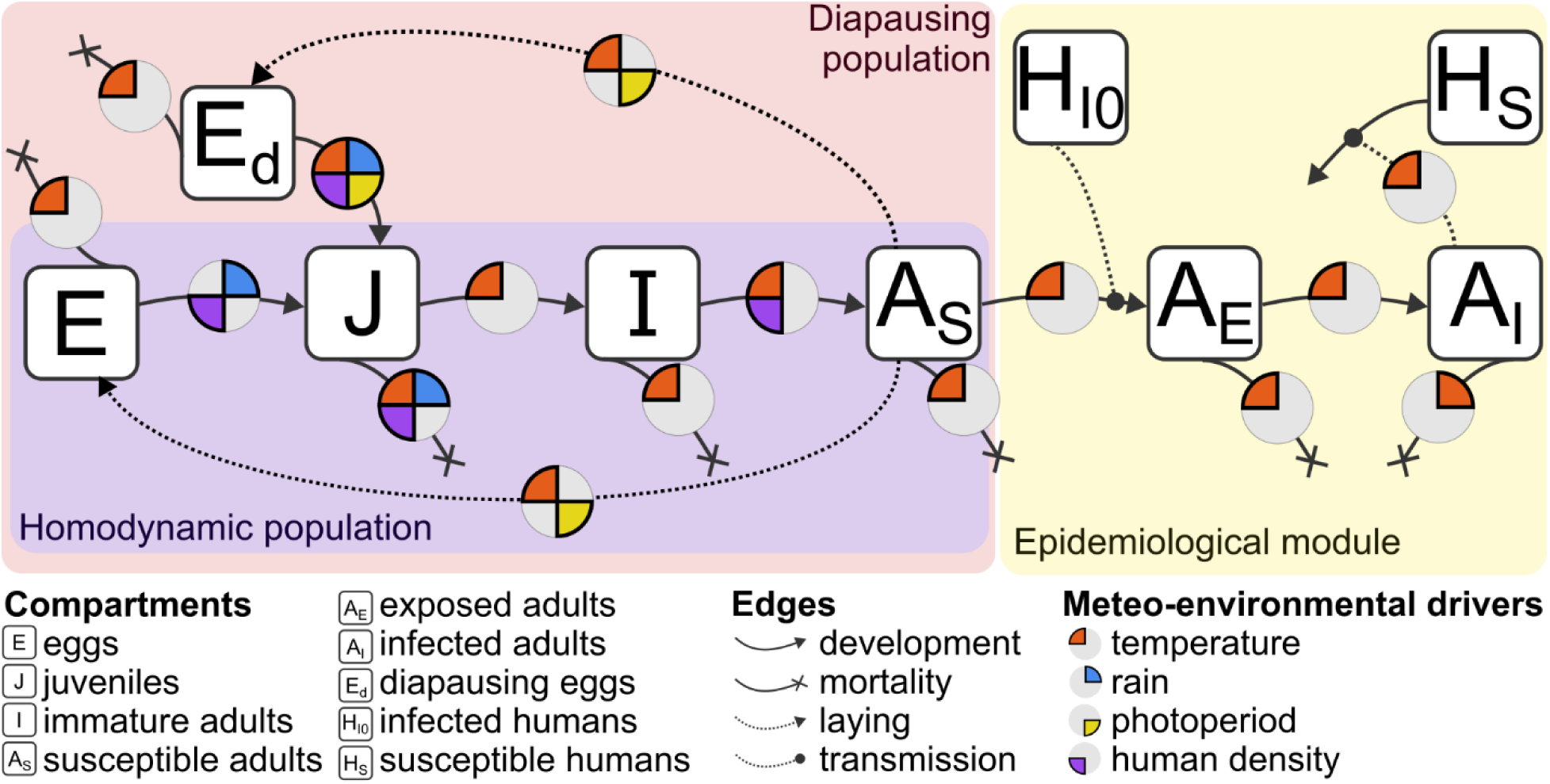
Population model schematic for Ae. albopictus (left). The epidemiological module to estimate dengue secondary cases is depicted on the right-hand side.

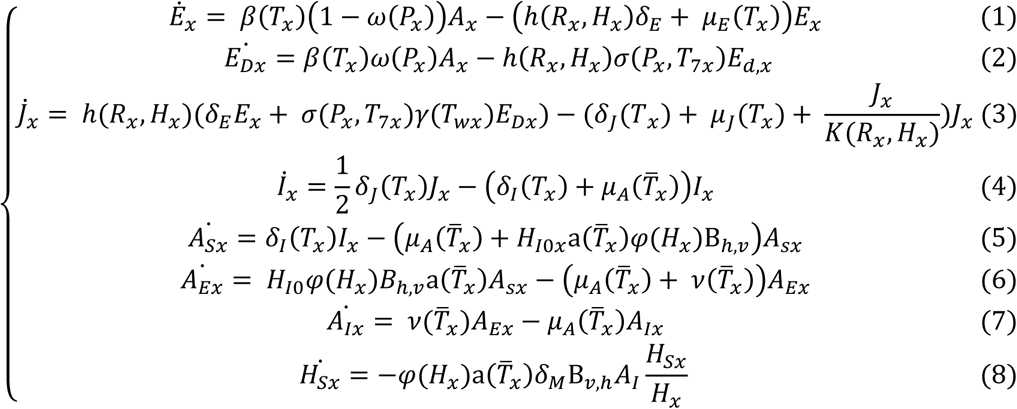

In the population module (Eq. 1-5), the local instantaneous temperature *T*_*x*_ regulates the rate of development of larvae and pupae (*δ*_*J*_) and immature adults (*δ*_*I*_), as well as mortality of eggs (*μ*_*E*_) and juveniles (*μ*_*J*_) and fertility (*β*), while mean daily temperature *T̅*_*x*_ determines the mortality rate of adults (*μ*_*A*_). The photoperiod *P*_*x*_ regulates the type of laid eggs (*ω*) and the spring hatching (*σ*) which also depends on the average temperature of the previous week *T*_7*x*_. Note that, if *ω* is fixed to 0, the same system of differential equations can simulate the population dynamic in absence of diapause (homodynamicity). Minimal winter temperature *T*_*wx*_ determines the fraction of viable diapausing eggs *γ*. Water availability of both natural (rainfall *R*_*x*_) and anthropic (human density *H*_*x*_) origin, regulate eggs hatching *h* and the density-dependent carrying capacity *K* (Metelmann et al., 2019).

In the epidemiological module (Eq. 5-8), female adults get exposed to dengue depending on their temperature-dependent biting rate *a*, multiplied by the coefficient of host preference for humans *φ*, the prevalence of infected humans representing imported cases *H*_*I*0*x*_, and the probability of virus transmission (human to vector) per bite *B*_*h*,*v*_, becoming infectious after the EIP (reciprocal of *v*). Similarly, secondary human cases get exposed to the virus with probability *B*_*v*,*h*_ per bite; *δ*_*M*_ is a scaling factor representing the maximum number of people that can be infected by a single mosquito (Metelmann et al., 2021; details about parameters in Table SI1).

With respect to version by Metelmann et al. (2021), the presented epidemiological module relies on ordinary differential equations. This simplification is justified by our focus on secondary infections at the end of the mosquito activity season, rather than on the complex chain of multiple reciprocal vector-host transmission.

### Epidemiological data to adjust the larval carrying capacity

Simulated secondary cases 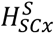 can be compared to reported autochthonous cases 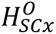 in France. They are estimated as a fraction of the recovered humans, i.e. the difference between susceptible *H*_*Sx*_(*t*_1_) at the end of a simulation *t*_1_ against their initial value *H*_*x*_ = *H*_*Sx*_(*t*_0_):

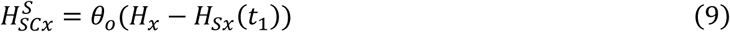

Where *θ*_*o*_ = 0.4 is the fraction of symptomatic (and therefore detectable) cases (Santis et al., 2023).

We based on secondary cases to adjust the carrying capacity *K*. The original model implies a direct proportionality between *K* and *H*_*x*_, while observations suggest that this relationship should not be linear; densely populated areas have vertical buildings which provide less per capita larval habitat, and usually reduced urban green spaces (Mercat et al., 2025). Therefore, we proposed an alternative formulation of *K*(*R*_*l*_, *H*_*l*_) by introducing the parameter *ε*:

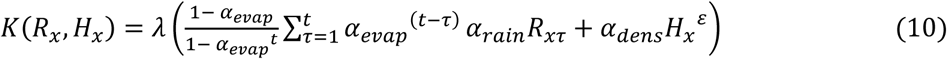

where the value of *ε* ≤ 1 decreases the value of human density (*H*_*τ*_), which we calibrate in the following section. In the original model formulation, *ε* = 1; when *ε* → 0, the contribution of additional population in providing breeding sites in already densely populated areas becomes less important.

To calibrate the value of *ε*, we used observed autochthonous dengue cases 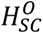 collected by the National Health Institute in 2024 and 2025. Only data with at least 5 cases were considered (Santé Publique France, 2024, 2025). We assumed that these cases were secondary and arising from the introduced case, i.e. that no further transmission occurred following mosquito control interventions. This assumption is consistent with a scenario for which timely and extensive vector control interventions were conducted after the detection of a positive case (Ministère des Solidarités et de la Santé, 2019). We simulated the number of secondary cases 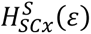 for an outbreak that occurred in *x* depending on parameter *ε*. MétéoFrance data from weather stations were used to run the model (https://www.data.gouv.fr/organizations/meteo-france). Eventually, we selected the value of *ε* that minimizes the Root Mean Square Logarithmic Error *RMSLE*(*ε*, *ϑ*) with observations 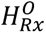:

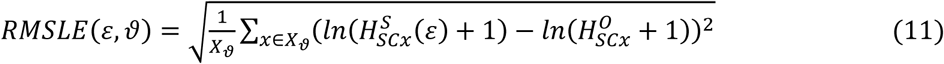

Where *X*_*ϑ*_ represents either the number of sites where an outbreak occurred in 2024, used for calibration (*ϑ* = “calibration”), or the number of sites where an outbreak occurred in 2025, used as validation (*ϑ* = “validation”). We chose *RMSLE* over other metrics to prioritize a correct reproduction of the order of magnitude of secondary cases that might vary by factors of 10. The. The empirical distribution of *ε* was estimated using the Non-Parametric Bootstrap procedure with 10^6^ Monte Carlo resamples of the observations.

### Entomological and epidemiological indicators

With the *ε*-adjusted model, for each scenario, time period and climate model, we simulated the following indicators from the literature (Table 2):

- **Climatic suitability for the establishment of a diapausing population**. Climatic suitability is computed as the growth rate of a small population within a year. It is computed via the indicator *E*_0*xy*_, i.e. the number of diapausing eggs at the end of the year *y*, when the population at location *x* is initialized to *E*_*Dx*_(0) = 1 *eggs*/*ha*. *E*_0*x*,*y*_ is eventually aggregated using the geometric mean over different time periods.
- **Average density of adult mosquitoes during the activity period**. In this case, after simulating a five-year spin-up period, we let the mosquito population evolve for the considered time period and calculate the annual mean of adults densities *A̅*_*x*_ from the 1^st^ of May (*t* = 60 *days*) to the 31^st^ of October (*t* = 304 *days*).
- **Length of the transmission season of dengue**. Based on the adult density simulations, we estimate the basic reproduction number for dengue (Caminade et al., 2017; Zardini et al., 2024):

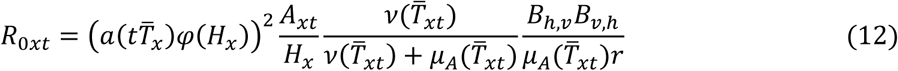

and derive the subsequent indicator of epidemic risk, the Length of the Transmission Season (LTS), i.e. the number of days for which *R*_0_ > 1.
- **Dengue secondary cases from introduction at peak transmission**, are estimated using Eq. 8. To fix an introduction date, we first calculated the median date 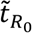 of the highest *R*_0_value for each scenario, period and climate model. Then we assumed the introduction of one infected case (*i* = 1) at 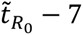 (roughly the order of magnitude of the incubation period) and let an epidemic evolve until the end of the year to estimate secondary cases.
- **Climatic suitability for the establishment of a non-diapausing population**. Analogously to *E*_0*xy*_, *A*_0*x*(*y*−1,*y*)_ is a measure of yearly growth rate of a small population by removing the possibility of diapause (*ω*(*P*_*x*_) = 0). In this case, no diapausing eggs are laid (*E*_*Dx*_(*t*) = 0); yearly growth rate is then calculated as the ratio between adult density at a given date (the 1^st^ of July, i.e. the 212^th^ day of the year) from a year (*y*) to the previous (*y* − 1), when the population is initialized to *A*_*x*_(*t* = 212 *days*, *y*) = 10. Values are then aggregated using the geometric mean for each scenario/climatic model/time horizon.

**Table 2.** Summary of the simulated entomological and epidemiological indicators.

| Indicator | Type | Description | Formulation |
| --- | --- | --- | --- |
| $E_{0x}$ | Entomological | Climatic suitability for the establishment of a diapausing population | $E_{0xy} = \frac{E_{dx}(t = 365, y)}{E_{dx}(t = 0, y) = 1} \quad (13)$ |
| $\bar{A}_x$ | | Average density of adult mosquitoes during the activity period | $\bar{A}_{xy} = \frac{\sum_{t=60}^{304} A_x(t)}{304 - 60 + 1} \quad (14)$ |
| $A_{0x}$ | | Climatic suitability for the establishment of a non-diapausing population | $A_{0x(y-1,y)} = \frac{A_x(t = 212, y)}{A_x(t = 212, y - 1) = 10} \quad (15)$ |
| $LTS_x$ | Epidemiologic<br>al | Length of the<br>transmission season of<br>dengue | $LTS = \sum_{t \in y} (R_{0x}(t) > 1) \quad (16)$ |
| $H_{SCx}$ | | Dengue secondary cases<br>from introduction at peak<br>transmission conditions | $H_{SCx} = \theta_0(H_x - H_{Sx}(t = 365)) \quad (17)$<br>(introduction in $\tilde{t}_{R_0} - 7$) |

## Results

### Carrying capacity adjustment

In 2024 and 2025, respectively 83 and 29 autochthonous dengue cases in France were notified. All the six major dengue outbreaks occurred in the Provence-Côte d’Azur-Alps region, in the Southeast. In 2024, the unadjusted carrying capacity model systematically overestimated the number of secondary cases; for example, in Vallauris, where 14 cases occurred late in the summer, 158 cases were simulated that year. The correction to the dependency between carrying capacity and human density, obtained with a value of *ε* = 0.6 (interquartile interval: 0.55, 0.65), allowed an estimation of the correct order of magnitude (10.4; Figure 2). The relevance of this correction was confirmed by simulating secondary cases in 2025 as validation; in Rognac, where 5 local cases occurred, the adjusted model decreased the estimated secondary cases from 60.3 to 7.6.

**Figure 2.**
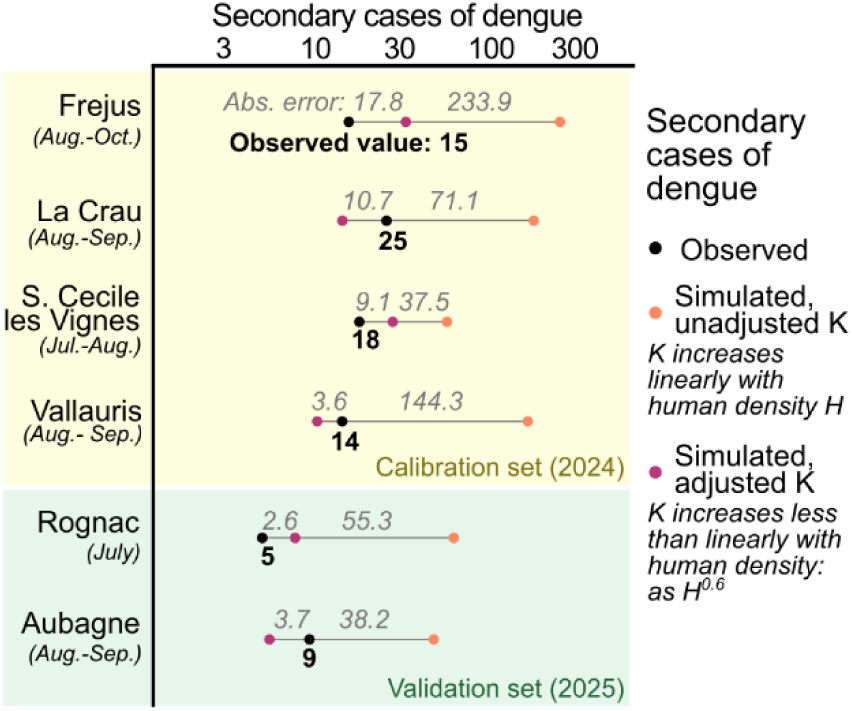
Observed and simulated (with both adjusted and unadjusted immature carrying capacity, and relative errors) secondary dengue cases for recent outbreaks in France (2024 – 2025)

This calibration allowed us to re-simulate the epidemic by letting the dengue outbreak extinguish naturally. Depending on the site, the early end of dengue transmission permitted a reduction of the incidence between 33% and 91% (55% on average; Tab SI3).

### Projected demographic and climatic change

In 1999, the population in France was about 58.3 million people (Figure 3a). Both demographic scenarios show an increasing trend, concentrated in large cities and suburbs. In the HP scenario, population reaches 68.4 million people in 2035 and keep growing to 75.2 million people in 2070. Large population values are still concentrated in large cities, along the western Atlantic side and in Mediterranean regions. By contrast, the MP scenario predicts a population increase in already densely populated areas, counterbalanced by a depopulation of rural areas, especially in the northeastern regions; therefore the population stabilizes around 66-65 million people already by 2035.

**Figure 3.**
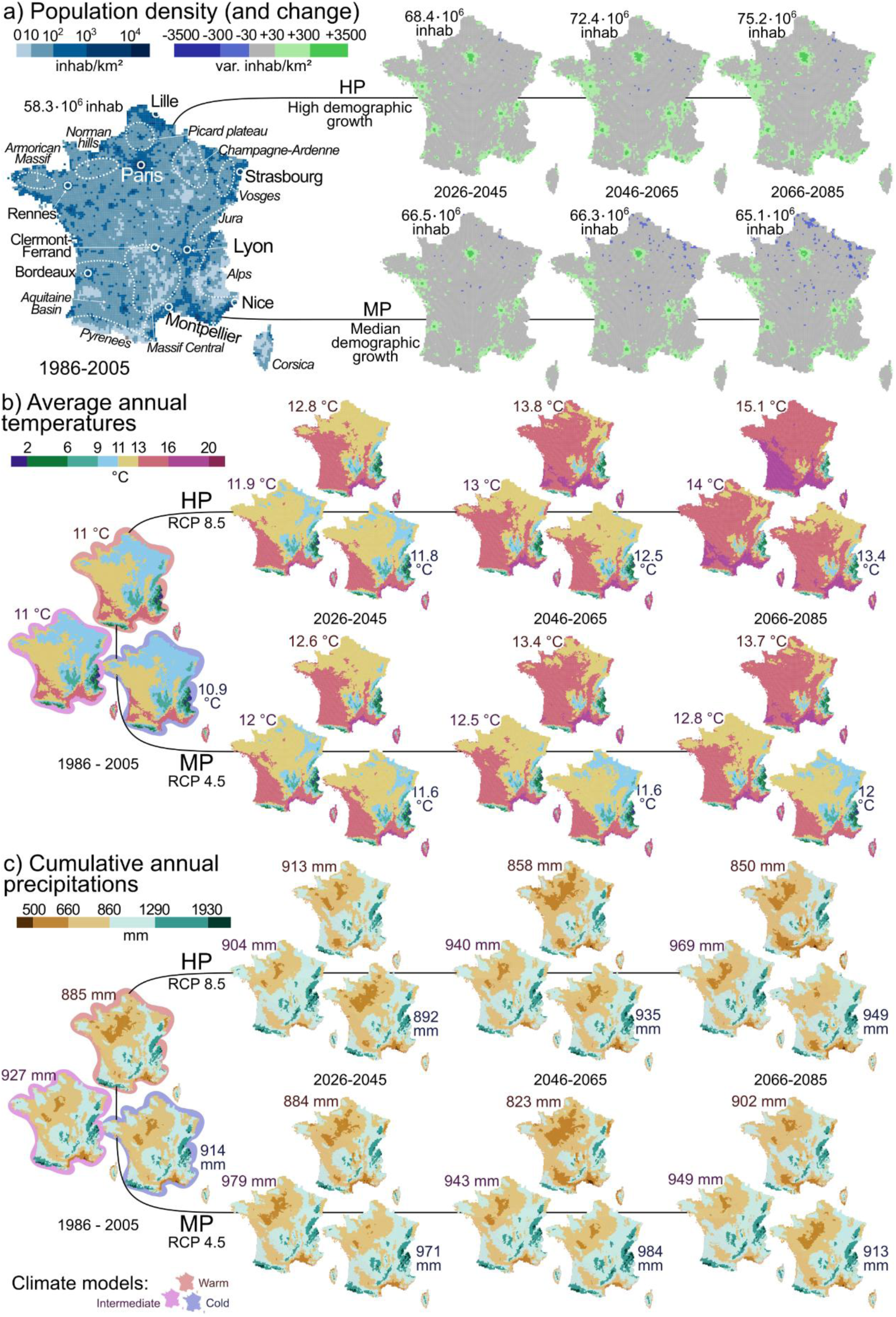
Summary of changes in key variables (population, average annual temperature, cumulative precipitation) depending on scenario, time period, and climate model.

Temperature is simulated to significantly increase in future over France (Figure 3b). During the historical period (1986-2005), average temperature values were about 11 °C. The warmest areas are located in the South, on the Mediterranean coasts and the Aquitaine Basin. The coldest areas correspond to northeastern France and the mountain ranges in the Alps, the Massif Central and the Pyrenees. Under the RCP8.5 scenario, these temperatures are simulated to increase by 0.9 °C (0.8–1.8) in 2026-2045 and up to 3 °C (2.4– 4.1 °C) in 2066-2085. Under the RCP4.5 scenario, temperature increases (+1 °C; 0.6–1.6 °C) are in a similar range with respect to RCP8.5 values in 2026-45, but are lower (+1.8 °C) toward the end of the century (1–2.7°C in 2066-2085).

Simulated future changes in annual rainfall are more uncertain (Figure 3c). Differences between climate models are larger than the differences between scenarios (the warm and the intermediate model being respectively the driest and the rainiest 6 times out of 7). Simulated average rainfall values in France range between 915 ± 65 mm; rainfall is higher over mountain ranges and drier conditions are simulated over the Mediterranean coasts and in the inner northwestern regions.

### Projected change in spatial suitability for *Ae. albopictus*

During the twenty years preceding the establishment of the Asian tiger mosquito in France, about 24% (20–27%) of the territory was already suitable (E_0_>1) to host this invasive species (Figure 4). Strongly suitable regions (E_0_>10) were located on the Mediterranean coasts, in large southern cities and Paris. Under the HP scenario, this percentage is increasing from 46% (41–57%) in the short term (2026-45) up to 92% (89–96%) in the long term (2066-85). Importantly, most of France is expected to become suitable for the establishment of *Ae. albopictus* for the worst case scenario (warm model, RCP8.5) in 2066-85 apart from the Alps and the Pyrenees. In the cold model, minor mountain ranges (the Massif Central, the Jura, the Vosges) are also simulated to be mosquito-free. In the short term, suitable areas expand faster under the MP scenario, exceeding half of the territory in 2026-45 (56%; 53–75%), and representing about 72% of the territory by 2066-85 (with high variability: 49–89%). In addition to the mountainous mosquito-free regions, inner rural areas are also remaining mosquito-free in the MP scenario in future (such as the Ardenne-Champagne region, the Picard plateau, the Armorican massif and the Norman hills; details on specific sites in Tab. SI4SI4).

**Figure 4.**
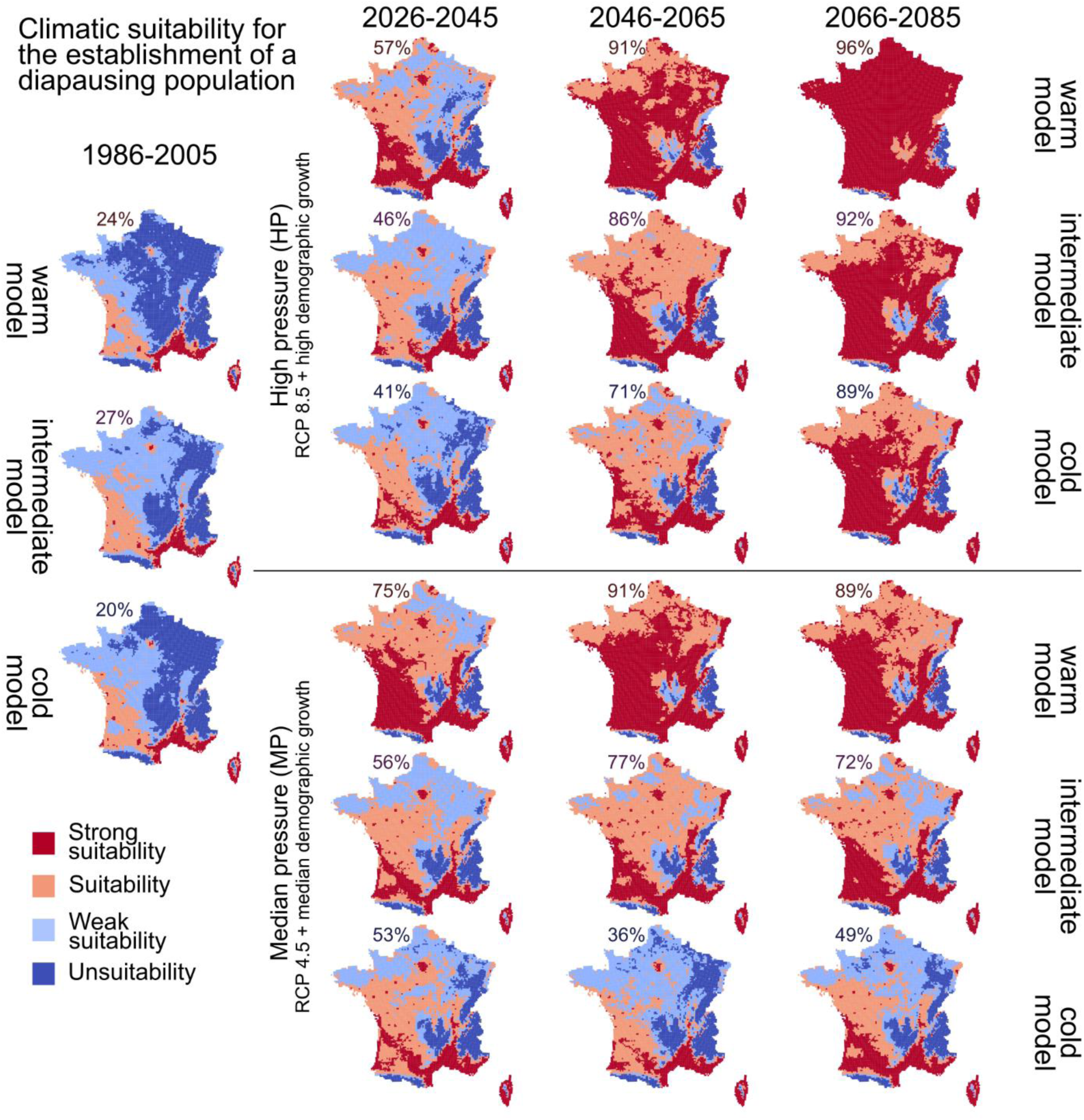
Climatic suitability scenarios for the establishment of Ae. albopictus. Strong suitability: E_0_>10; Suitability: E_0_>1; weak suitability: E_0_>0.1; unsuitable otherwise. % of territory simulated to be suitable (E_0_>1).

### Projected change in the duration of the dengue transmission season

Areas at risk of dengue transmission (LTS ≥ 7 days, or number of weeks for which R_0_ for dengue ≥ 1, Figure 5) represent a subset of those climatically suitable for the establishment of the vector (Figure 4 and SI1). Historically, these areas are in the southern part of the country, as well as in Corsica, and correspond to about 12% (10–14%) of France. Under the HP scenario, the area suitable for dengue transmission expand progressively, reaching 24% (22–62%) of the country in the short term and 80% (71%–95%) in the long term. In the warmest model, most of France becomes suitable for dengue transmission for at least 8 weeks per year in future; however, the duration of the dengue transmission season starts to decrease in the southernmost regions (in 2066-2080, the extent of the area where LTR>15 weeks for the warm model is smaller compared to the intermediate one, where transmission is possible for more than 23 weeks a year in 1% of the territory). Under the MP scenario, simulated changes in dengue transmission risk strongly depends on the climate model. Following a sharp acceleration, the area suitable for dengue transmission reach 35% (31–57%) of the territory in the short term, its progression slows in the rest of the century (47%; 31–82%); in the warm model, the MP scenario almost overlaps with the HP scenario. Dengue-free areas match the mosquito-free areas, but include also densely populated regions (details on specific sites are provided in Tab. SI4).

**Figure 5.**
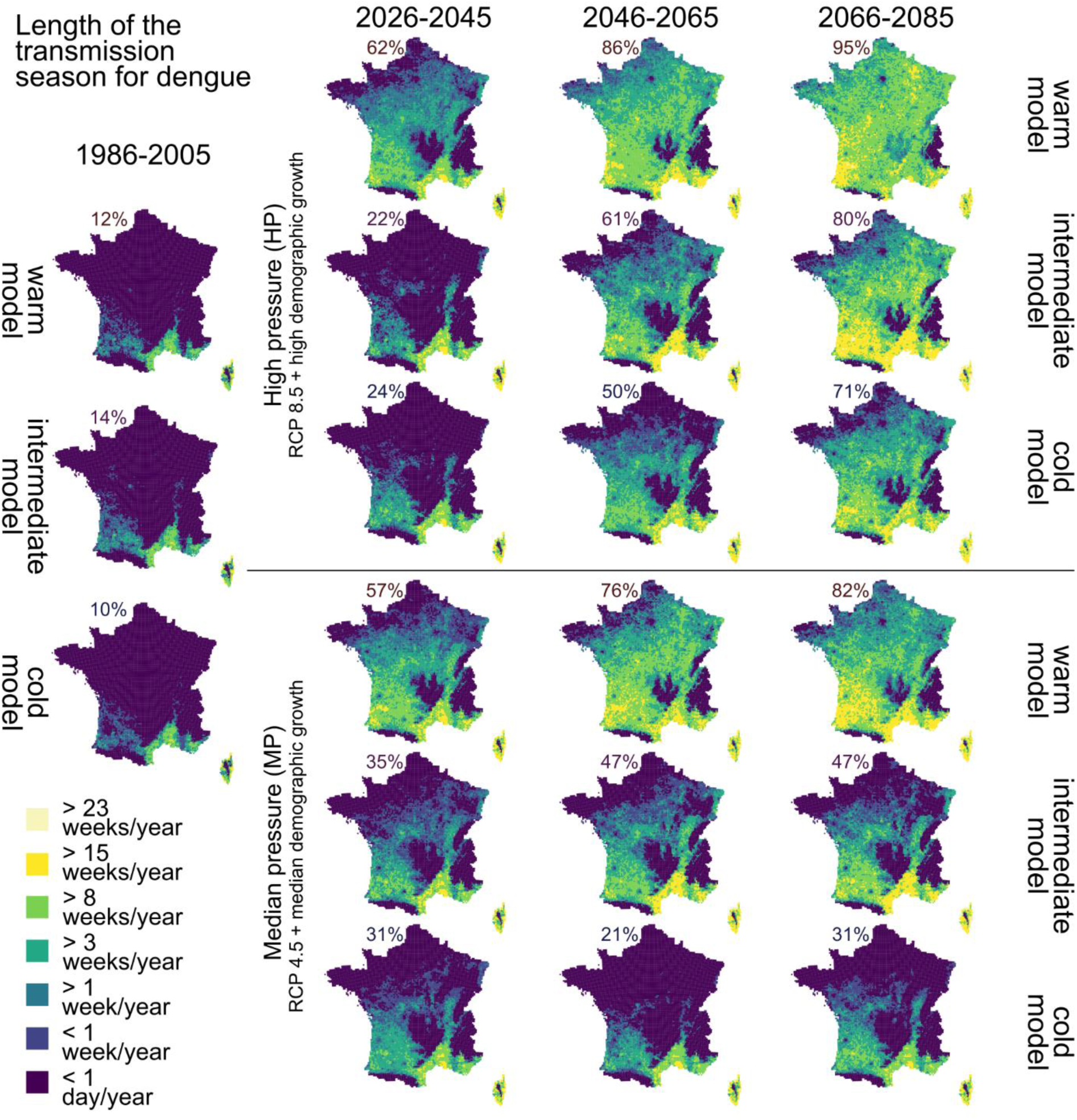
Projection of the duration of the transmission season of dengue (LTS: the number of days/weeks per year with R_0_>7). % of territory with LTS>1 week.

### Projected change in the epidemic size of dengue outbreaks

Simulated secondary cases range from 0, corresponding to a most (87%; 78-85%) of the French territory in 1986-2006, to around 60 (Figure 6). This indicator was estimated by assuming an introduction at peak transmission conditions; in France, this date, averaged for each model-scenario-time period, ranges between the end of June and the end of August (Figure SI2). Consistently with the definition of R_0_, we assumed a threshold of one secondary case to indicate regions at risk of dengue transmission. Under the HP scenario, the area at risk of dengue transmission depends on the climatic model in the short term, with a median value of 32 (31-73%). This uncertainty progressively decreases toward the end of the century, when areas at risk include a large part of France (88%; 82-96%). The simulated number of secondary cases increases as well. In Montpellier, the average number of secondary cases increases from 2.4 (1.9-3.6) in 1986-2006 to 10.0 (9.9-15.2) in 2066-2085. Such an increase is more evident in the neighboring rural settlement of Montarnaud where the number of secondary cases increases from 12.3 (6.6-21.0) to 31.6 (13.3-59.8). For these southern locations, secondary cases are higher in the intermediate model rather than the warm one. In Paris, the maximum number of cases increases from 0 (0-0) to 2.6 (1.7-13.4). Dengue-free regions still include mountain ranges and inner northern areas ranging from Brittany to Alsace. Under the MP scenario, the region at risk include 47% (40%-69%) of the territory in the short term, with a slight increase simulated at the end of the century (55%) and larger uncertainties (36-90%). The simulated number of secondary cases of dengue in 2066-2085 range from 9.0 (8.2-17.9) in Montpellier, to 46.2 (26.7-46.6) in Montarnaud, to 0.3 (0-3.6) in Paris.

**Figure 6.**
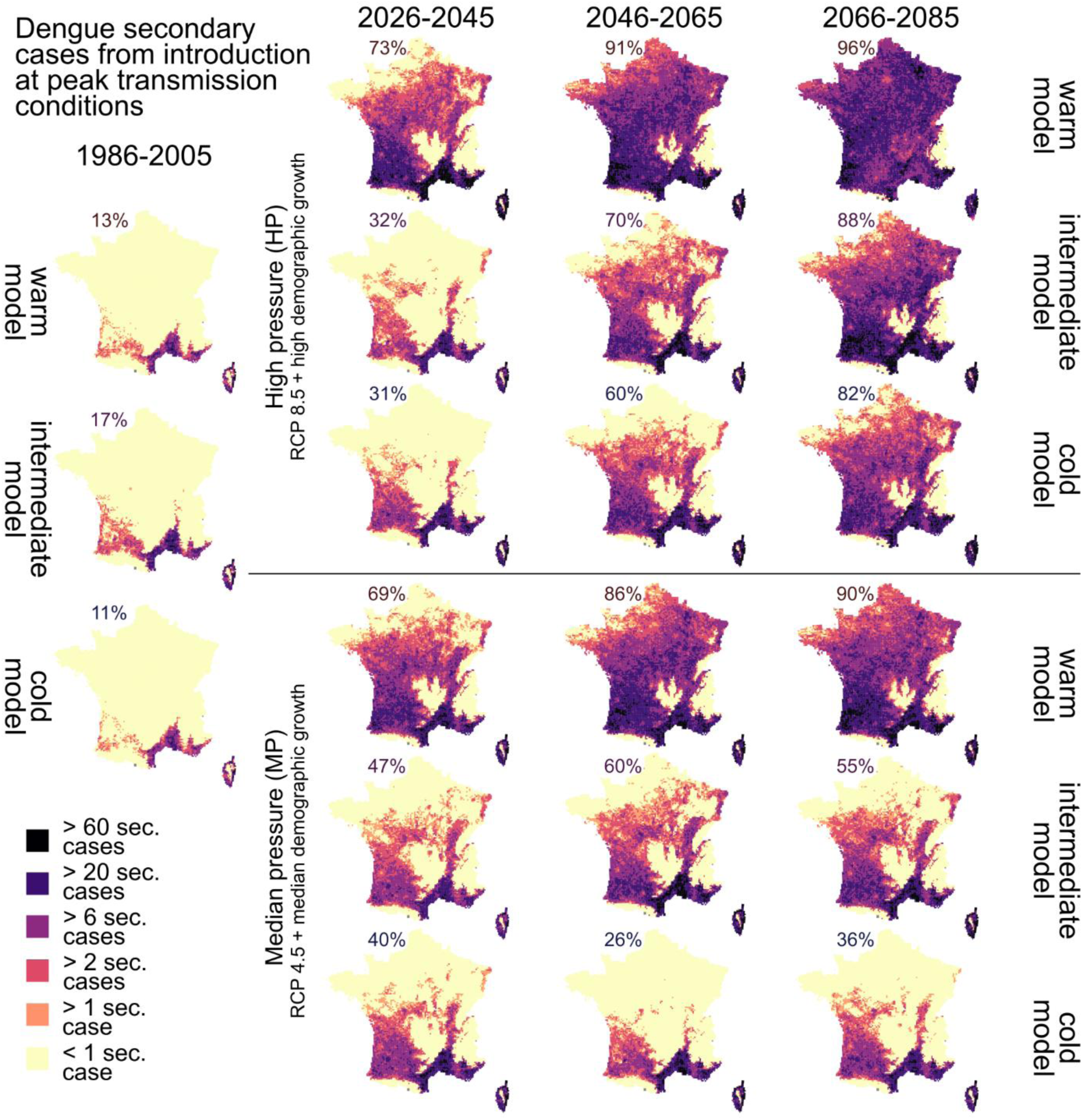
Projected number of secondary cases due to an uncontrolled introduction of a case of dengue one week before the date corresponding to the highest transmission risk.

To provide further insights on the dengue transmission period, we simulated the introduction of a dengue case at the beginning of each month, for each scenario, time period and climate model (Fig.s SI3 – SI29). In the HP scenario, the dengue transmission season of dengue might start as early as April along the Mediterranean coast in future.

### Projected seasonality of mosquito dynamics

Highly suitable sites (E_0_>1) tend to be associated with higher summer temperatures (which sustain rapid gonotrophic cycles, lower mortality, higher fertility) and higher winter temperatures (which increase the chances of survival of diapausing and non-diapausing eggs; Figure 7a,b). Under the HP scenario, the warmest sites such as Nice, are projected to host a stable homodynamic *Ae. albopictus* population (A_0_>1) in 2066-85. Such homodynamic population could encompass 0.6% of the French territory (0.2-0.7; Figure SI30) by the end of the century. Absence of diapause is more sporadic under the MP scenario (0-0.1%). Temperatures being similar, densely populated places (human densities ranging between 2500 and 5000 inhab/km²), with larger carrying capacity, tend to be more suitable to host such homodynamic populations than surrounding rural areas (human densities between 50 and 100 inhab/km²).

**Figure 7.**
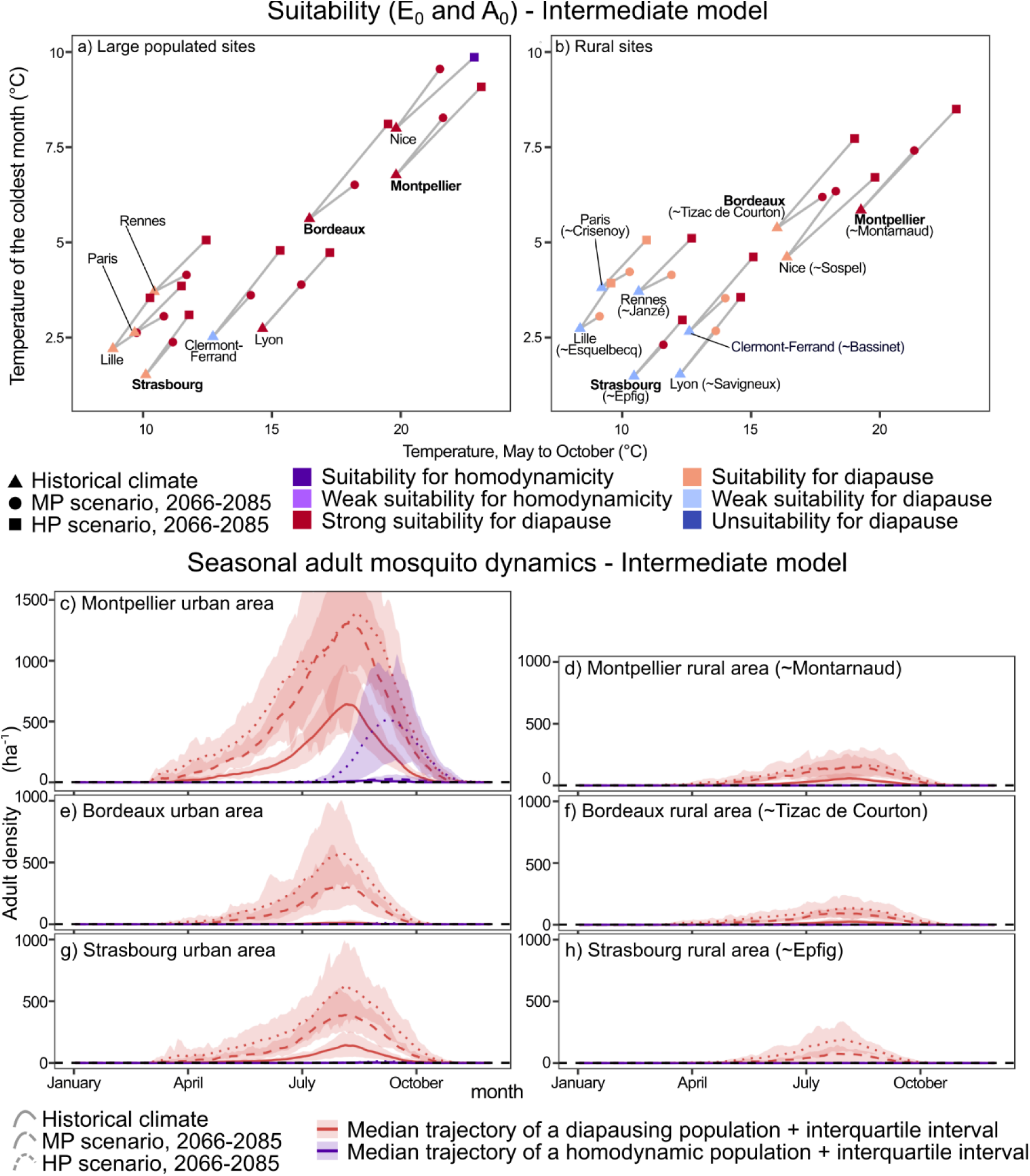
Projected change in suitability of the establishment of diapausing and homodynamic Ae. albopictus mosquitoes in urban (human density ranging between 2500 and 5000 inhab./km²) and rural (human density ranging between 50 and 100 inhab./km²) areas and their seasonal dynamics depending on the time period and scenario. Suitability for homodynamicity: A_0_ and E_0_>1; weak suitability for homodynamicity: A_0_>0.1 and E_0_>1; strong suitability for diapause: A_0_<0.1 and E_0_>10; suitability for diapause: A_0_<0.1 and E_0_>1; weak suitability for diapause: A_0_<0.1 and E_0_>0.1. Unsuitability for diapause: A_0_ and E_0_<0.1.

The difference in carrying capacity between urban and rural areas affects the seasonal dynamics of adult mosquitoes: large populated places, such as Montpellier, Bordeaux and Strasbourg host systematically more abundant populations (in the order of 10^2-3^ adult mosquitoes/ha; see Figure 7c,e,g) than their rural counterparts (in the order of 10^1-2^; Figure 7d,f,h) during the activity period. Climate change will alter such seasonality. In the Mediterranean region of Montpellier, where 1986-2005 baseline climate conditions were already favorable for the persistence of a diapausing population (solid red line in Figure 7c,d), a warming climate is expected to both increase the abundance of mosquitoes and lengthen their activity season by at least one month (from 241 days with more than 1 adult/ha to 274-282). However, we show a population decrease during summer in the warmest scenario (dotted red line in Figure 7c,d; such decrease is more evident in the warm and cold model, Figure SI31c,d, SI33c,d). The sporadic overwintering of a homodynamic *Ae. albopictus* strain might be sustained in urban centers and has an impact on mosquito presence at the end of the activity season (dotted purple line in Figure 7c). In the Bordeaux area, on the Atlantic coast, climate change will intensify and lengthen the activity season of adult mosquitoes by almost two months (from 211 days to 267-269 in Bordeaux; Figure 7e,f); in the warm model, both declines in summer adult populations, and sporadic homodynamic activity, are simulated (Figure SI31). Changing climate conditions have caused the presence of a diapausing strain to settle in Strasbourg (from 0 to 194-226 days; Figure 7g,h; details on specific sites in Tab. SI4).

### Projected seasonality in dengue transmission risk

High temperatures from spring to autumn increase the duration of the transmission season for dengue (LTS; Figure 8a,b). However, densely populated places (Figure 8a,c,e,g) which experience R_0_ values no greater than 2, have shorter and less intense transmission seasons compared to low population density locations (Figure 8b,d,f,h), where R_0_ can exceed 10 under high warming scenarios.

**Figure 8.**
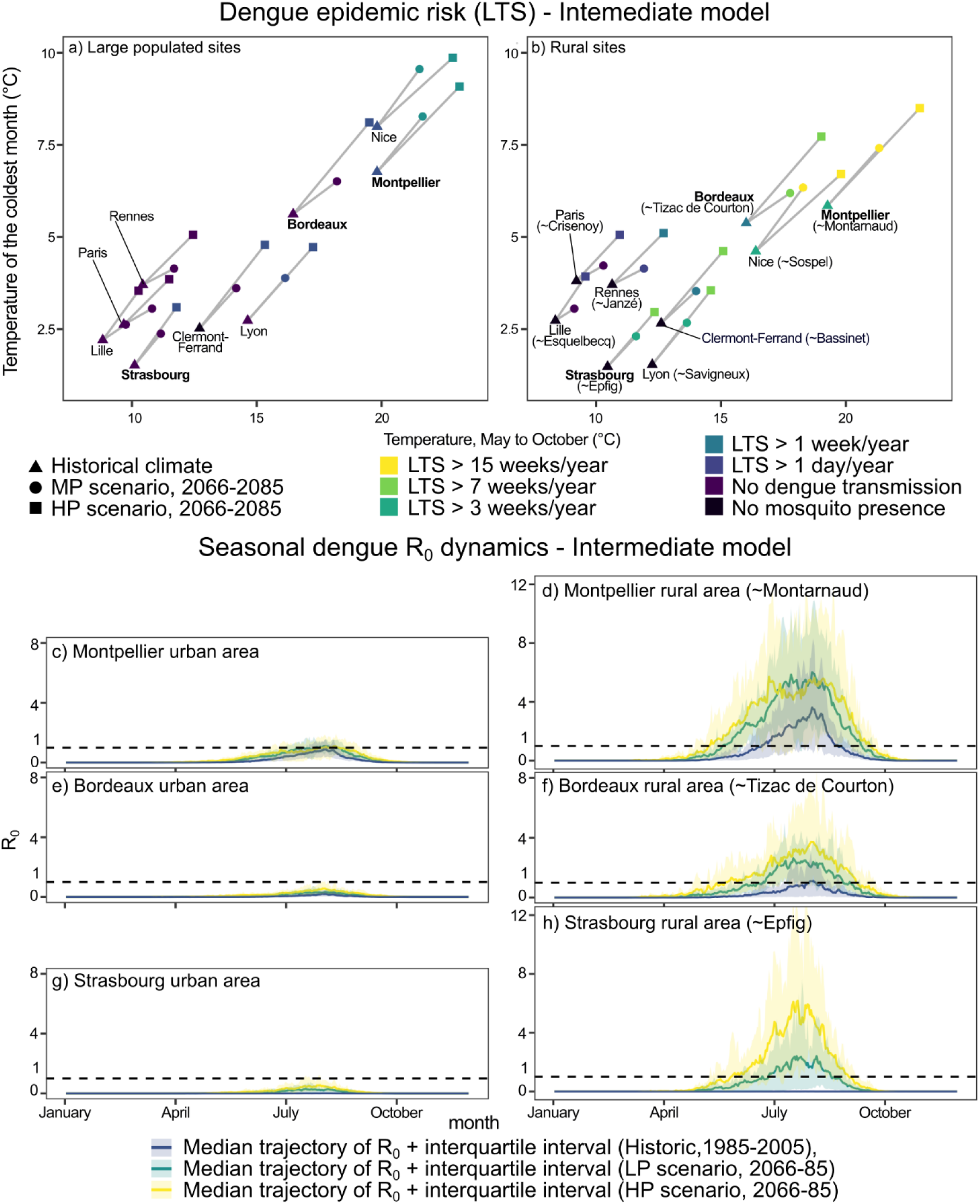
Projected change in the duration of the dengue transmission season in urban and rural areas and its seasonal dynamics depending on the selected time period and scenario.

In urban Mediterranean sites, climatic conditions were already theoretically compatible with sporadic dengue transmission in 1985-2005, climate change further causes a lengthening of the transmission season (from LTS = 0 to 14-17 in Montpellier under future projections, from 59 to 119-131 in Montarnaud; Figure 8b,c). In rural Mediterranean areas, a simulated decrease in R_0_ occurs during the warmest summer days and is a consequence of the contraction of the adult population. In the warm model, the simulated adult population decrease leads to a R_0_ value for dengue below 1, with a longer LTS under the MP scenario compared to the HP (118 against 63 in Montarnaud, Figure SI32). In the cold model R_0_ only reaches a plateau, with no clear simulated decrease (Figure SI34).

In both Atlantic and continental areas like Bordeaux and Strasbourg, the effect of climate change will both increase the intensity and the duration of the transmission season, with a stronger effect on rural sites, while R_0_ rarely exceeds the threshold of 1 in urban centers (Figure 8e,f,g,h). Contrary to Bordeaux, simulated dengue transmission was impossible (R_0_ = 0) in the Strasbourg area in 1986-2005 (with a colder climate). Nonetheless, under the HP scenario, maximum R_0_ values in Strasbourg exceed those of Bordeaux in 2066-85 (4.9 against 3.2). However, this HP scenario has no impact on the duration of the transmission season, which remain longer in the Bordeaux area (LTS in Tizac de Courton, Bordeaux area: from 0 to 64-99; in Epfig, Strasbourg area: from 0 to 45-90; details on specific sites in Tab. SI4).

### Sensitivity to the pressure scenarios

To reduce the number of scenarios, we selected two out of the 44 possible combinations of the climatic and demographic scenarios. This choice does not allow us to disentangle all potential combined effects of climatic and population changes. We therefore assessed the relative weight of both factors by fixing one and letting the second evolve. When exploring climate effects, population was set according to the 1986-2005 scenario, and reciprocally when exploring human density effect, the ‘historical’ climatic scenario was used, assessing the relative change in suitability for the establishment and for dengue transmission.

This sensitivity analysis revealed that simulated changes in entomological and epidemiological variables mostly depend on climate change scenarios, which determine more than 99% of the variations. The largest contribution of population changes correspond to a 0.7% increase in surface suitable for the establishment of the vector and a 0.5% increase in surface suitable for dengue transmission at the end of century under the HP scenario (Figure SI35-36).

## Discussion

### *Ae. albopictus* will be ubiquitous in France by the half of the 21^st^ century

As most scenarios and models suggest, almost the entire French population will likely live in areas inhabited by *Ae. albopictus* by mid-century. Only mountain ranges, Alps and Pyrenees *in primis*, will remain unsuitable for this species, while some low-populated areas in the North, such as the Ardenne-Champagne region, the Picard plateau, the Armorican massif and the Norman hills may experience sporadic presence of the vector. This result is consistent with other modelling studies, conducted at larger spatial scale. Pardo-Araujo et al. (2024) suggested that France will be suitable to host *Ae. albopictus* activity for at least four months a year in the second half of the century, except for Alpine areas. The same study suggests that, *Ae. albopictus* may be active for at least three months a year in the south of the British isles (consistent with findings by Metelmann et al., 2019) and could spread to the southern Baltic regions. Kraemer et al. (2019) showed that *Ae. albopictus* might also spread to the Baltic area in the RCP6.0 scenario. In addition, Atlantic and continental Europe (including northern France) could become suitable for the establishment of this species, although with a low confidence in this projection. On the other hand, Brass et al. (2024) suggested that Atlantic and continental Europe has nowadays the potential to host adult population with density of about 100 adults/ha during summer.

The ubiquitous establishment of *Ae. albopictus* has natural consequences on the expansion of the area at risk of dengue transmission, with however large uncertainties related to the emission scenario and the selected climate model. In this study, we estimated two different measures of epidemiological risk: (i) the length of the transmission season (LTS), widely used in the literature to compare the impacts of scenarios, and (ii) the secondary cases, primarily used to study the dynamics of recent arbovirus outbreaks in France and Italy (Tegar et al., 2025, 2025; White et al., 2025). Ryan et al. (2019) found that, by 2050, dengue could potentially be transmitted everywhere in France in a low emission scenario (RCP2.6), with little variation in the following decades. LTS could increase up to 3-4 months in the high emission scenario (RCP8.5) at the end of the century; this finding aligns well with our estimates. Note that the same study suggests that dengue could also be transmitted in lowland areas of the Scandinavian region in future. More cautiously, Colón-González et al. (2021), in a comparative study integrating different epidemiological models and climate scenarios, showed that the increase in simulated LTS in France will not exceed one month with respect to the 1970-00 historical baseline. Both studies are based on global climate models that have a lower spatial resolution compared to our regional climate estimates.

### Increased risk for rural areas

Although simulated mosquito abundance is usually low in rural areas, low-density human settlements experience and will experience the stronger effects of climate change on dengue transmission risk. In rural areas, we simulate both longer transmission seasons and an increased number of secondary cases compared to urban centers. Even if counterintuitive, such finding matches the distribution of recently reported arboviral outbreaks in Europe, which very rarely occur in densely-inhabited areas to date (Sacco et al., 2024; Santé Publique France, 2024, 2025). In the model, this is the result of two relationships, the first one relating human density and larval carrying capacity, and the second linking human density and the biting rate of mosquitoes. In our adjusted model, the carrying capacity grows less than linearly with human density. This directly affects the simulated abundance of adult mosquitoes. Instead, human density is inversely proportional to the biting rate; human hosts will receive less bites *per capita* in densely populated places. R_0_, which depends on the ratio between vectors (estimated with adult mosquito density) and hosts (estimated with human density; Smith et al., 2012), decreases with human density other factors being equal. Projected demographic changes, which predict a maximum increase of 10 million people in France in the next 50 years, will not affect the repartition of population density between populated and rural areas, with the former expected to absorb the greater proportion of this increase. Consistently, our sensitivity analysis suggests that climate – and not demographic – trends will dictate the increase of epidemic dengue risk in France.

Nonetheless, large cities will not be immune from the risk of arboviral transmission. Our results need to be contextualized with respect to the spatial resolution of the data. In fact, scarcely populated neighborhood exists in large urban centers, but they are not captured by the spatial resolution of climate projections. These areas, hosting urban parks, graveyards, and urban sprawl, have all characteristics needed to host *Ae. albopictus* mosquitoes: presence of irrigation water, suitable humidity conditions, presence of human hosts, low vegetation cover that provides sugary meals and an adequate microclimate to rest (Bartholomée et al., 2025; Clauzon et al., 2025; Mercat et al., 2025).

### Expected changes in *Ae. albopictus* seasonality

In most temperate areas of Europe, where *Ae. albopictus* has established, population dynamics follow a typical pattern, which consists in a population increase in spring, one or more summer peaks in activity corresponding to favorable weather conditions, and a slow population decrease in autumn (Taconet et al., 2025). It is extremely rare not to observe *Ae. albopictus* mosquitoes in summer, and winter presence of mosquitoes is anecdotal out of indoor context (such as in houses or greenhouses). Depending on the climatic scenario, we might witness significant changes in seasonality. In the Mediterranean region, future increase in winter temperatures may favor the selection of population active all-year-round. In Nice, in which the average temperature in January will be around 10 °C in fifty years in the warmest scenario, climate conditions may favor the viability of a small non-diapausing population, whose dynamics are characterized by a secondary abundance peak delayed in autumn. This feature is already observed in tropical areas, and, according to Del Lesto et al. (2022), may also occur in southern Europe within a few decades. These authors suggested a lower limit of 10 °C for population to overwinter.

Another expected abrupt change is the potential collapse of mosquito population during summer, reducing also dengue transmission for short and extremely warm periods. We successfully reproduced this phenomenon in many areas of southern France, especially in the warmest scenario simulations. *Ae. albopictus* thermal tolerance has an upper limit, experimentally determined to be around 36-40 °C (Souza & Weaver, 2024). Based on these lab-based experiments, other authors have found a reduction in the simulated abundance of mosquito population and dengue transmission risk, in warm climates. For example, Ryan et al. (2019) and Colón-González et al. (2021) project a decrease in the duration of the dengue transmission season in tropical Africa in the next decades. In southern Europe, Garrido Zornoza et al. (2024) showed that heat waves have a detrimental effect on *Ae. albopictus* populations in already warm locations. Despite the growing evidence of the negative impact of extreme heat events on *Aedes* mosquitoes *in vivo* (Alfaro et al., 2025), empirical evidence of this phenomenon is somehow limited *in natura*. So far, recent climate change has more probably favored, and not inhibited, the recent spread of dengue in tropical regions (Childs et al., 2025). In southern Europe, heat waves have contributed to dengue outbreaks (Virgillito et al., 2025); some authors suggest that, as it adapted to cool climates, *Ae. albopictus* might also adapt to even warmer conditions (Couper et al., 2025).

It should be noted that, in our simulations, simulated mosquito population collapses in the HP scenario do not represent a decline relative to the historical period, but they represent a relative decrease compared to the MP scenario.

### Caveat and further directions

In this study, we modelled the impact of climate and demographic changes on *Ae. albopictus* population and dengue transmission risk in France. While we have considered uncertainties related to different emission and population scenarios and climate hypothesis, future boundary conditions influencing dengue spread may still differ from our projections.

*Aedes aegypti* is a more efficient vector to transmit dengue than *Ae. albopictus* (Kristan et al., 2025). This species is widespread in tropical and semi-tropical countries; it is now present in southern Europe (Cyprus) and peripherical areas, such as Madeira (Portugal), the eastern coasts of the Black Sea in Russia, and the northeastern coasts of Türkiye (ECDC, 2025). During the first part of the 20^th^ century, this species was also widespread over the Mediterranean coasts, before being eradicated as a side effect of DDT spraying to control malaria post WWII (Holstein, 1967; Wint et al., 2022). Empirical evidence indicates that the thermal tolerance of *Ae. aegypti* has shifted toward warmer temperatures compared to that of *Ae. albopictus* (Souza & Weaver, 2024). There is currently no consensus regarding the potential establishment of *Ae. aegypti* in Europe. Some authors (Pardo-Araujo et al., 2024; Ryan et al., 2019) suggested that future climatic conditions in continental Europe may become suitable for *Ae. aegypti*, while others propose that such colonization is already possible (Da Re et al., 2021), as already observed in the past. Conversely, Kraemer et al. (2019) excluded this possibility. In this study, we focused on *Ae. albopictus*, since it is already established in Europe and for which extensive local data is available.

*Ae. albopictus* is expected to keep evolving, as it has done over the last century. Here, we specifically examined the hypothesis that diapause may disappear entirely; however, it is more likely that this species will gradually adapt to local climate conditions, as observed in North America (Batz et al., 2020; Urbanski et al., 2012). With milder winter, mosquitoes may lower their critical photoperiod threshold, thereby shortening their diapause period: longitudinal field entomological studies are urgently needed to support this hypothesis.

In this study, we adopted a common epidemiological modeling framework by considering horizontal transmission only. However, vertical transmission of arboviruses has been observed in endemic countries for both *Ae. albopictus* and *Ae*. *aegypti* (Ferreira-De-Lima & Lima-Camara, 2018). More research is needed to characterize the risk of vertical transmission in Europe, which may allow the interannual persistence of arboviruses in the environment without the need to reintroduce infected cases returning from endemic countries.

This study builds upon a previous analysis that focused on the impact of recent climate change on *Ae. albopictus* and dengue transmission risk in Europe (Radici et al., 2025) to provide higher spatial resolution estimates for France (to ∼81 to ∼64 km²). However, even if we used calibrated regional climate model projections, they still do not account for microclimatic and landscape effects at the sub-kilometric scale. Such microclimatic conditions will ultimately determine local interactions among vectors, hosts and the environment (Herbreteau et al., 2025).

### Suggestions to face the increasing risk of dengue transmission

*Aedes albopictus,* and *Aedes-*borne viruses, will become an increasingly public health and nuisance threat in the upcoming years in France and Europe, particularly due to increasingly favorable climate change conditions. Policymakers, vector control operators, and public health officers already have solutions to prevent and control this threat (integrated vector management, vaccines, antiviral drugs). This study intended to provide multiple scenarios at high spatial resolution in order to anticipate future risk and improve these strategies.

An active surveillance of *Ae. albopictus* should be conducted in areas predicted to be suitable. In already colonized areas, one should monitor the vector adaptation to heat, its winter activity, and its potential of vertical transmission. Currently, surveillance for arboviruses is dictated by a seasonal calendar; in the future, a predictive model based on weather observations and forecasts will be beneficial to adjust this calendar and guide vector management. With rising temperatures, longer activity seasons and the gradual adaptation of existing populations there will likely be increasingly frequent and widespread epidemics of arboviruses that will require novel and timely control interventions. Such arboviruses epidemics will demand the mobilization of more human resources and the prioritization of interventions. In this sense, it is useful to draw inspiration from integrated control measures undertaken in tropical countries where the virus is endemic, which also include extensive prevention measures.

The future of *Ae. albopictus* and dengue transmission risk in temperate regions will ultimately depend on climate change, which in turn depends on anthropogenic greenhouse gas emissions. While climate change had no role in bringing the mosquito into Europe, climate change paved the way for its colonization. The future of the so-called tropical diseases ultimately depends on the climate policies of today.

## Acknowledgements

The authors acknowledge the support of RIVOC funded by the Région Occitanie (France) for the VectoClim project. In addition, A.R. acknowledges precious scientific discussions with Claire Garros, Frédéric Simard, Emmanuel Pont, Ying Long, Paul Taconet, Benjamin Le Roy, Antoine Mignotte, Grégory Lambert, Guillaume Lacour, Daniele Da Re, Hugo Tessier. A.R. acknowledges also the use of the i-Trop HPC and would like to thank the team of the i-Trop platform for their technical support.

Authors declare no competing interests.

