## Supplementary Information for "*Aedes albopictus* and dengue transmission risk in France over the 21^st^ century"

### Metelmann et al. model parameters

In the modelling framework established by Metelmann et al., (2019) both the carrying capability $K_{it}$ and the hatching $h_{it}$ of a site $i$ at time $t$ depend on host density $H_{it}$ and rainfall $r_{it}$ :

$$K_{xt} = \lambda \frac{1-\alpha_{evap}}{1-\alpha_{evap}{}^{t}} \sum_{\tau=1}^{t} \alpha_{evap}{}^{(t-\tau)} \left( \alpha_{rain} r_{x\tau} + \alpha_{dens} H_{x\tau} \right)$$

$$h_{xt} = (1-\varepsilon_{rat}) \frac{(1+\varepsilon_0) e^{-\varepsilon_{var}(r_{xt}-\varepsilon_{opt})^2}}{e^{-\varepsilon_{var}(r_{xt}-\varepsilon_{opt})^2} + \varepsilon_0} + \varepsilon_{rat} \frac{\varepsilon_{dens}}{\varepsilon_{dens} + e^{-\varepsilon_{fac} H_{xt}}}$$

Parameter values of the abovementioned equations and those presented in the main text with the corresponding source are provided below:

*Table SI1 - Model parameters*

| Parameter | Variable name (units) | Value/formula | Source |
|---|---|---|---|
| $CTT_S$ | critical temperature over one week in spring (°C ) | $11.0$ | (Metelmann et al., 2019) |
| $CPP_S$ | critical photoperiod in spring (hours) | $11.25$ | (Metelmann et al., 2019) |
| $\sigma(T,P)$ | spring hatching rate (day$^{-1}$) | $\begin{cases} 0 \text{ if } T_7 < CTT_S \text{ or } P < CPP_S \\ 0.1 \text{ otherwise} \end{cases}$ | (Metelmann et al., 2019) |
| $CPP_A(L)$ | critical photoperiod in autumn (hours) | $10.058 + 0.08965L$ | (Metelmann et al., 2019) |
| $\omega(P)$ | fraction of eggs going into diapause | $\begin{cases} 0 \text{ if } P < CPP_A \text{ or day} < 183 \\ 0.5 \text{ otherwise} \end{cases}$ | (Metelmann et al., 2019) |
| $\delta_E$ | normal egg development rate (day$^{-1}$) | $1/7.1$ | (Metelmann et al., 2019) |
| $\delta_J(T)$ | Juvenile development rate (day$^{-1}$) | $1/(83.85 - 4.89\,T + 0.08\,T^2)$ | (Metelmann et al., 2019) |
| $\delta_I(T)$ | first pre-blood meal rate (day$^{-1}$) | $1/(50.1 - 3.574\,T + 0.069\,T^2)$ | (Metelmann et al., 2019) |
| $\mu_E$ | egg mortality rate (day$^{-1}$) | $-\ln\left(0.955\, e^{-0.5\left(T-18.8/21.53\right)^6}\right)$ | (Metelmann et al., 2019) |
| $\mu_J$ | juvenile mortality rate (day$^{-1}$) | $-\ln\left(0.977\, e^{-0.5\left(T-21.8/16.6\right)^6}\right)$ | (Metelmann et al., 2019) |
| $\mu_A(\bar{T})$ | adult mortality rate (day$^{-1}$) | $-\ln\left(0.677\, e^{-0.5\left(\bar{T}-20.29/13.2\right)^6} 0.069\, \bar{T}^{0.1}\right)$ | (Metelmann et al., 2019) |

| | | | |
|---|---|---|---|
| $\gamma(T_w)$ | survival probability of diapausing eggs (winter$^{-1}$) | $0.93\ e^{-0.5(T_w-11.68/15.67)^6}$ | (Metelmann et al., 2019) |
| $\beta(T)$ | egg laying rate (day$^{-1}$) | $\begin{cases} 33.2\ e^{-0.5(T-70.3/14.1)^2}(38.8-T)^{1.5} \text{ if } T \leq 38.8 \\ 0 \text{ otherwise} \end{cases}$ | (Metelmann et al., 2019) |
| $\lambda$ | capacity parameter (larvae days ha$^{-1}$) | $10^6$ | (Metelmann et al., 2019) |
| $\alpha_{evap}$ | Normalization parameter evaporation | 0.9 | (Metelmann et al., 2019) |
| $\alpha_{dens}$ | Normalization parameter of host density (km$^2$) | $10^{-5}$ | (Metelmann et al., 2019) |
| $\alpha_{rain}$ | Normalization parameter of rain (mm$^{-2}$) | $10^{-2}$ | (Metelmann et al., 2019) |
| $\varepsilon_{rat}$ | Proportion of hatching due to rain | 0.2 | (Metelmann et al., 2019) |
| $\varepsilon_0$ | Correcting additive parameter for rain-related hatching | 1.5 | (Metelmann et al., 2019) |
| $\varepsilon_{var}$ | Stretching parameter for rain-related hatching (mm$^{-2}$) | 0.05 | (Metelmann et al., 2019) |
| $\varepsilon_{opt}$ | Correcting additive parameter for human-induced hatching (mm) | 8 | (Metelmann et al., 2019) |
| $\varepsilon_{dens}$ | for rain-related hatching | $10^{-2}$ | (Metelmann et al., 2019) |
| $\varepsilon_{fac}$ | Stretching parameter for human-induced hatching (km$^2$ km$^{-2}$ | $10^{-2}$ | (Metelmann et al., 2019) |
| $a$ | Mosquito biting rate | $(0.0043T + 0.0943)/2$ | (Blagrove et al., 2020; Caminade et al., 2017; |

| | | | |
|---|---|---|---|
| | | | Zardini et al., 2024) |
| $\varphi$ | Human biting preference | $\begin{cases} 0.9 \text{ if human density} > 50 \text{ hab km}^{-2} \\ 0.5 \text{ otherwise} \end{cases}$ | Value of 0.5 by (Caminade et al., 2017) |
| $EIP\ (\nu^{-1})$ | Extrinsic (mosquito) incubation period | $1.03(4 + \mathrm{e}^{5.15\,-\,0.123\mathrm{T}})$ | (Caminade et al., 2017; Metelmann et al., 2021) |
| $r$ | Human recovery rate for dengue | $1/7$ | (Caminade et al., 2017) |
| $\mathrm{B}_{v,h}$ | Probability of transmission of the disease from an infected vector to a susceptible host | $0.5$ | (Blagrove et al., 2020) |
| $\mathrm{B}_{h,v}$ | Probability of transmission of the disease from an infected host to a susceptible vector | $0.31$ | (Metelmann et al., 2021) |
| $\delta_M$ | Maximal number of hosts bitten by a mosquito per ha | $4.5$ | (Metelmann et al., 2021) |

The definition of the weather and environmental variables is provided in Table SI2.

*Table SI2 – Weather and environmental variables and their definitions*

| **Weather and environmental variable** | **Meaning** |
|---|---|
| $T$ | Instantaneous temperature (°C), approximated using a sinusoidal function between on daily minimum and maximum (Metelmann et al., 2019) |
| $\bar{T}$ | Mean daily temperature (°C) |
| $T_7$ | Mean temperature of the last week (°C) |
| $T_w$ | Lowest winter (December-January-February) temperature (°C) |
| $R$ | Daily rainfall (mm) |
| $H$ | Human host density (hab/km$^2$); the original format of the GPWv4 data (Doxsey-Whitfield et al., 2015) is absolute count, but we converted this into density (people/km$^2$) to be consistent with the model input requirements |
| $P$ | Photoperiod (hours) |

| | |
|---|---|
| $L$ | Latitude (°) |

### Results

In the main text, we used observed epidemiological data to calibrate the value of $\varepsilon = 0.6$. To do so, we simulated the number of secondary cases by stopping the simulation at the date of the last known case. This is an approximated modelling implementation of the control by vector management officers. However, it provides an approximation of the order of magnitude for the number of secondary cases.

In table 3, we summarize the comparison between observed and simulated secondary cases for 6 locations in France . Furthermore, we computed the number of secondary cases if no control measures were implemented (i.e. letting the epidemic run until natural extinction).

| Site | Observed secondary cases | Simulated secondary cases | Simulated secondary cases (no control) | Absolute decrease due to control | Relative decrease due to control |
|---|---|---|---|---|---|
| Frejus | 15 | 24.3 | 59.7 | 35.4 | 59.3% |
| La Crau | 25 | 22.5 | 53.2 | 30.7 | 57.7% |
| S. Cecile les Vignes | 18 | 96.4 | 144.5 | 48.1 | 33.3% |
| Vallauris | 14 | 7.2 | 21.4 | 14.2 | 66.4% |
| Rognac | 5 | 51.5 | 158.1 | 106.6 | 67.4% |
| Aubagne | 9 | 1.6 | 17.5 | 15.9 | 90.9% |
| Tot | 86 | 203.5 | 454.4 | 250.9 | 55.2% |

*Table 3 - Comparison between observed and simulated secondary cases, with and without control interventions.*

#### Projected adult mosquito density between May and October

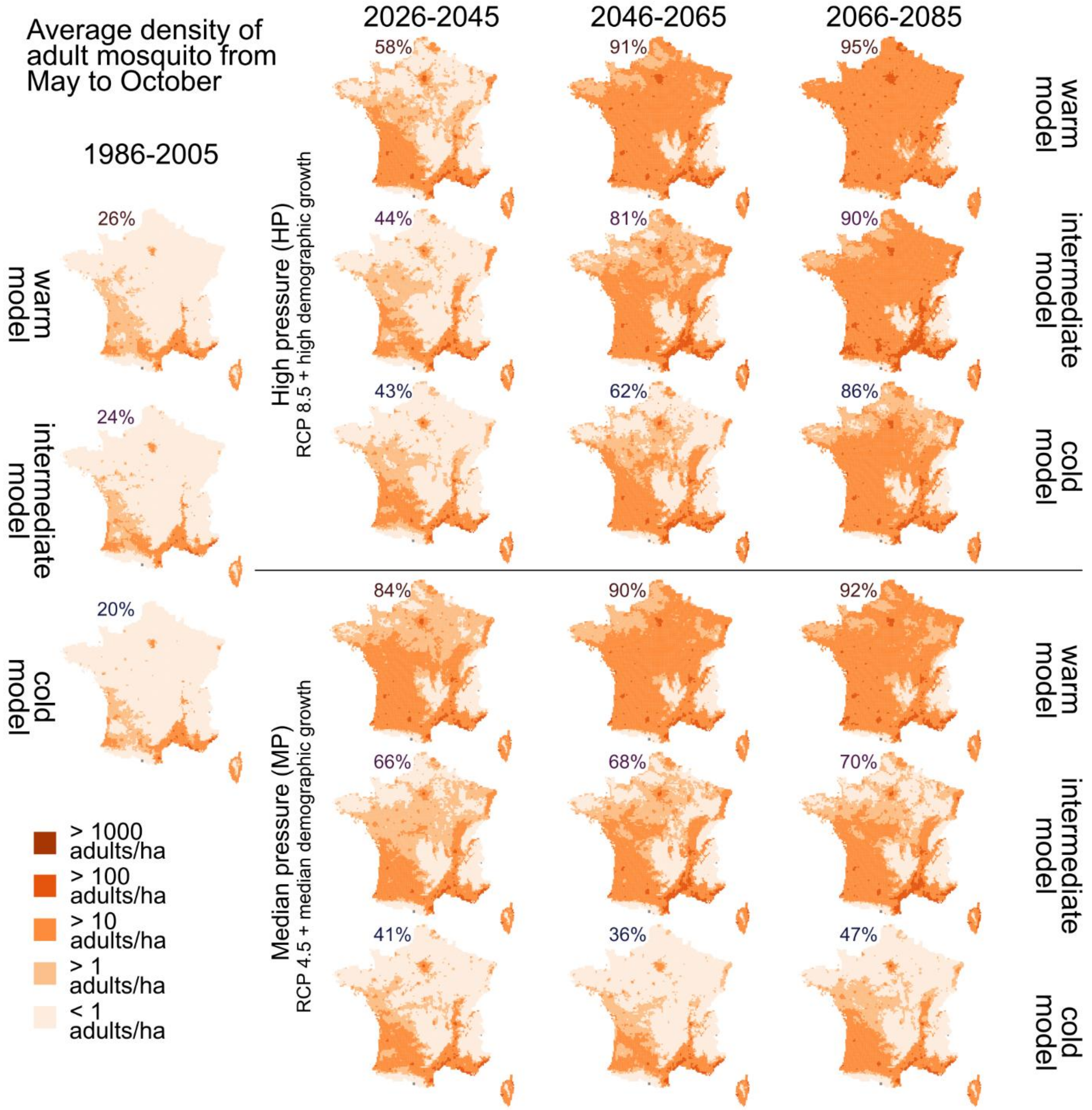


*Figure 1 - Projected adult density (adults/ha) for the May-Oct activity season for different scenarios and climate models.*

#### Projected change for the day of the year with the highest transmission risk

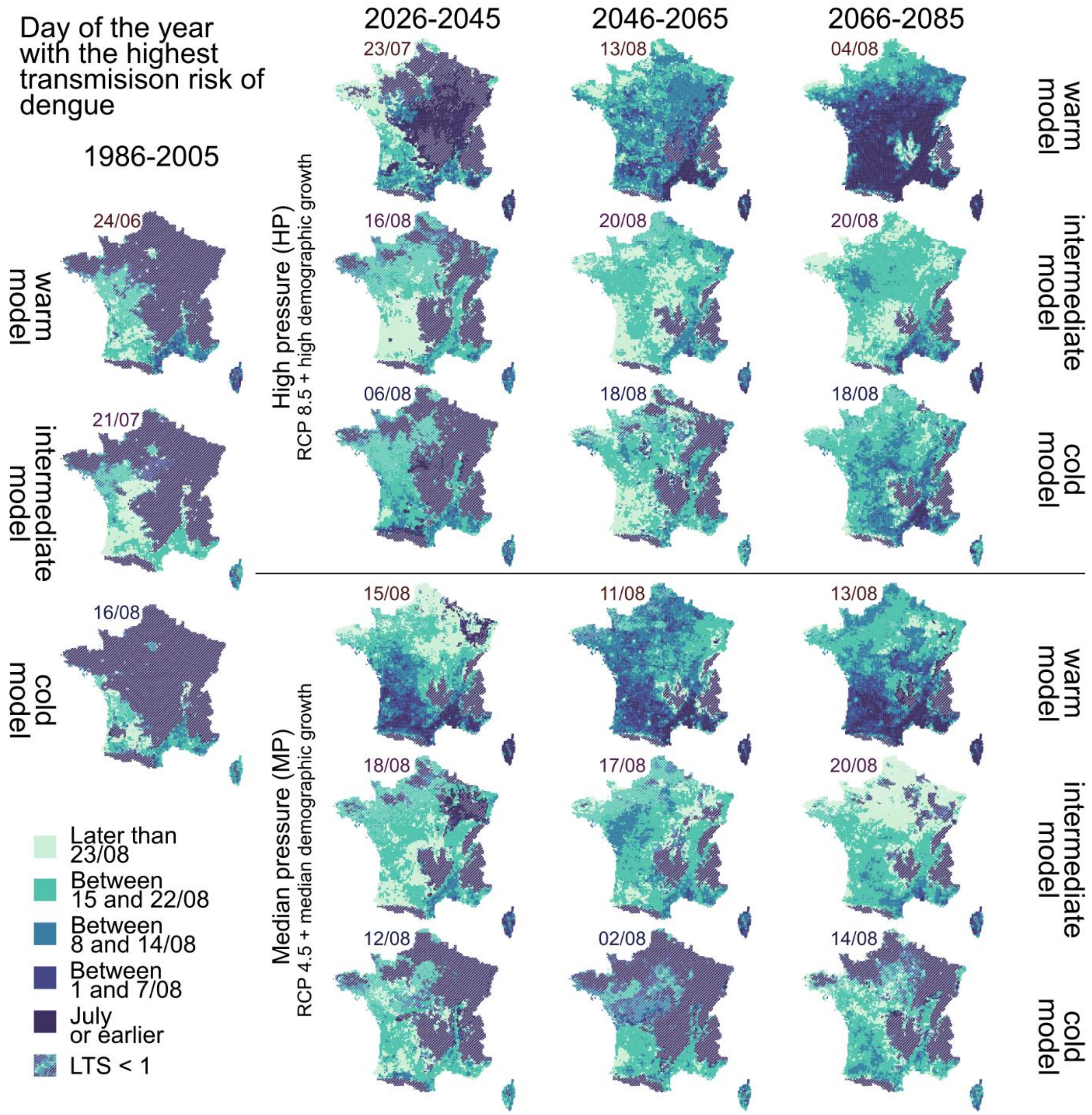


*Figure 2 - Day of the year with highest $R_0$ for dengue, for different scenarios and climate models.*

#### Projected size of a dengue outbreak by month of introduction (intermediate model)

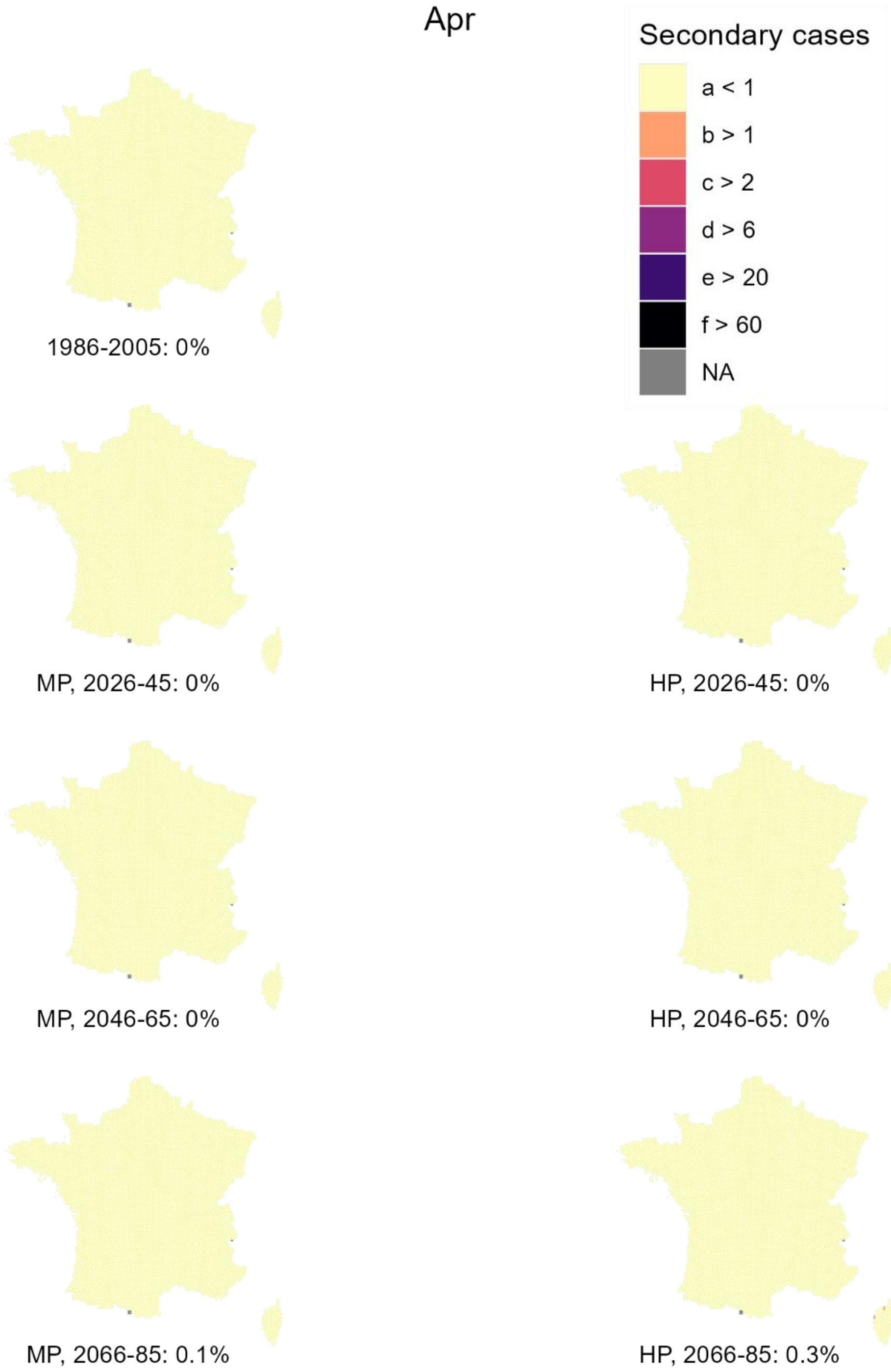


*Figure 3 - Projected number of secondary dengue cases generated in April by the introduction of a positive case on the first day of the month (Intermediate model) for each scenario and time horizon. In brackets, we show the fraction of territory where at least 1 secondary case occurs.*

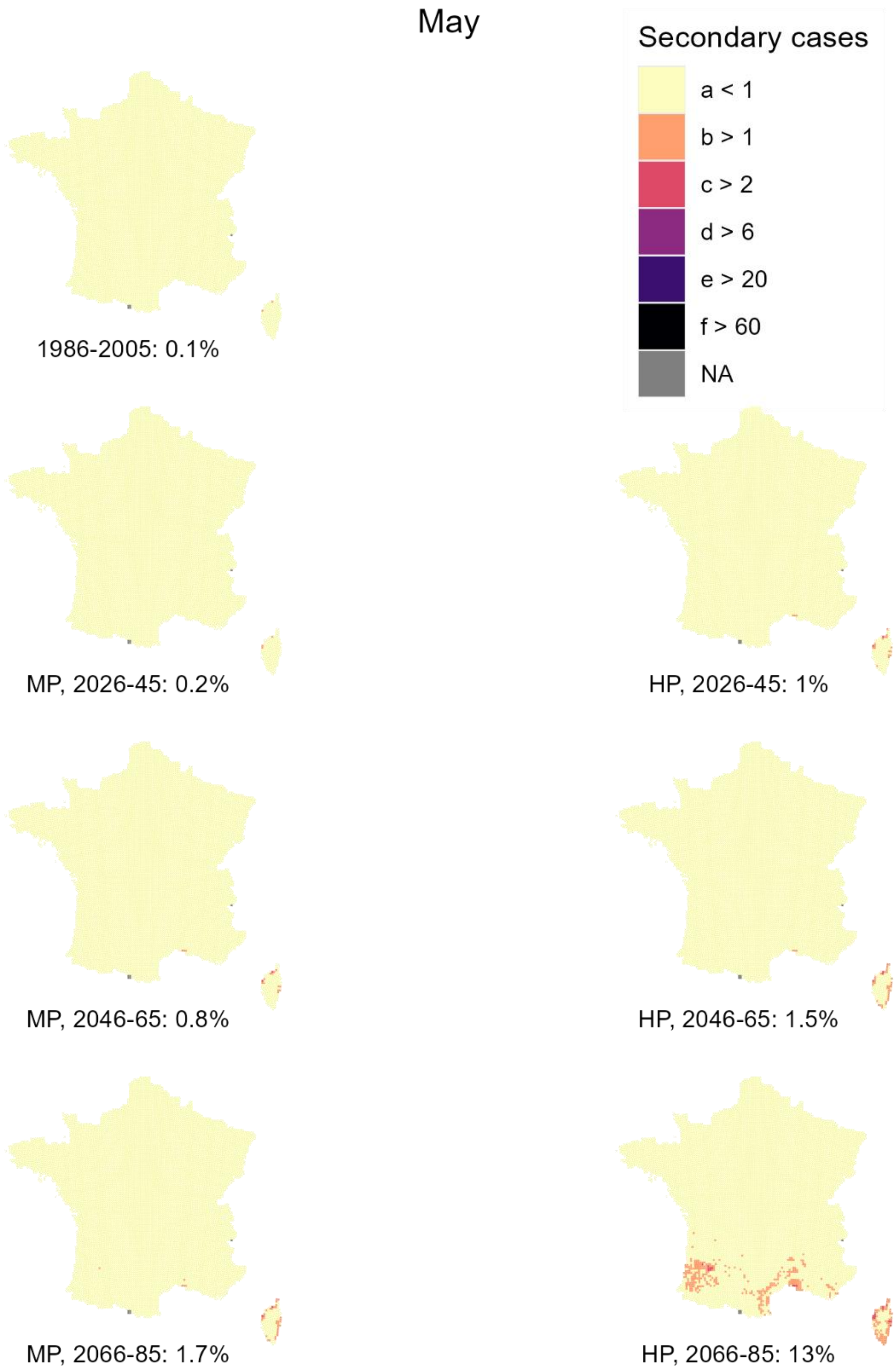


*Figure 4 - Projected number of secondary dengue cases generated in May by the introduction of a positive case on the first day of the month (Intermediate model) for each scenario and time horizon. In brackets, we show the fraction of territory where at least 1 secondary case occurs.*

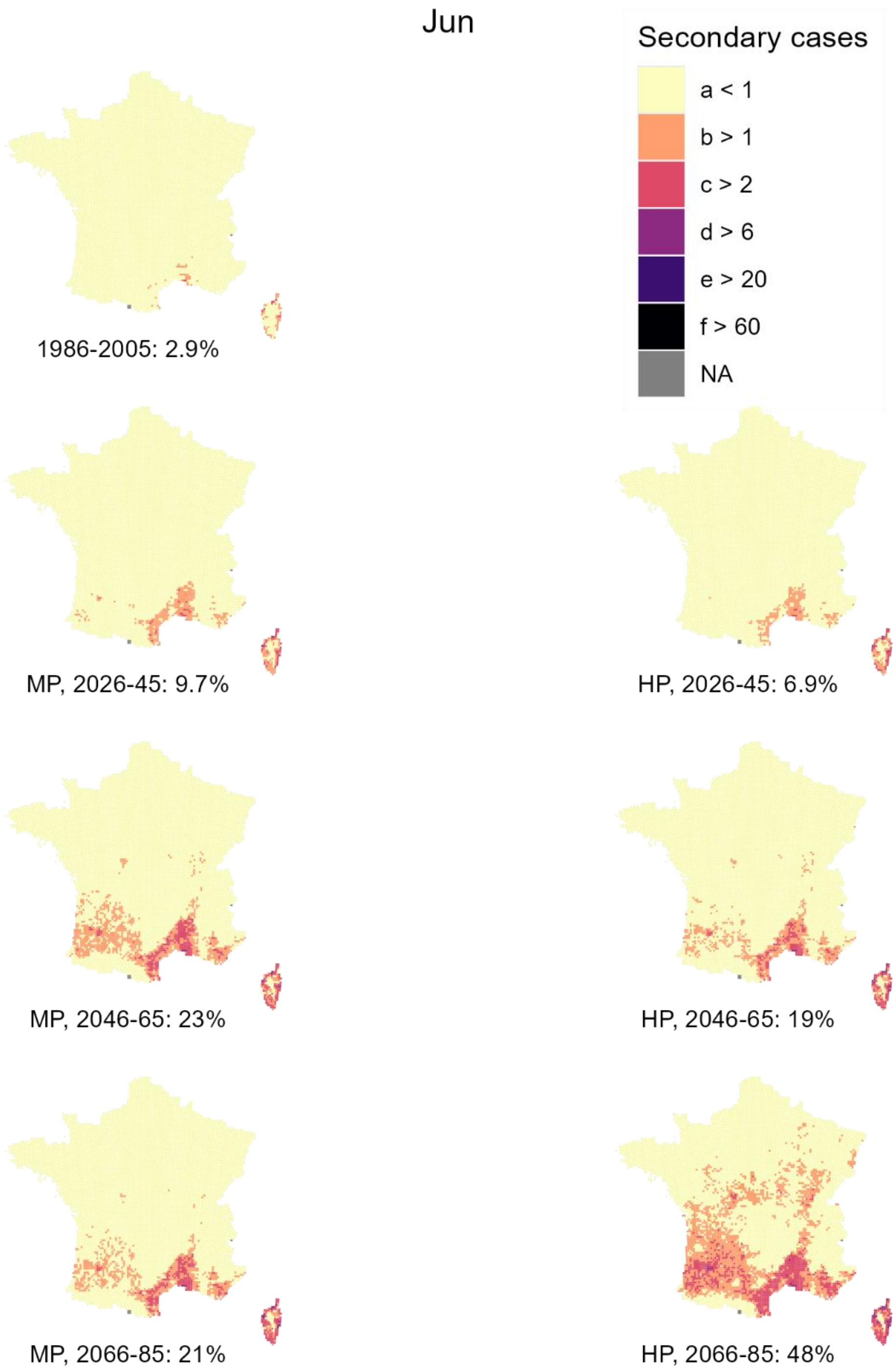


*Figure 5 - Projected number of secondary dengue cases generated in June by the introduction of a positive case on the first day of the month (Intermediate model) for each scenario and time horizon. In brackets, we show the fraction of territory where at least 1 secondary case occurs.*

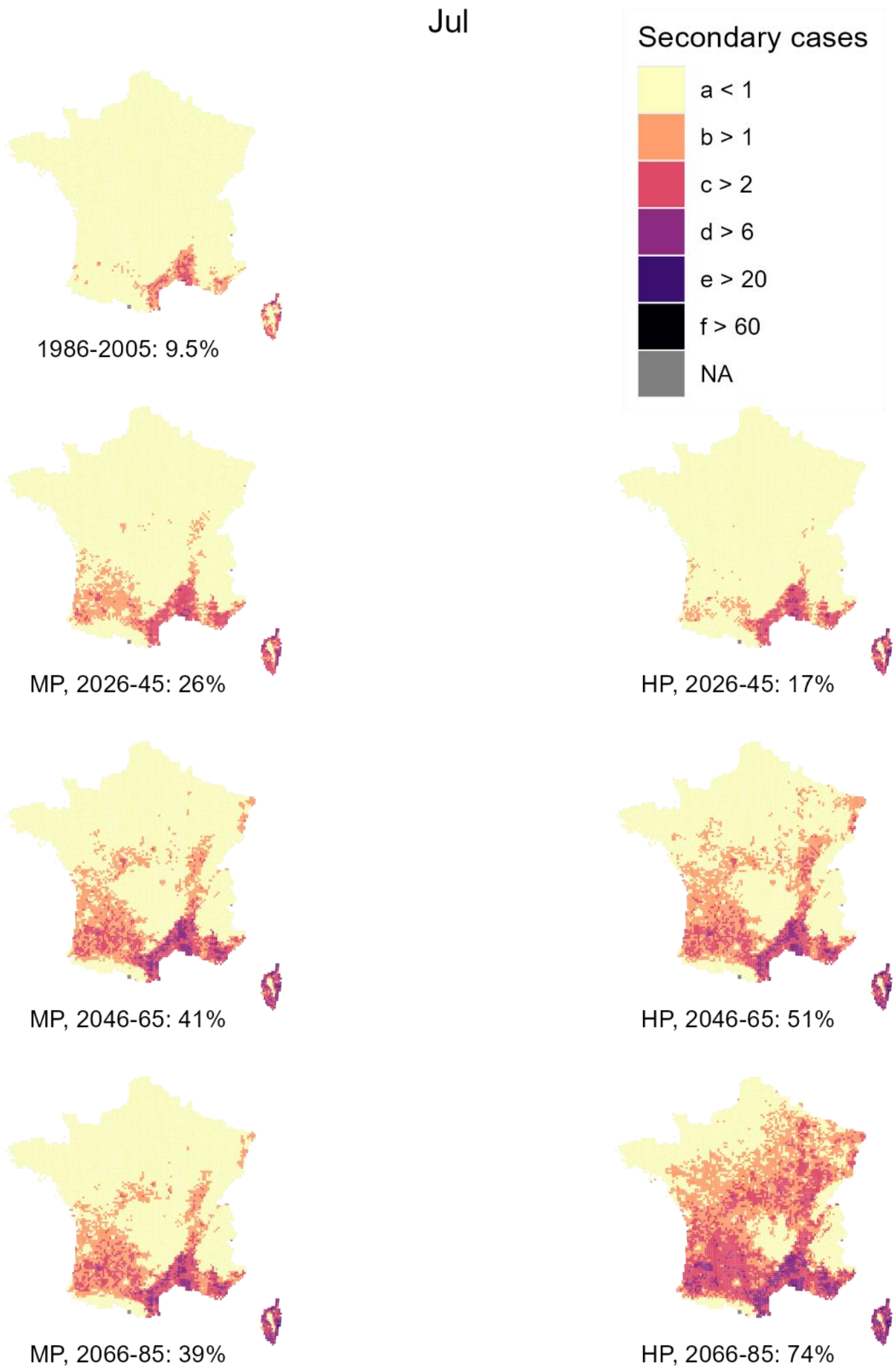


*Figure 6 - Projected number of secondary dengue cases generated in July by the introduction of a positive case on the first day of the month (Intermediate model) for each scenario and time horizon. In brackets, we show the fraction of territory where at least 1 secondary case occurs.*

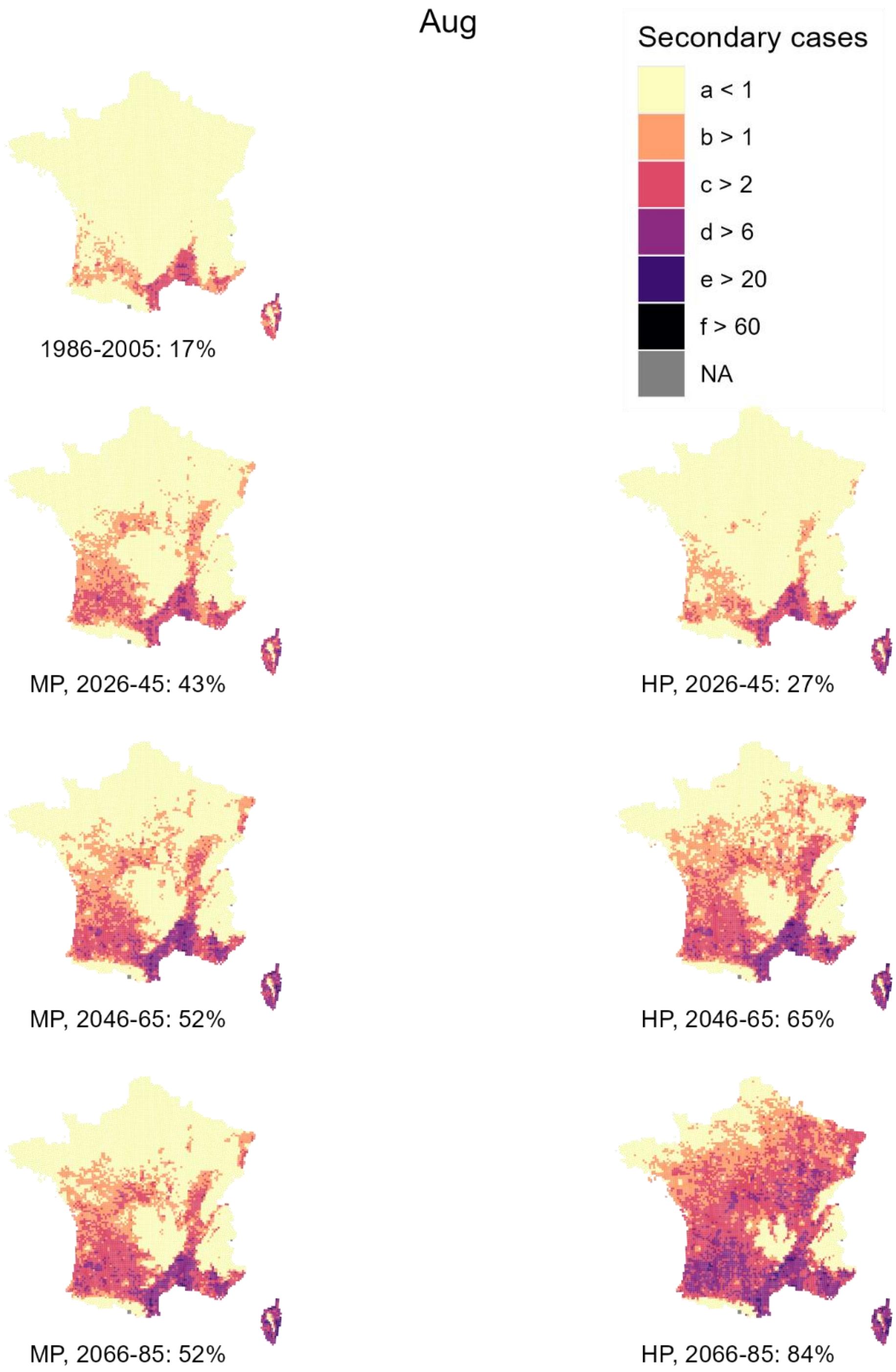


*Figure 7 - Projected number of secondary dengue cases generated in August by the introduction of a positive case on the first day of the month (Intermediate model) for each scenario and time horizon. In brackets, we show the fraction of territory where at least 1 secondary case occurs.*

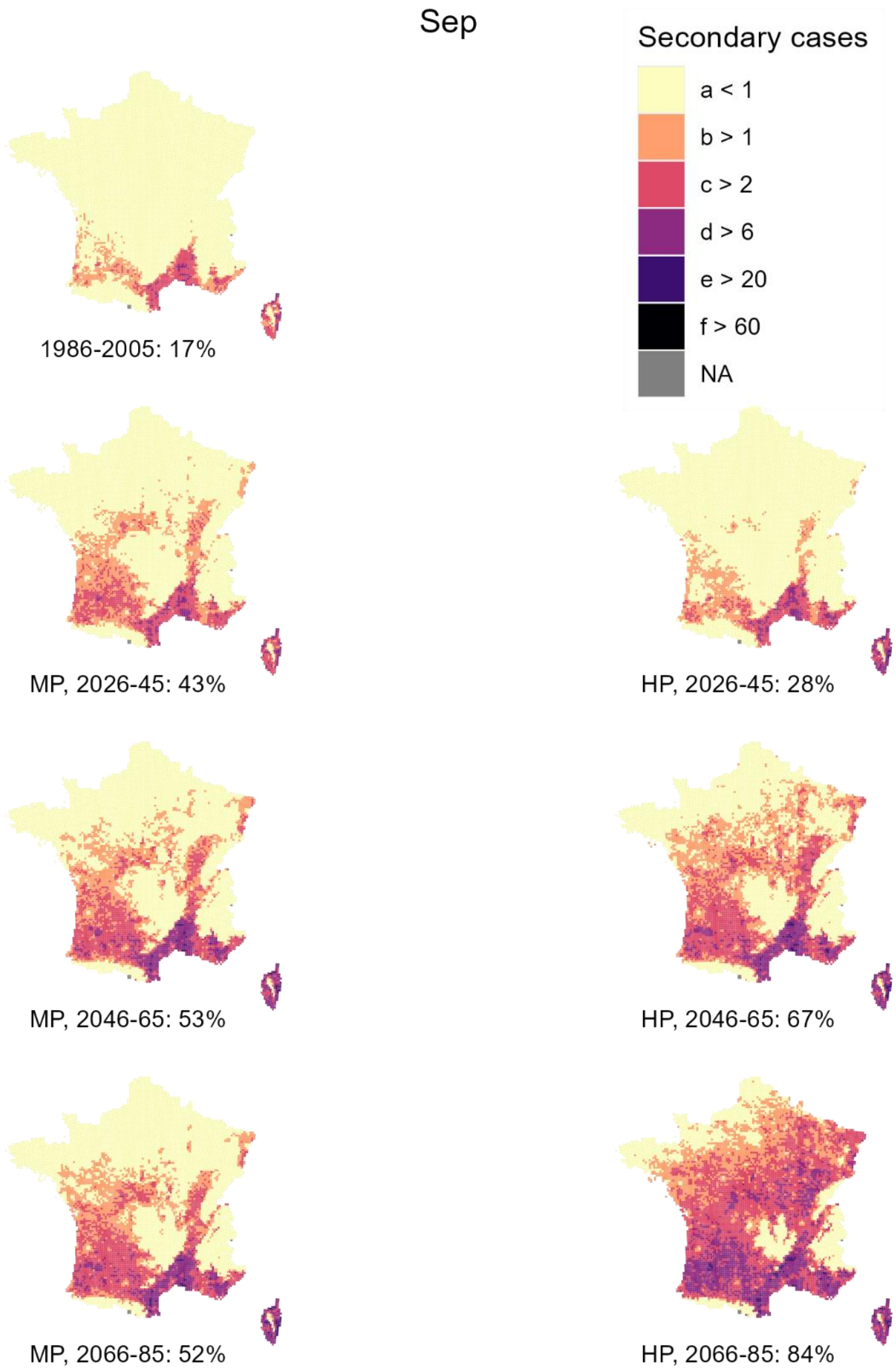


*Figure 8 - Projected number of secondary dengue cases generated in September by the introduction of a positive case on the first day of the month (Intermediate model) for each scenario and time horizon. In brackets, we show the fraction of territory where at least 1 secondary case occurs.*

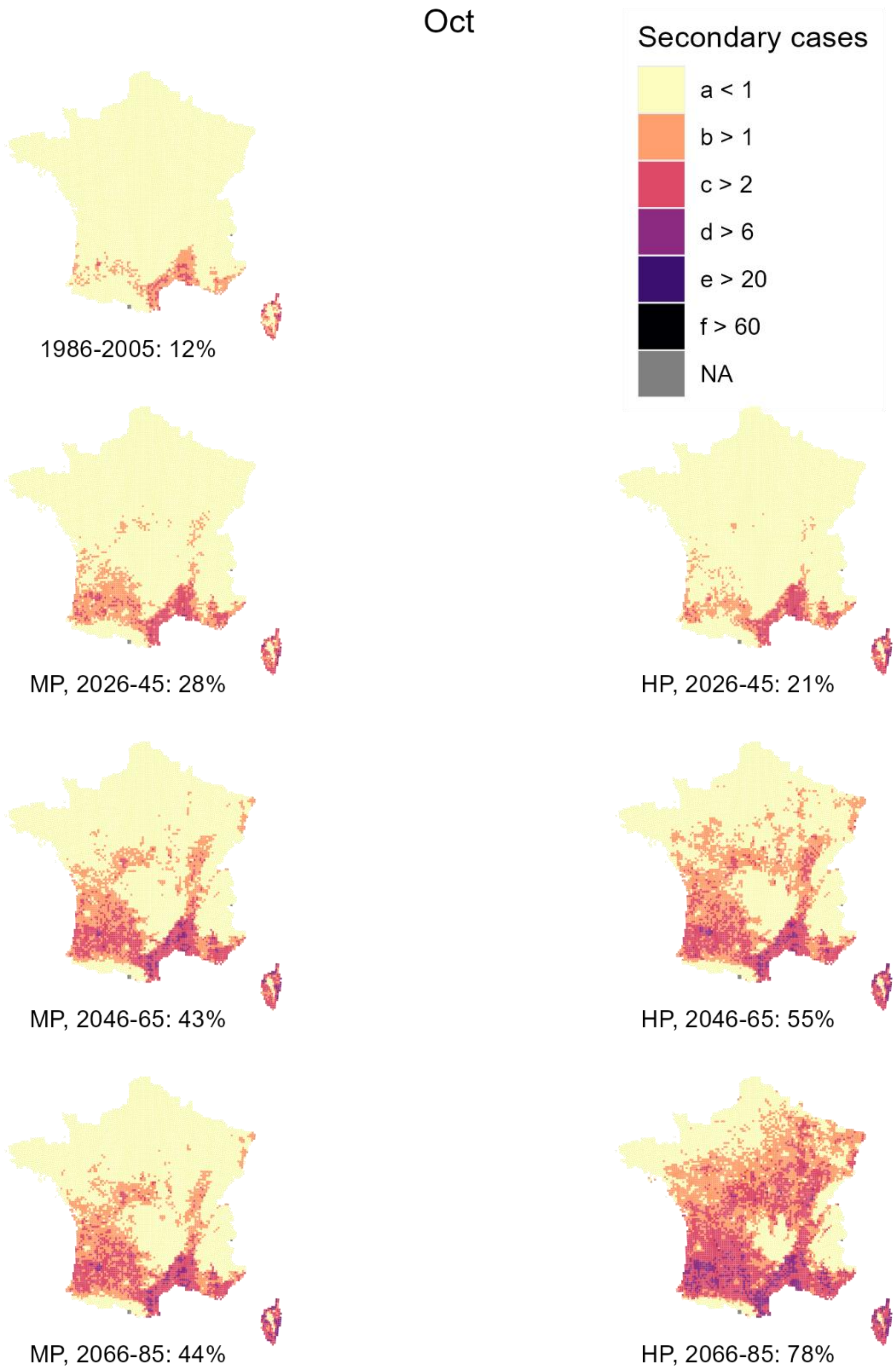


*Figure 9 - Projected number of secondary dengue cases generated in October by the introduction of a positive case on the first day of the month (Intermediate model) for each scenario and time horizon. In brackets, we show the fraction of territory where at least 1 secondary case occurs.*

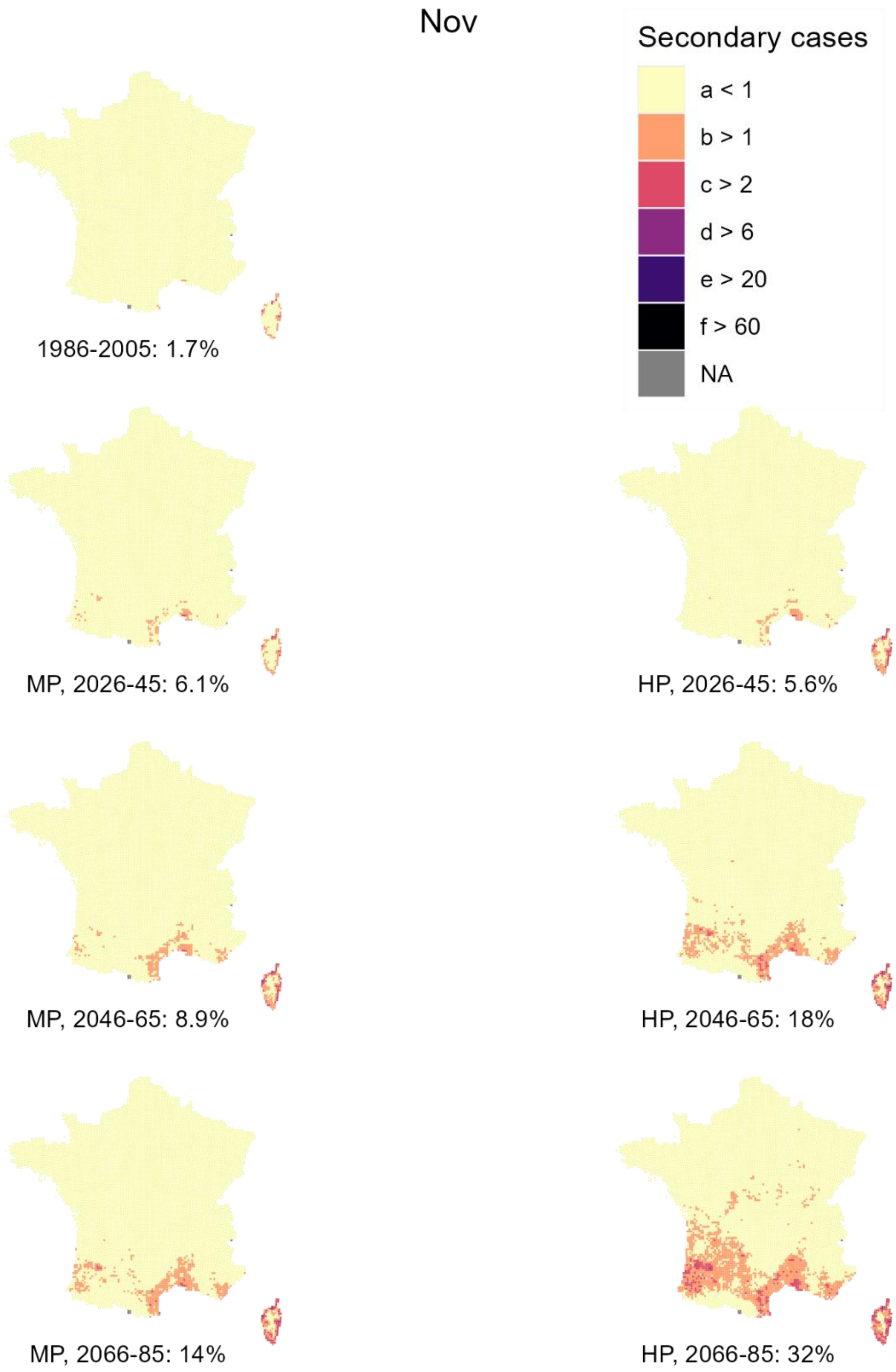


*Figure 10 - Projected number of secondary dengue cases generated in November by the introduction of a positive case on the first day of the month (Intermediate model) for each scenario and time horizon. In brackets, we show the fraction of territory where at least 1 secondary case occurs.*

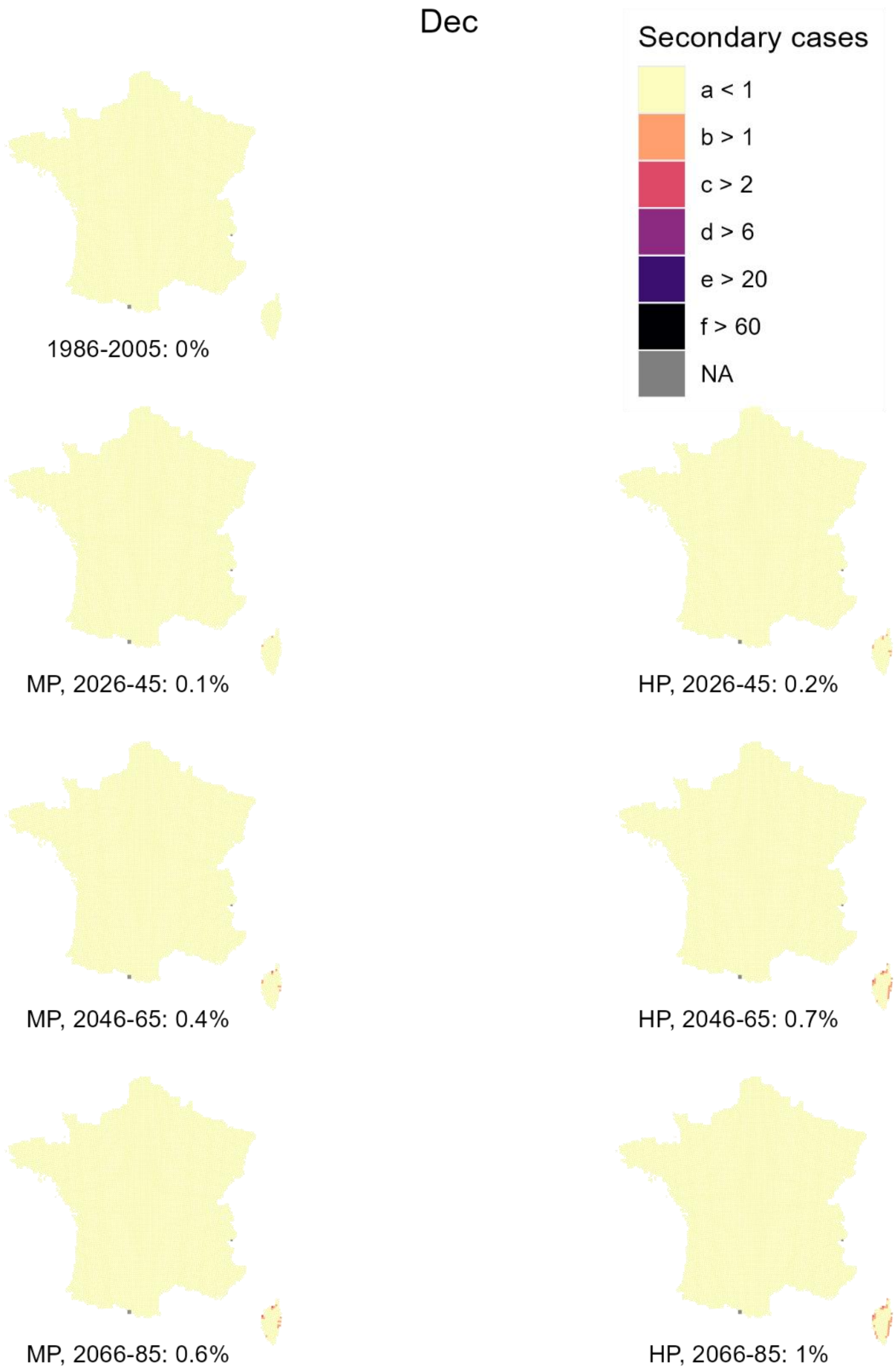


*Figure 11 - Projected number of secondary dengue cases generated in December by the introduction of a positive case on the first day of the month (Intermediate model) for each scenario and time horizon. In brackets, we show the fraction of territory where at least 1 secondary case occurs.*

#### Projected size of a dengue outbreak by month of introduction (warm model)

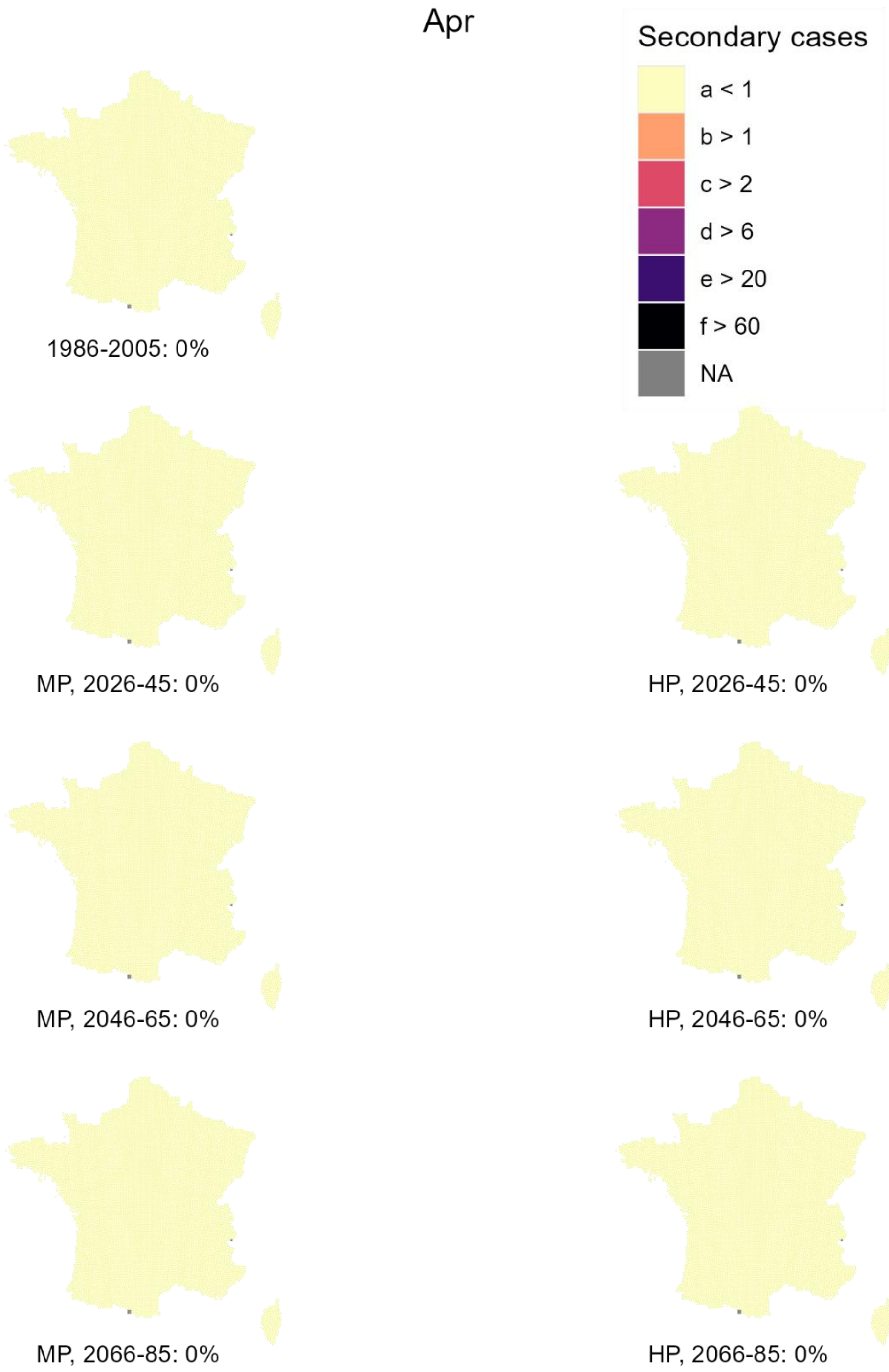


*Figure 12 - Projected number of secondary dengue cases generated in April by the introduction of a positive case on the first day of the month (Warm model) for each scenario and time horizon. In brackets, we show the fraction of territory where at least 1 secondary case occurs.*

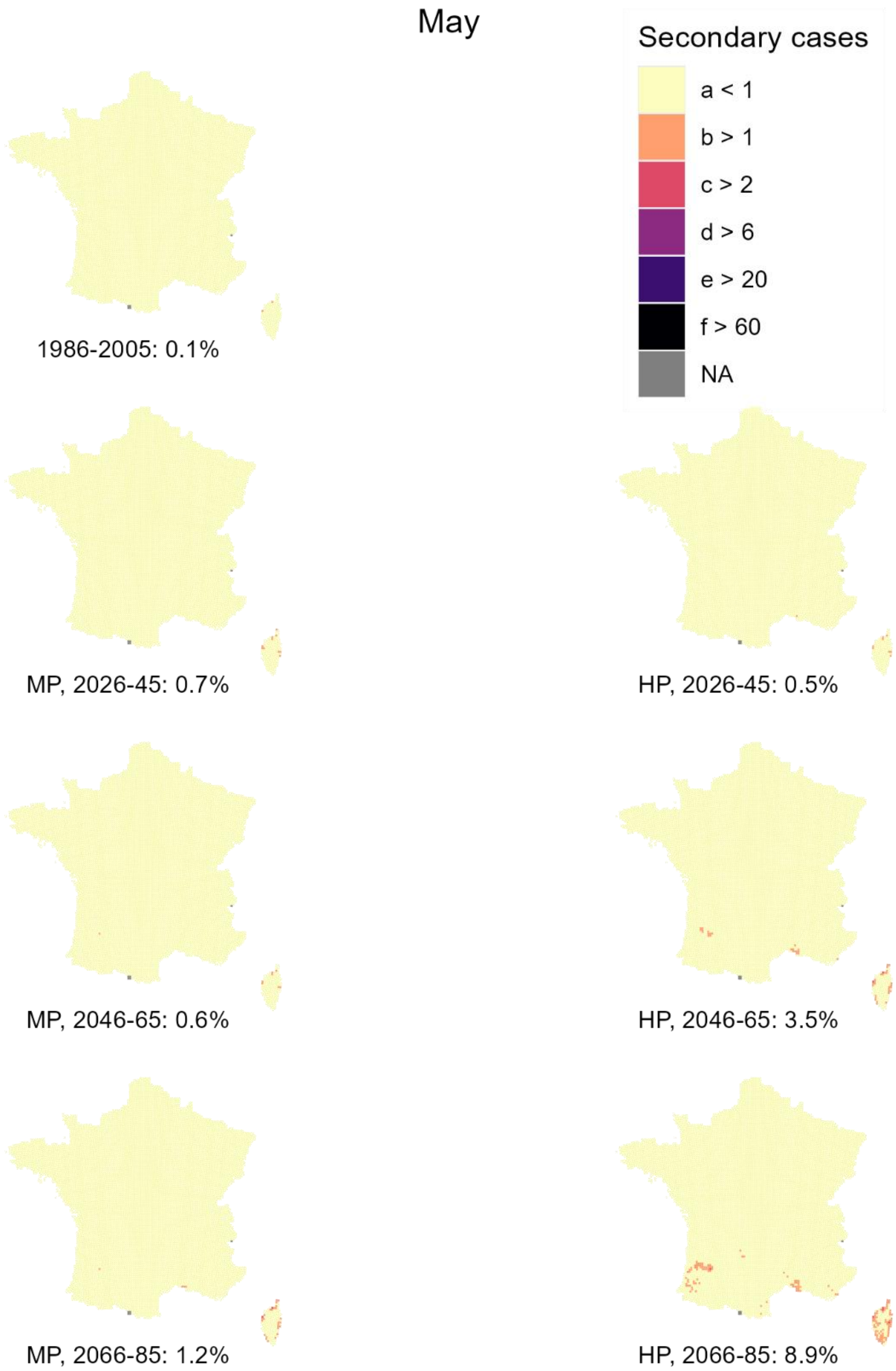


*Figure 13 - Projected number of secondary dengue cases generated in May by the introduction of a positive case on the first day of the month (Warm model) for each scenario and time horizon. In brackets, we show the fraction of territory where at least 1 secondary case occurs.*

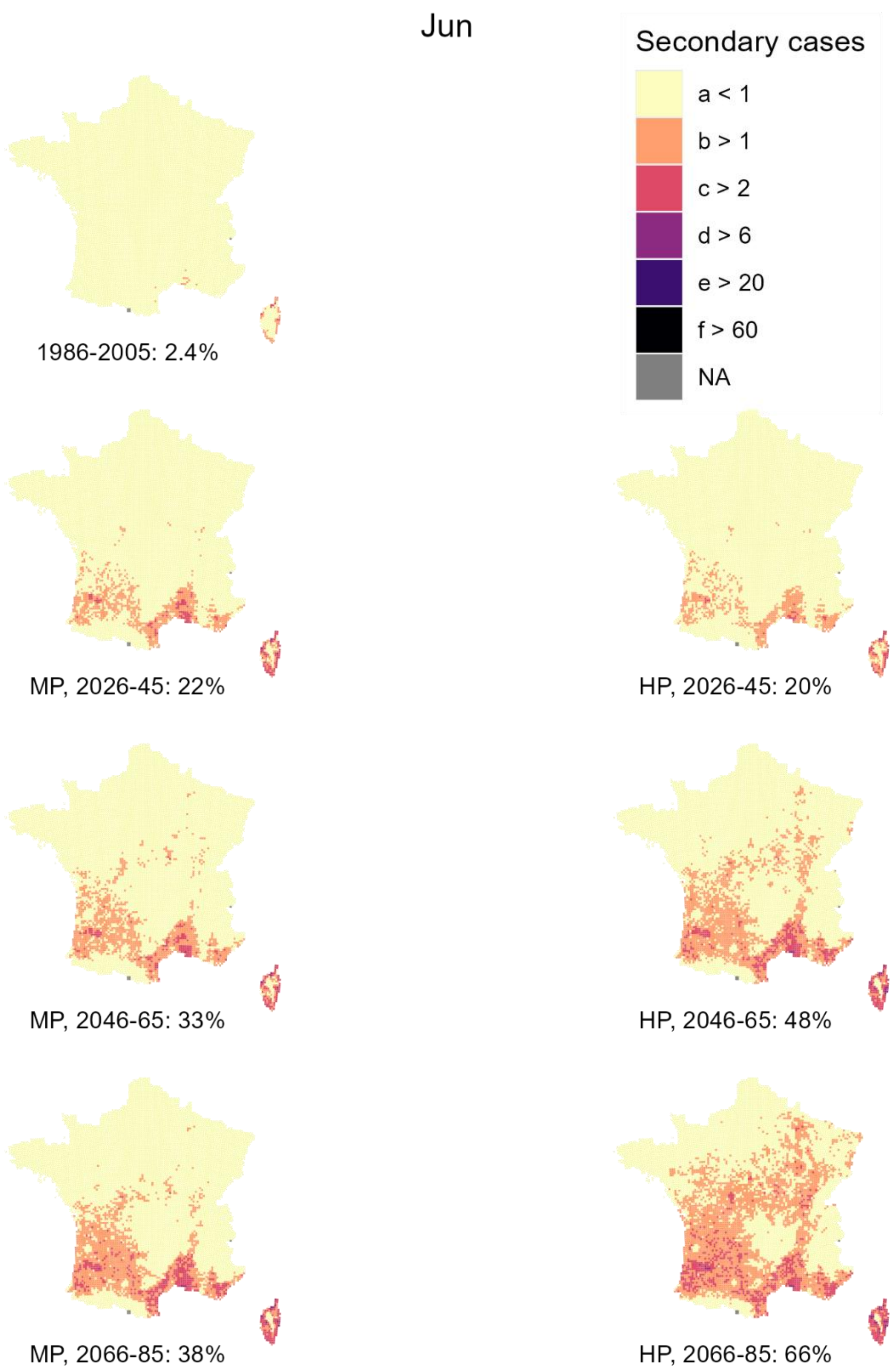


*Figure 14 - Projected number of secondary dengue cases generated in June by the introduction of a positive case on the first day of the month (Warm model) for each scenario and time horizon. In brackets, we show the fraction of territory where at least 1 secondary case occurs.*

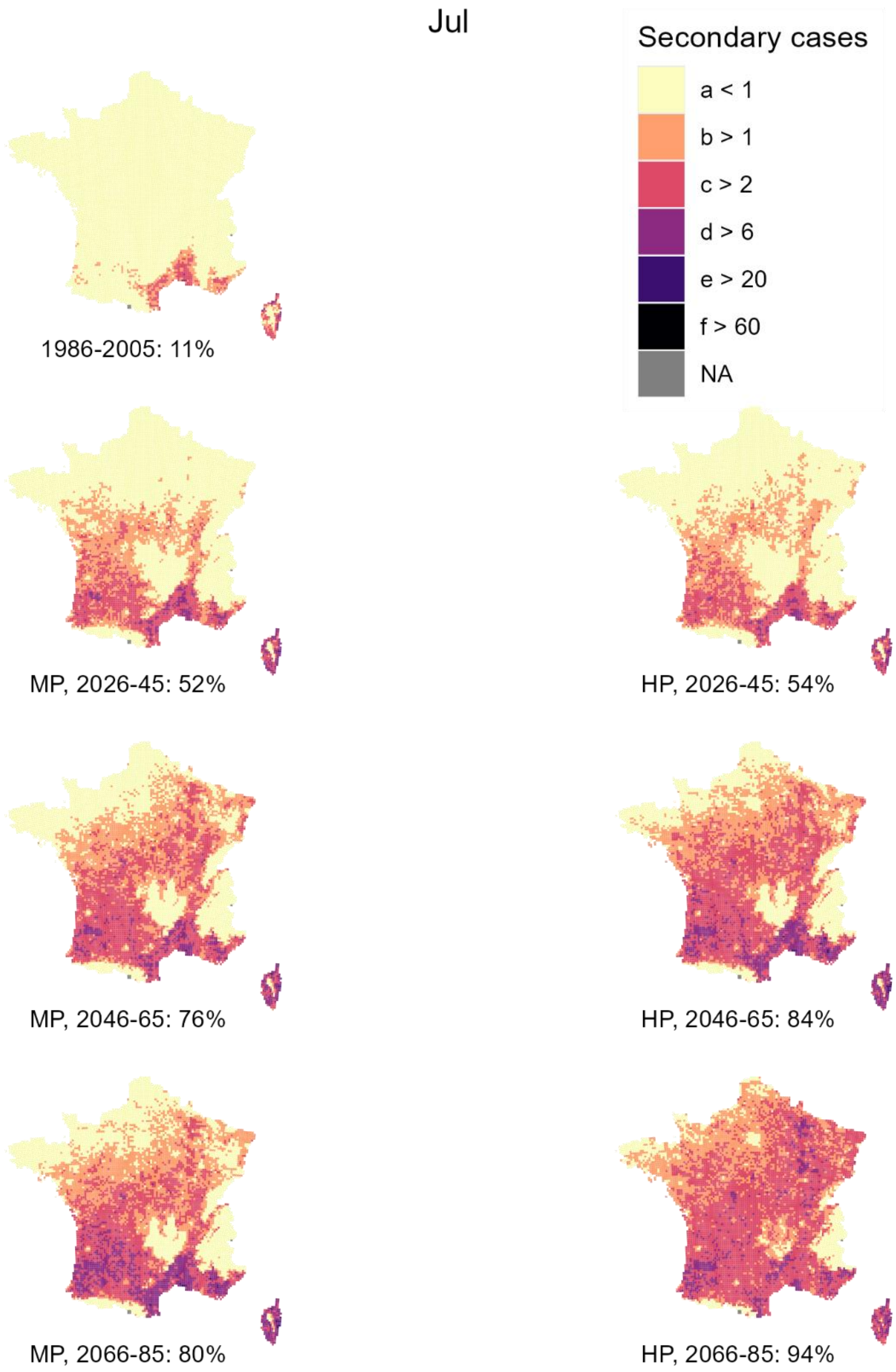


*Figure 15 - Projected number of secondary dengue cases generated in July by the introduction of a positive case on the first day of the month (Warm model) for each scenario and time horizon. In brackets, we show the fraction of territory where at least 1 secondary case occurs.*

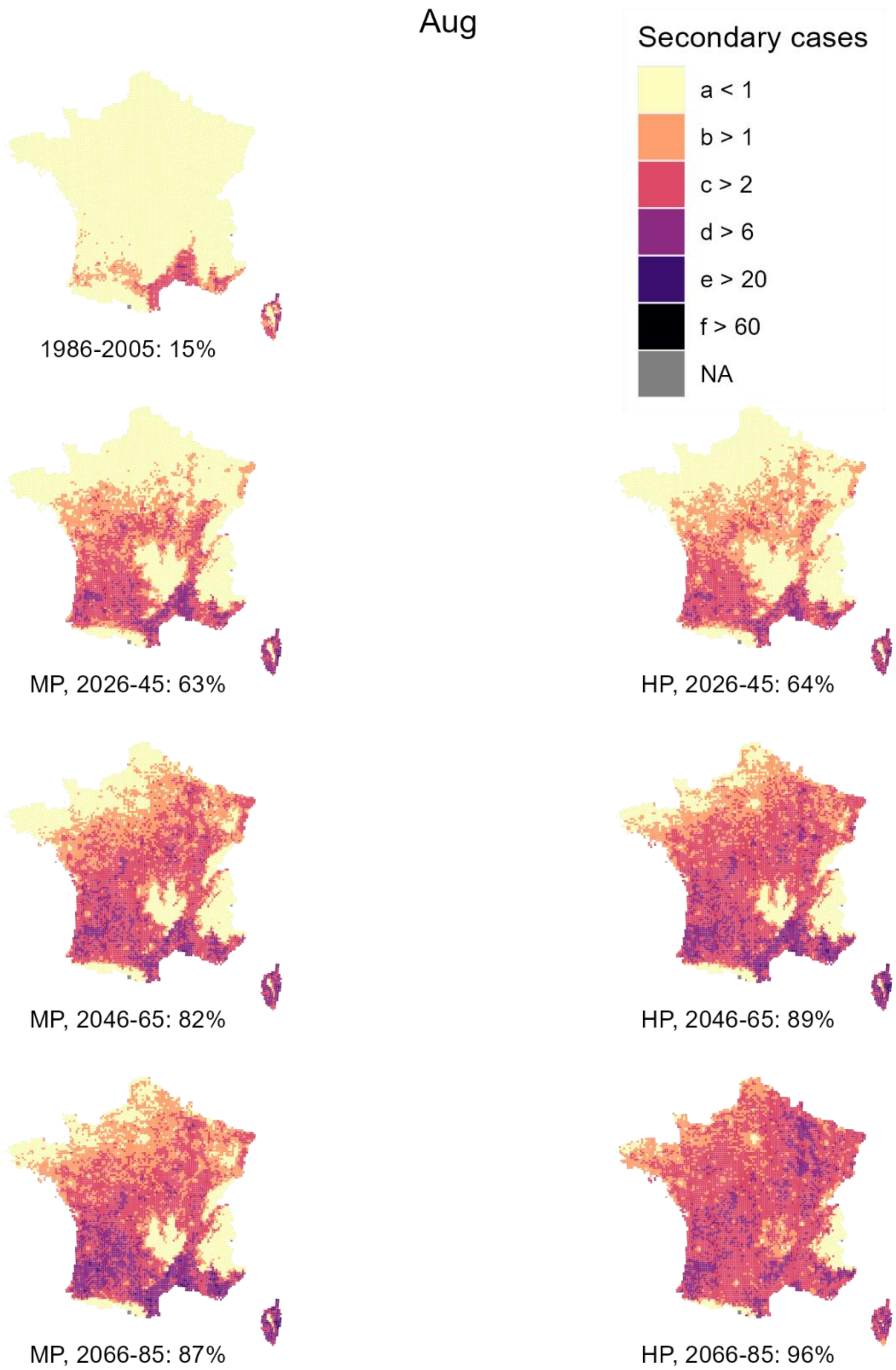


*Figure 16 - Projected number of secondary dengue cases generated in August by the introduction of a positive case on the first day of the month (Warm model) for each scenario and time horizon. In brackets, we show the fraction of territory where at least 1 secondary case occurs.*

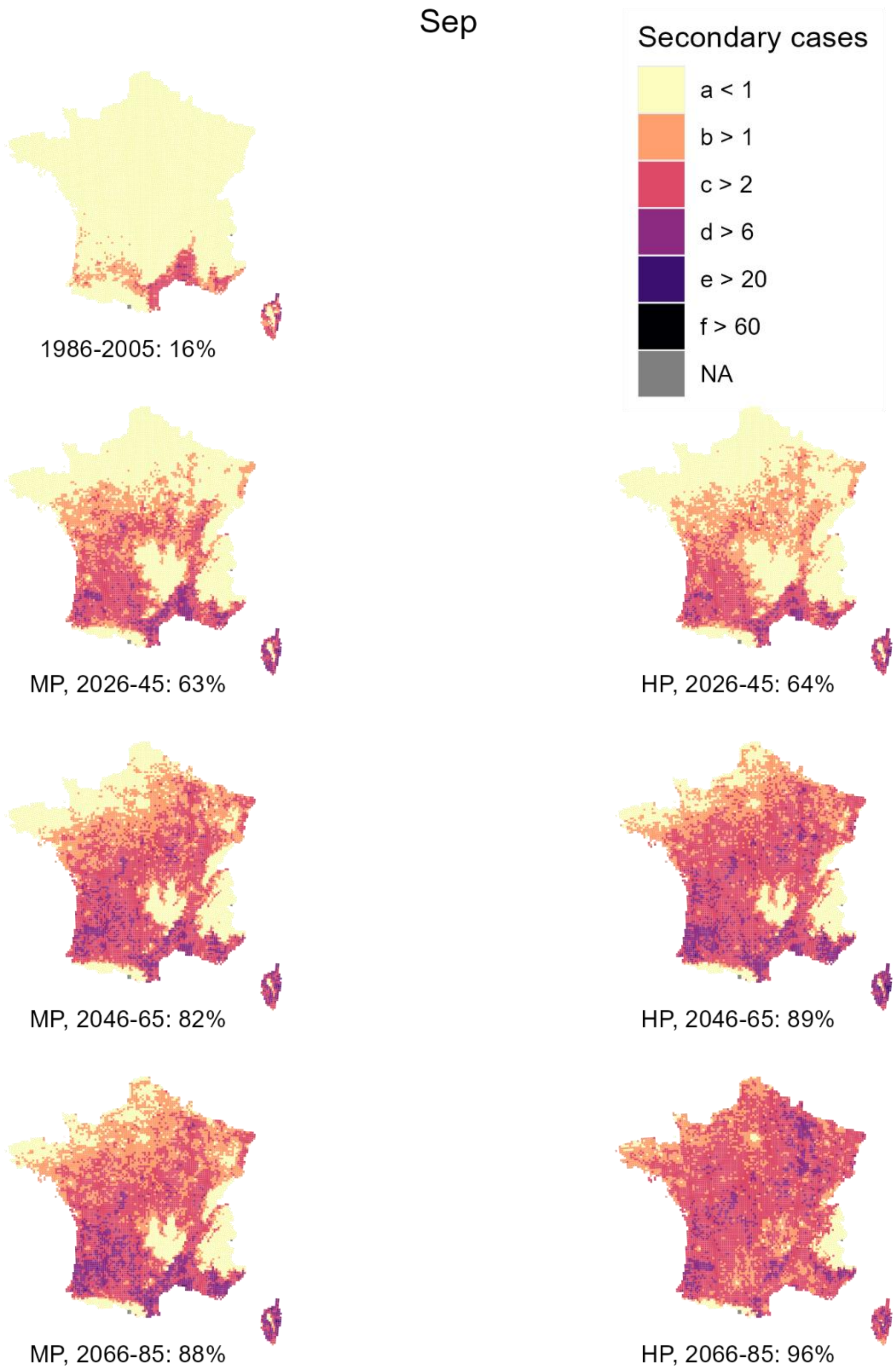


*Figure 17 - Projected number of secondary dengue cases generated in September by the introduction of a positive case on the first day of the month (Warm model) for each scenario and time horizon. In brackets, we show the fraction of territory where at least 1 secondary case occurs.*

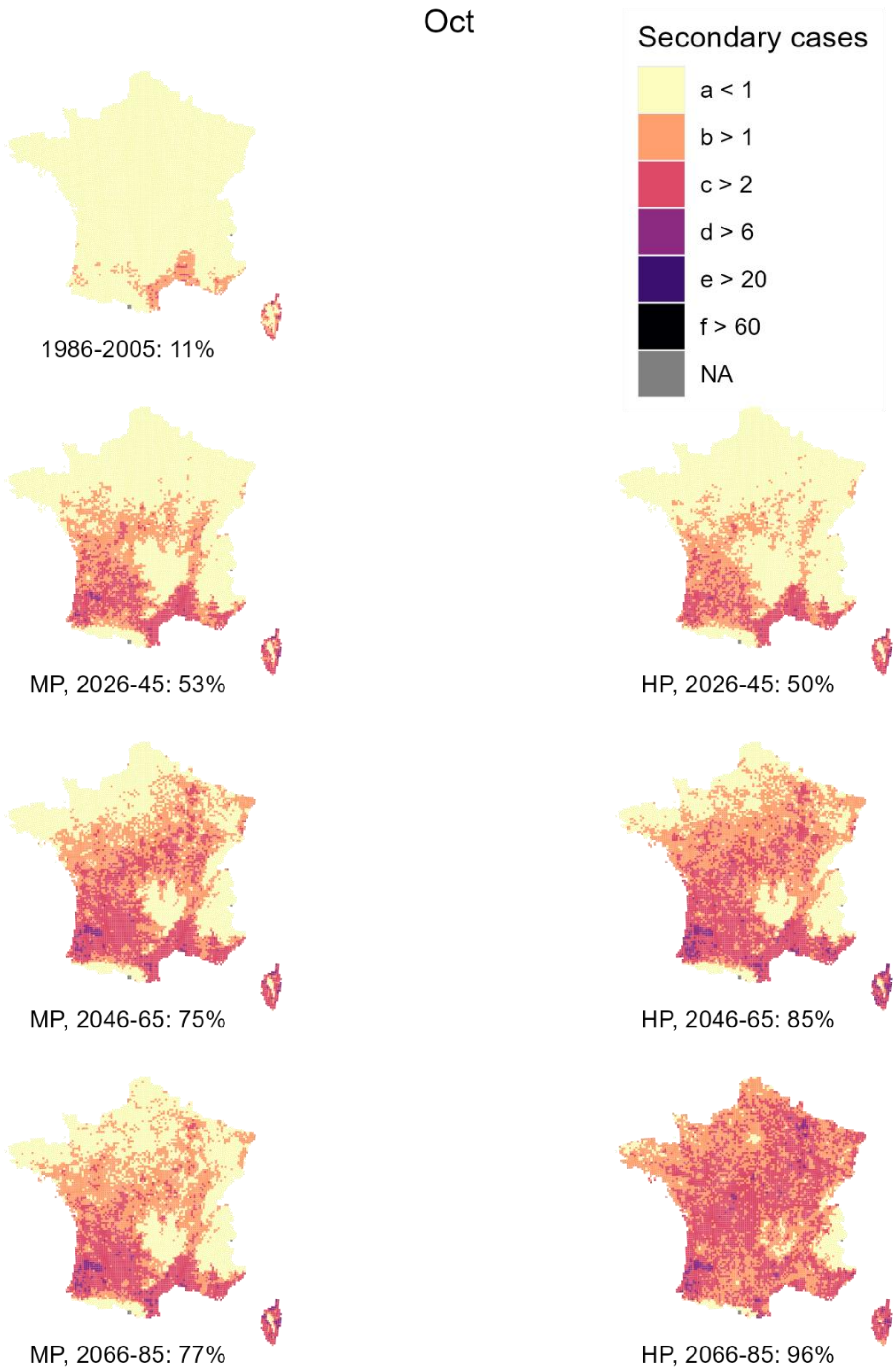


*Figure 18 - Projected number of secondary dengue cases generated in October by the introduction of a positive case on the first day of the month (Warm model) for each scenario and time horizon. In brackets, we show the fraction of territory where at least 1 secondary case occurs.*

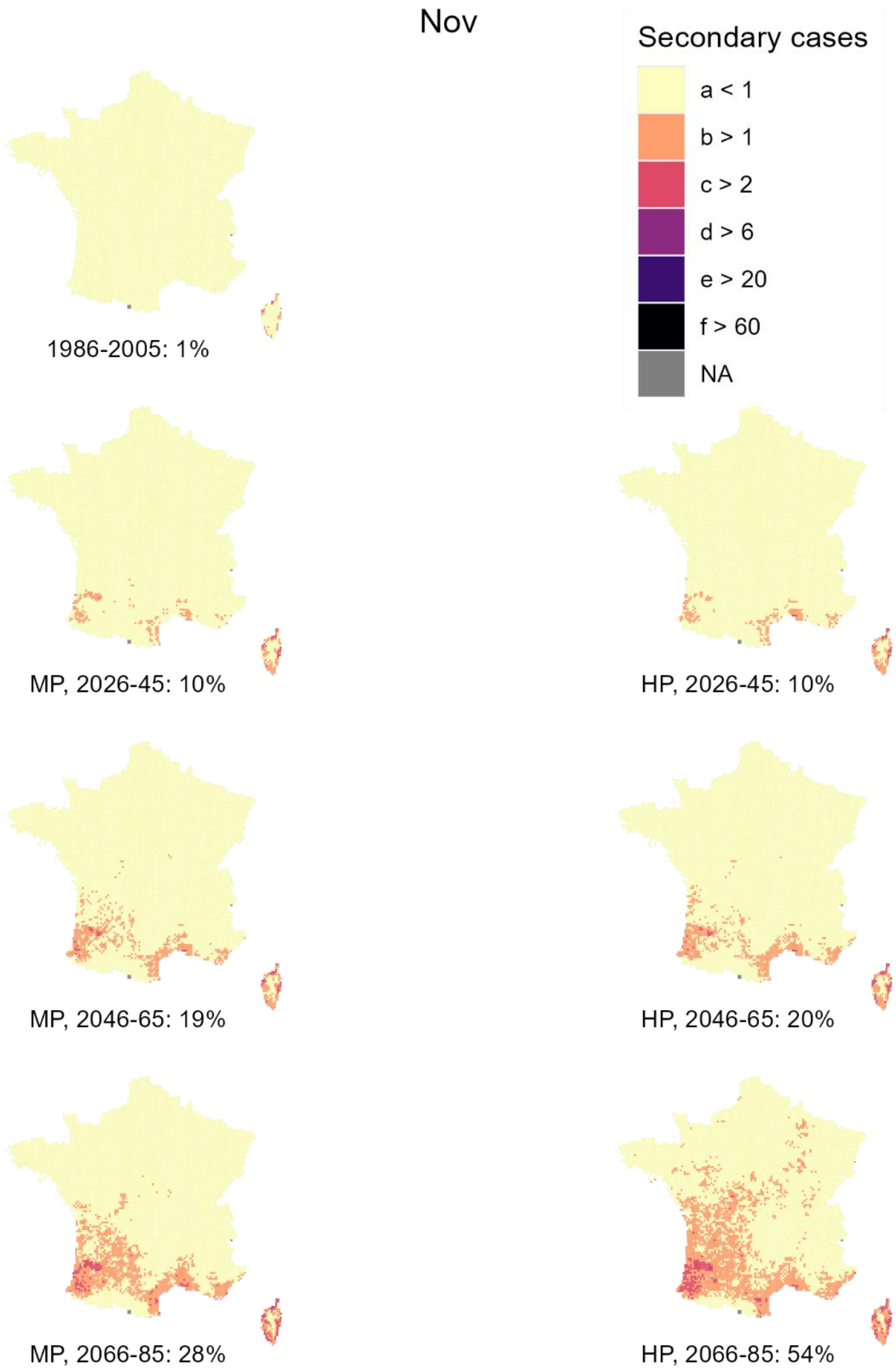


*Figure 19 - Projected number of secondary dengue cases generated in November by the introduction of a positive case on the first day of the month (Warm model) for each scenario and time horizon. In brackets, we show the fraction of territory where at least 1 secondary case occurs.*

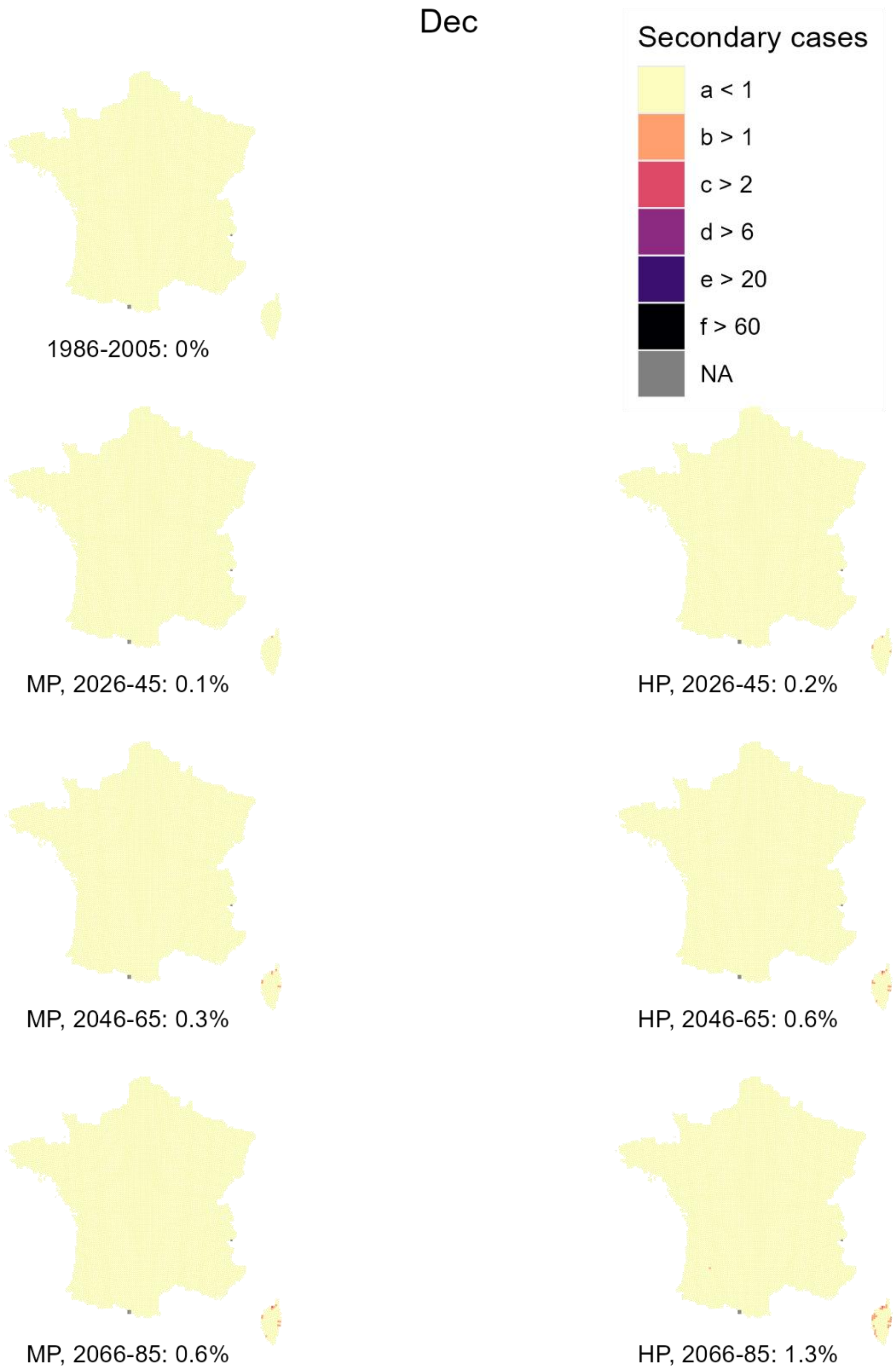


*Figure 20 - Projected number of secondary dengue cases generated in December by the introduction of a positive case on the first day of the month (Warm model) for each scenario and time horizon. In brackets, we show the fraction of territory where at least 1 secondary case occurs.*

#### Projected size of a dengue outbreak by month of introduction (cold model)

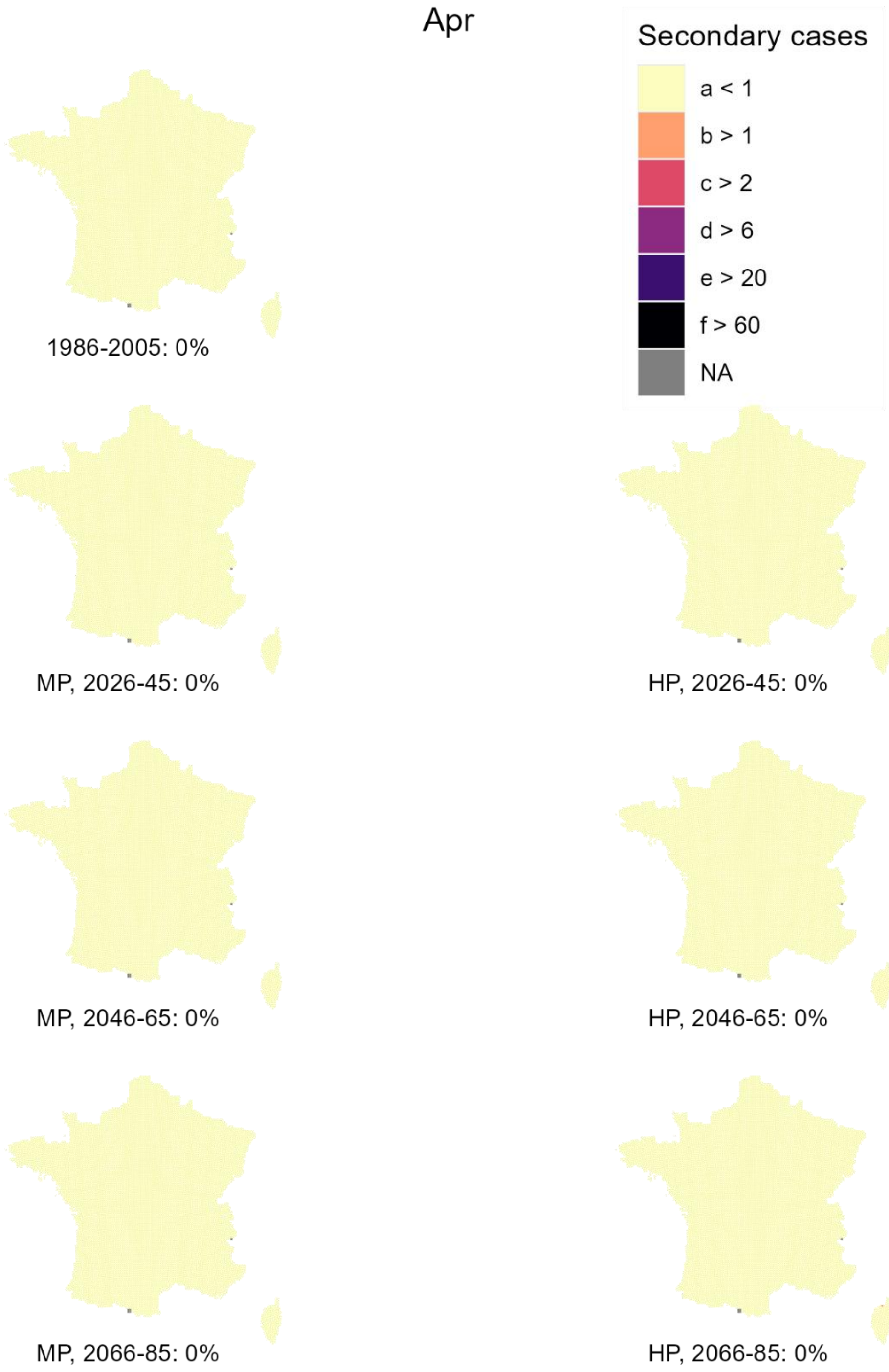


*Figure 21 - Projected number of secondary dengue cases generated in April by the introduction of a positive case on the first day of the month (Cold model) for each scenario and time horizon. In brackets, we show the fraction of territory where at least 1 secondary case occurs.*

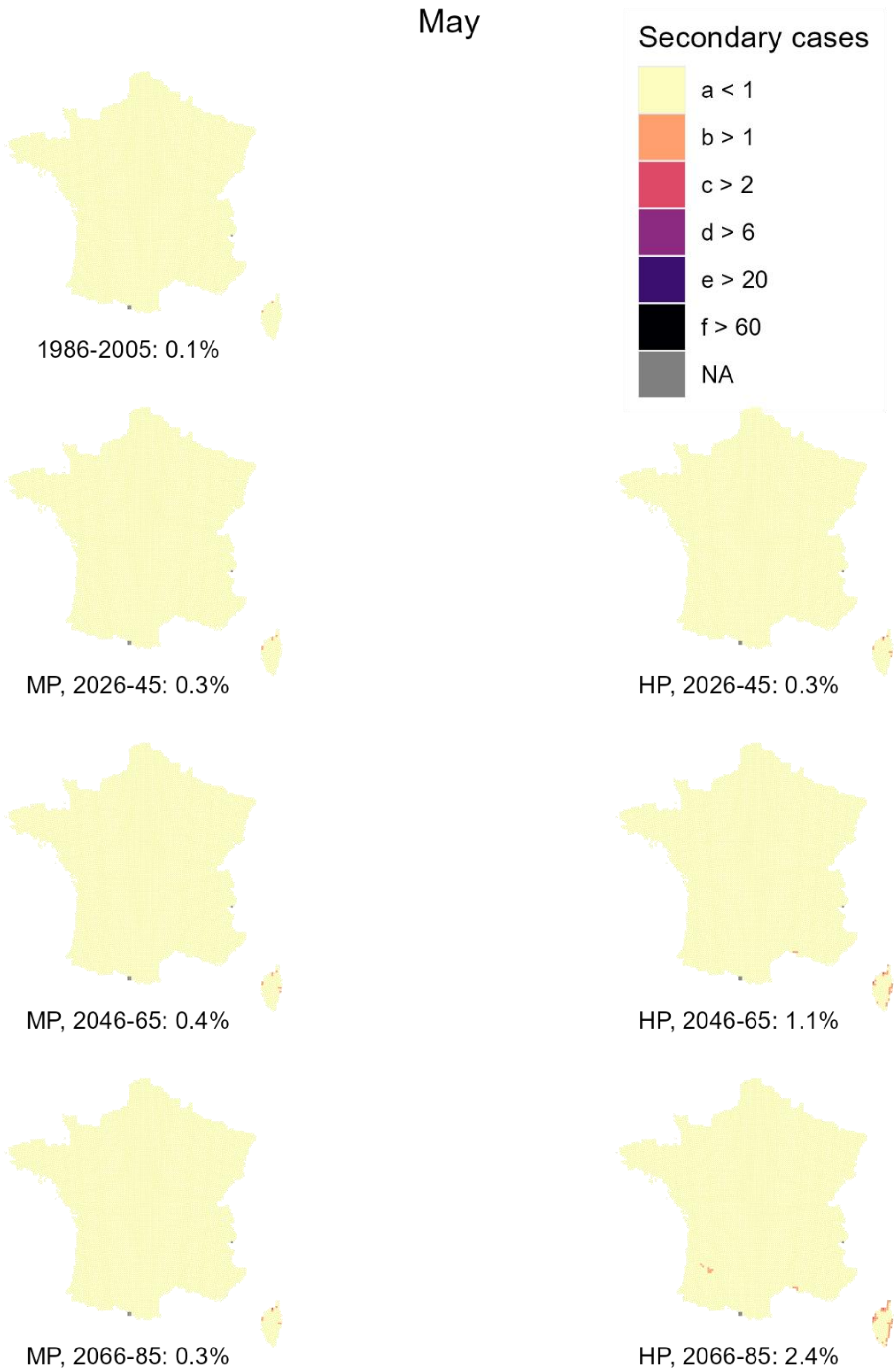


*Figure 22 - Projected number of secondary dengue cases generated in May by the introduction of a positive case on the first day of the month (Cold model) for each scenario and time horizon. In brackets, we show the fraction of territory where at least 1 secondary case occurs.*

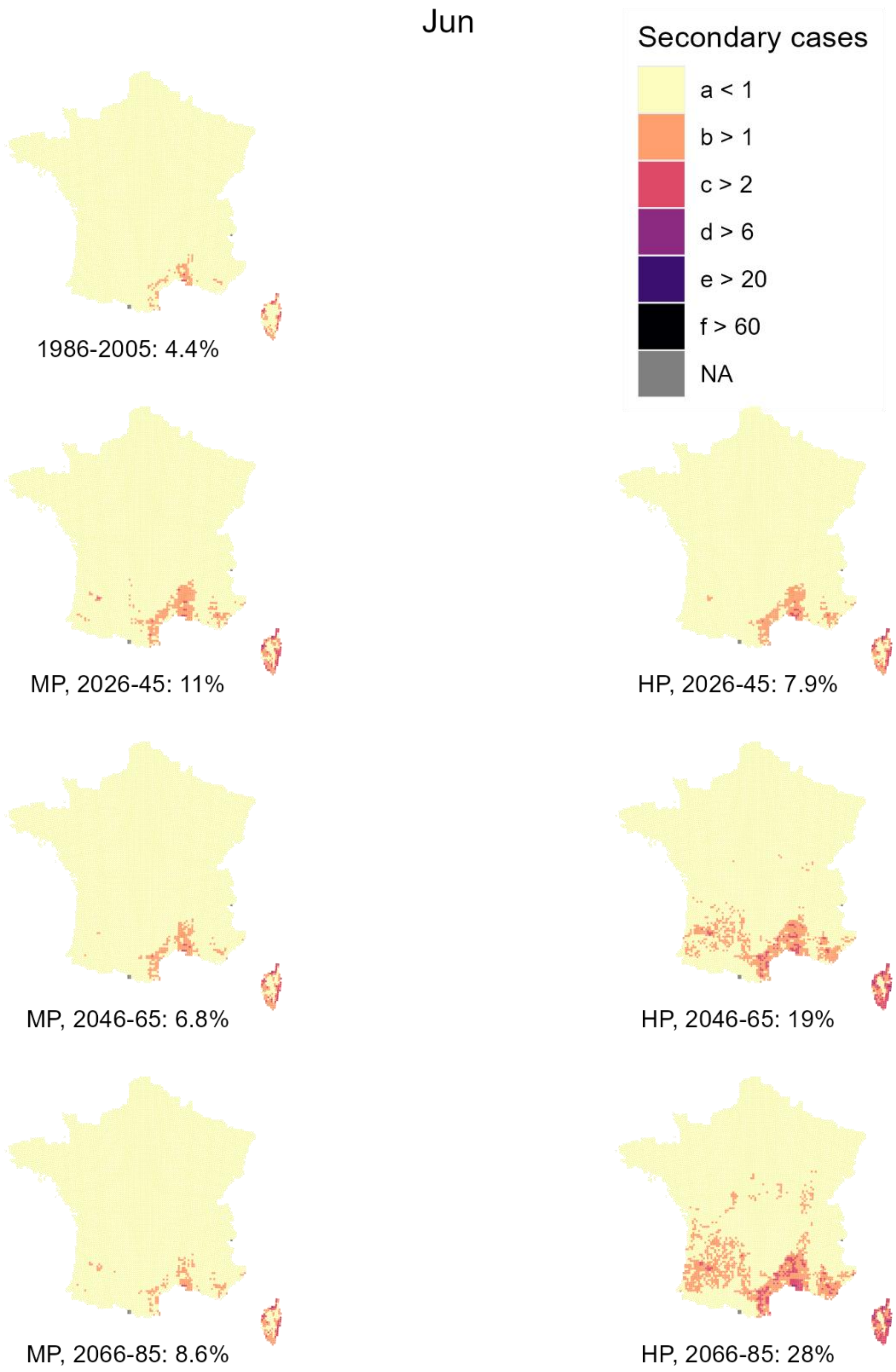


*Figure 23 - Projected number of secondary dengue cases generated in June by the introduction of a positive case on the first day of the month (Cold model) for each scenario and time horizon. In brackets, we show the fraction of territory where at least 1 secondary case occurs.*

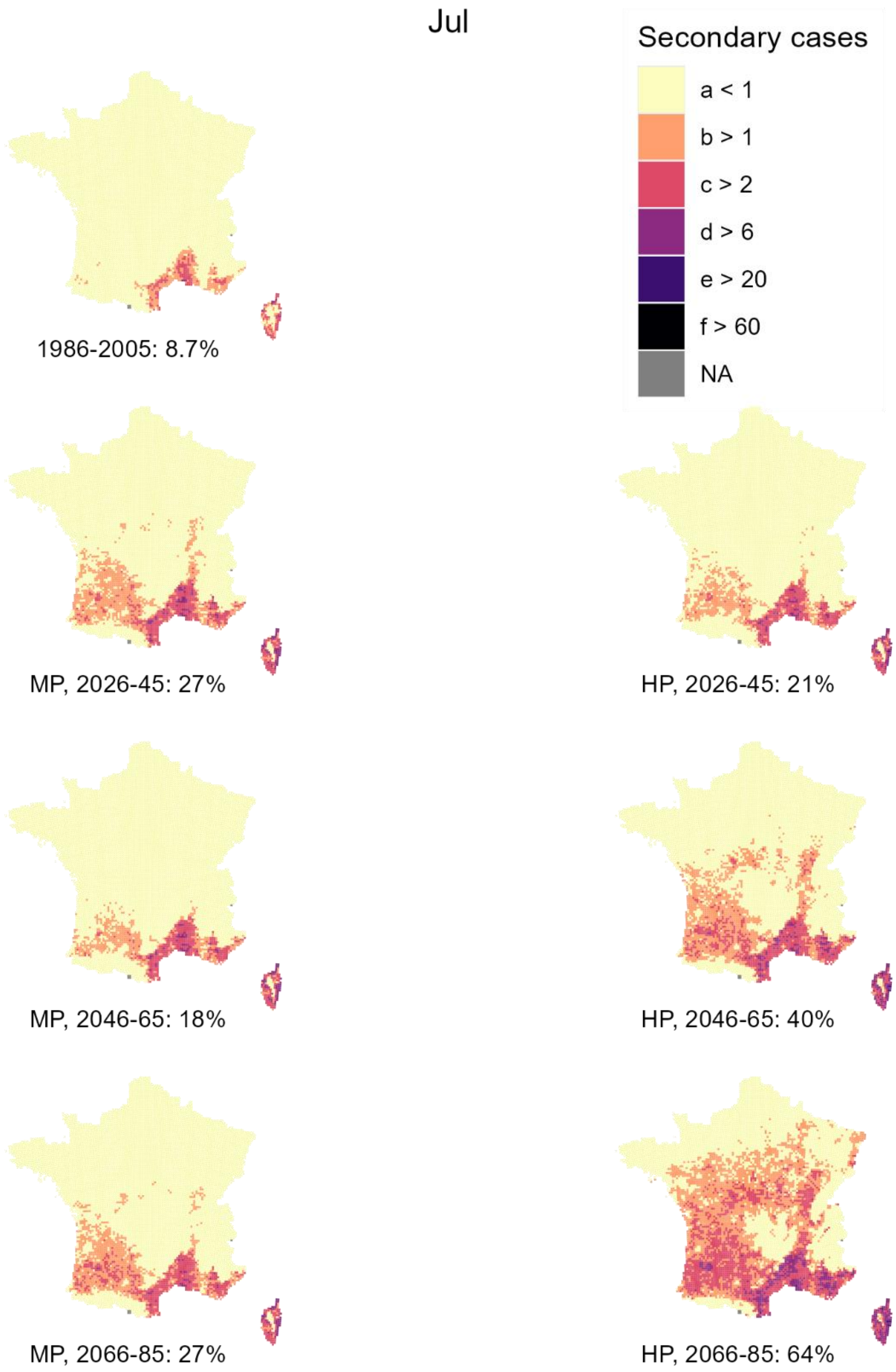


*Figure 24 - Projected number of secondary dengue cases generated in July by the introduction of a positive case on the first day of the month (Cold model) for each scenario and time horizon. In brackets, we show the fraction of territory where at least 1 secondary case occurs.*

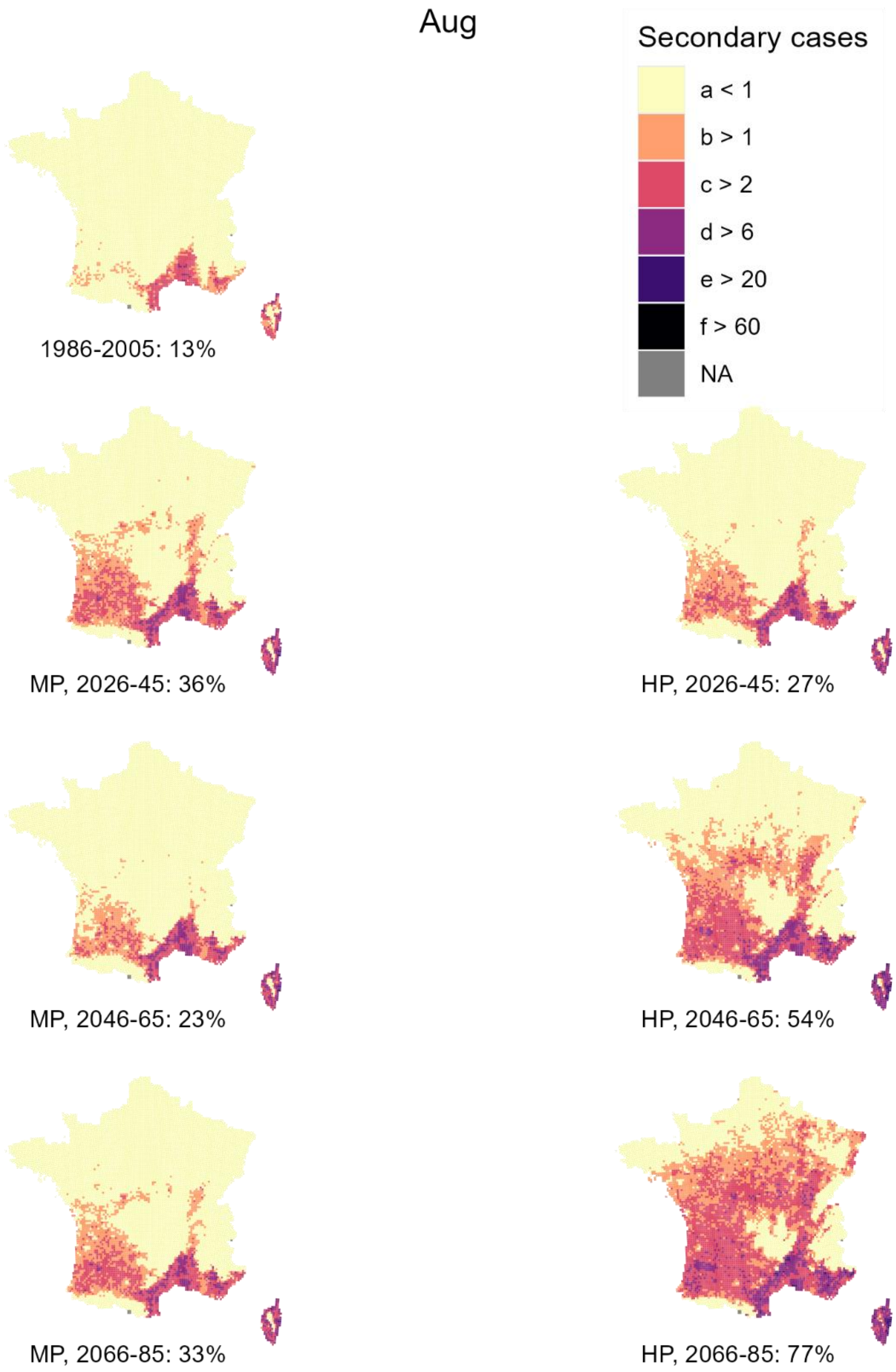


*Figure 25 - Projected number of secondary dengue cases generated in August by the introduction of a positive case on the first day of the month (Cold model) for each scenario and time horizon. In brackets, we show the fraction of territory where at least 1 secondary case occurs.*

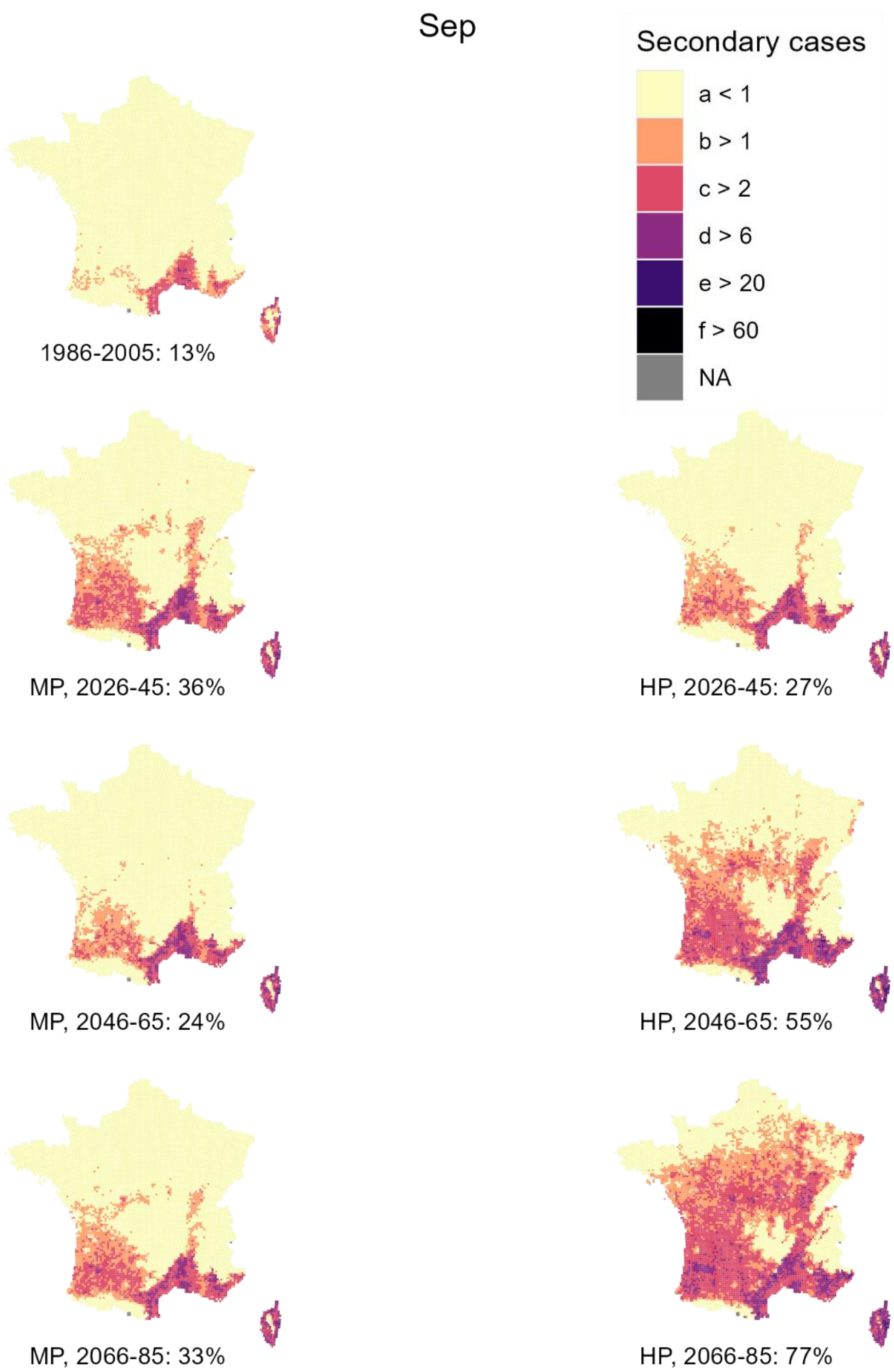


*Figure 26 - Projected number of secondary dengue cases generated in September by the introduction of a positive case on the first day of the month (Cold model) for each scenario and time horizon. In brackets, we show the fraction of territory where at least 1 secondary case occurs.*

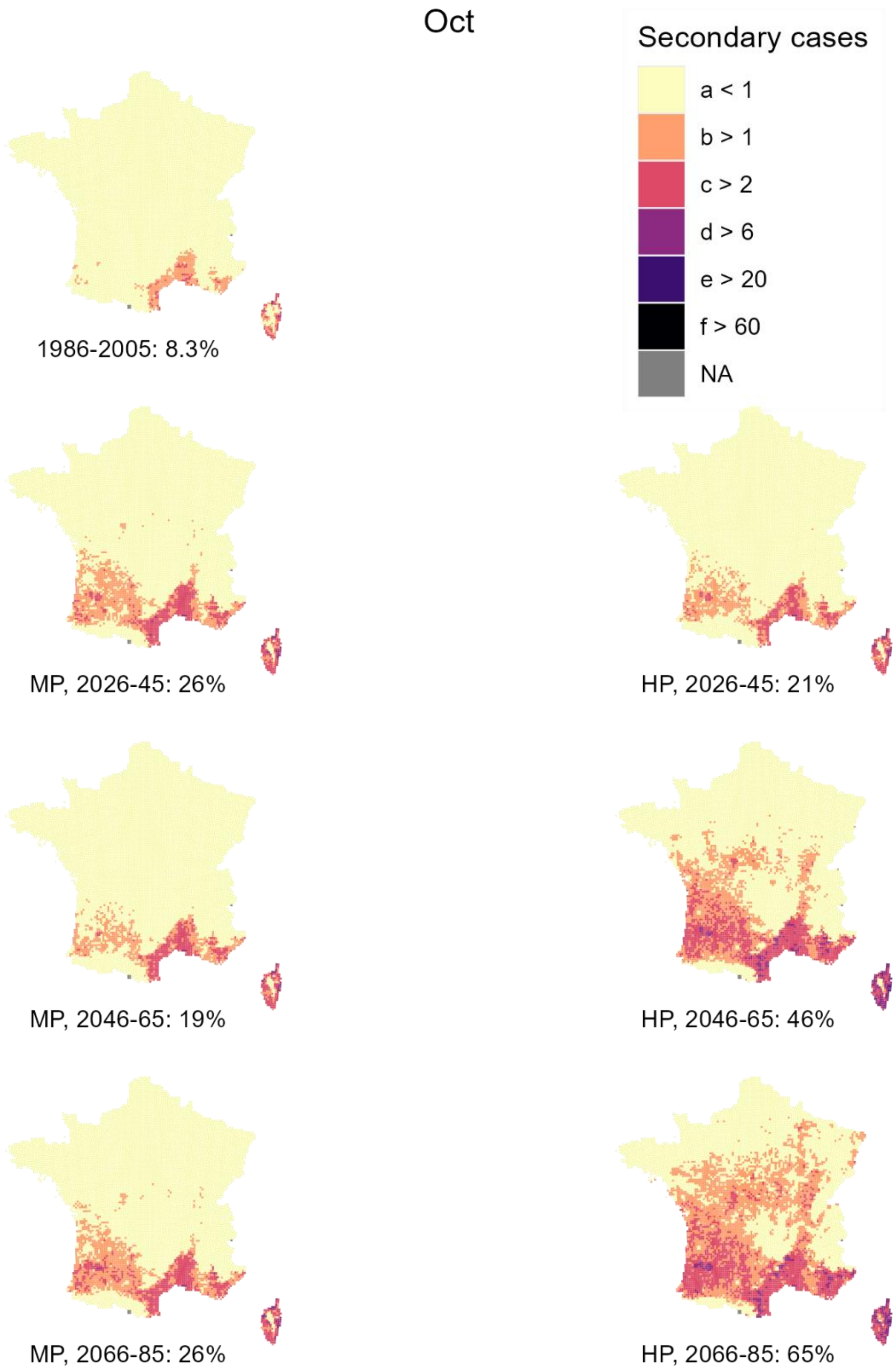


*Figure 27 - Projected number of secondary dengue cases generated in October by the introduction of a positive case on the first day of the month (Cold model) for each scenario and time horizon. In brackets, we show the fraction of territory where at least 1 secondary case occurs.*

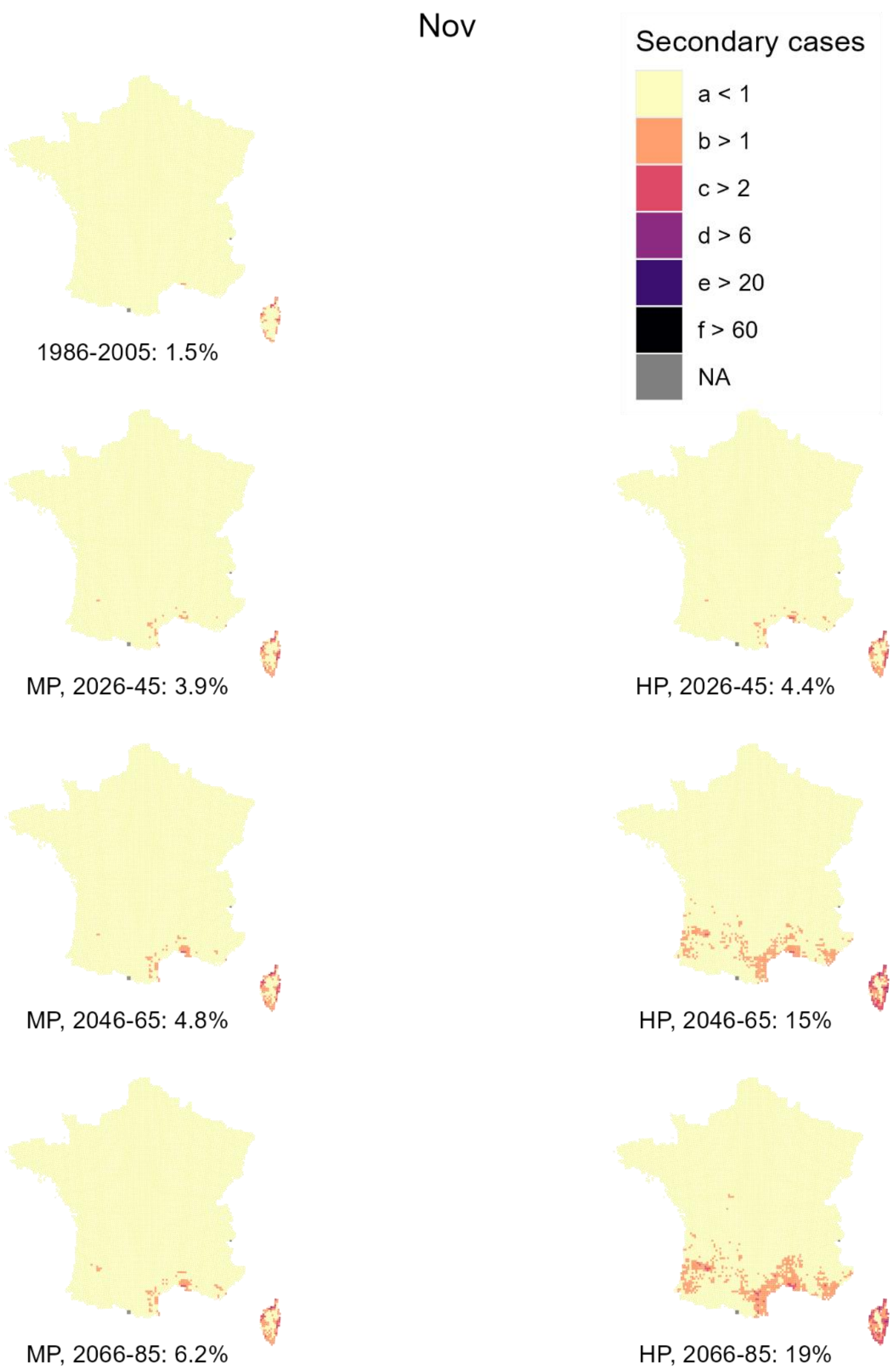


*Figure 28 - Projected number of secondary dengue cases generated in November by the introduction of a positive case on the first day of the month (Cold model) for each scenario and time horizon. In brackets, we show the fraction of territory where at least 1 secondary case occurs.*

141

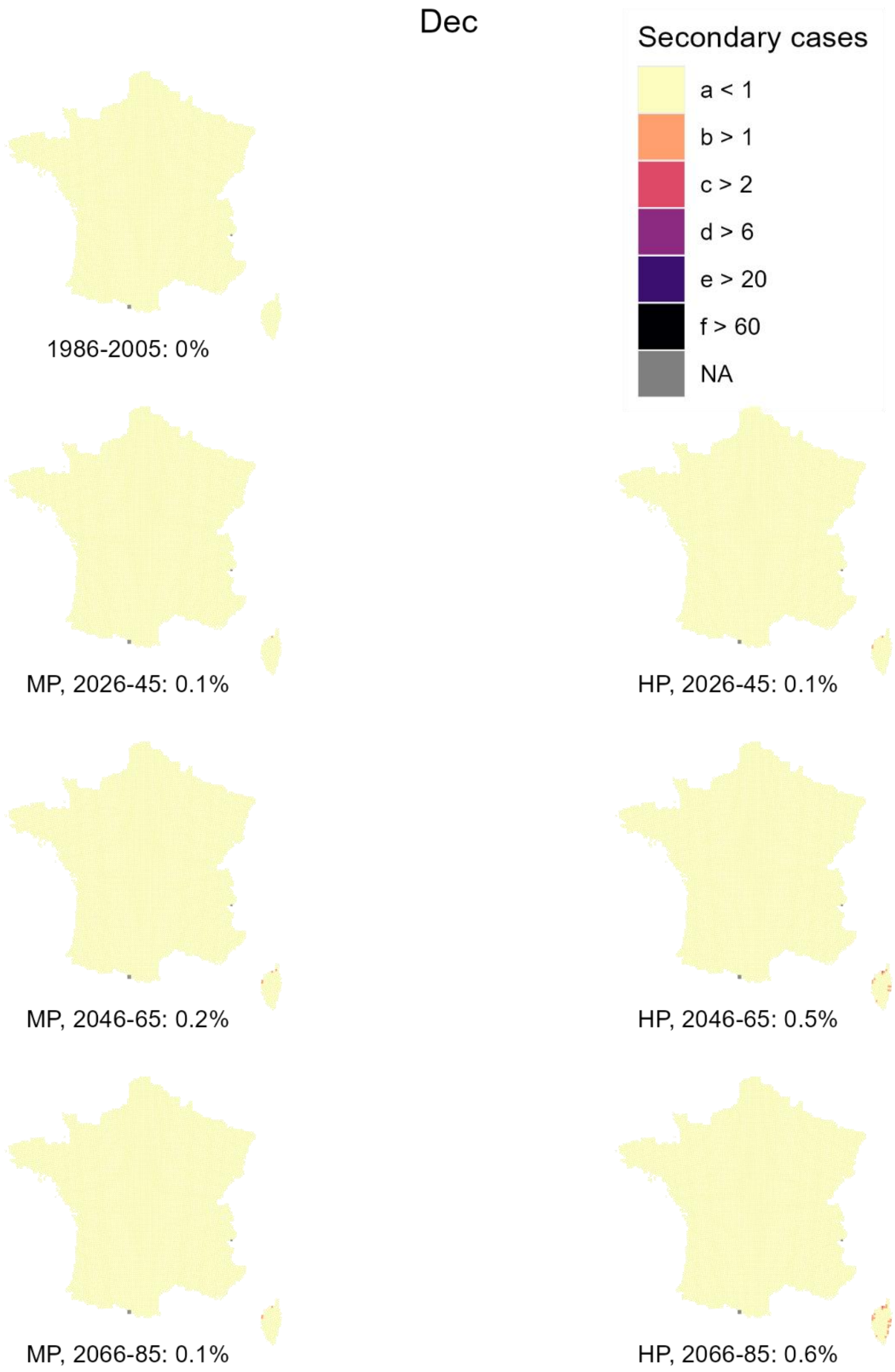


142 *Figure 29 - Projected number of secondary dengue cases generated in December by the introduction of a positive case on*
143 *the first day of the month (Cold model) for each scenario and time horizon. In brackets, we show the fraction of territory*
144 *where at least 1 secondary case occurs.*

#### Projected climatic suitability for a homodynamic (non-diapausing) Ae. albopictus population

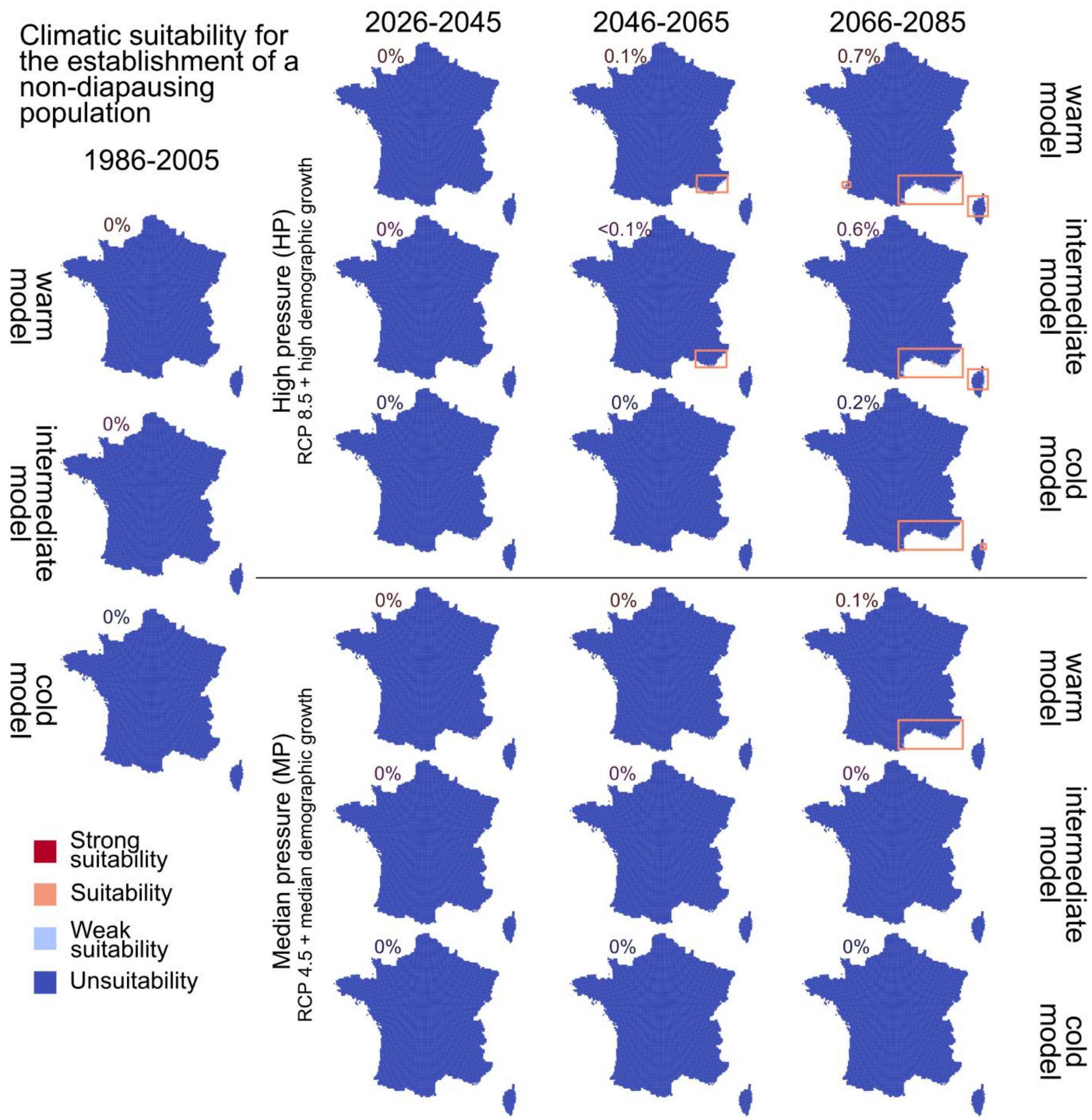


*Figure 30 - Projection of the climatic suitability for the establishment of Ae. albopictus without diapause. Strong suitability: $A_0 > 10$; Suitability: $A_0 > 1$; weak suitability: $A_0 > 0.1$; unsuitable otherwise. % of territory either "suitable" or "strongly suitable"*

#### Projected seasonality of mosquito dynamics

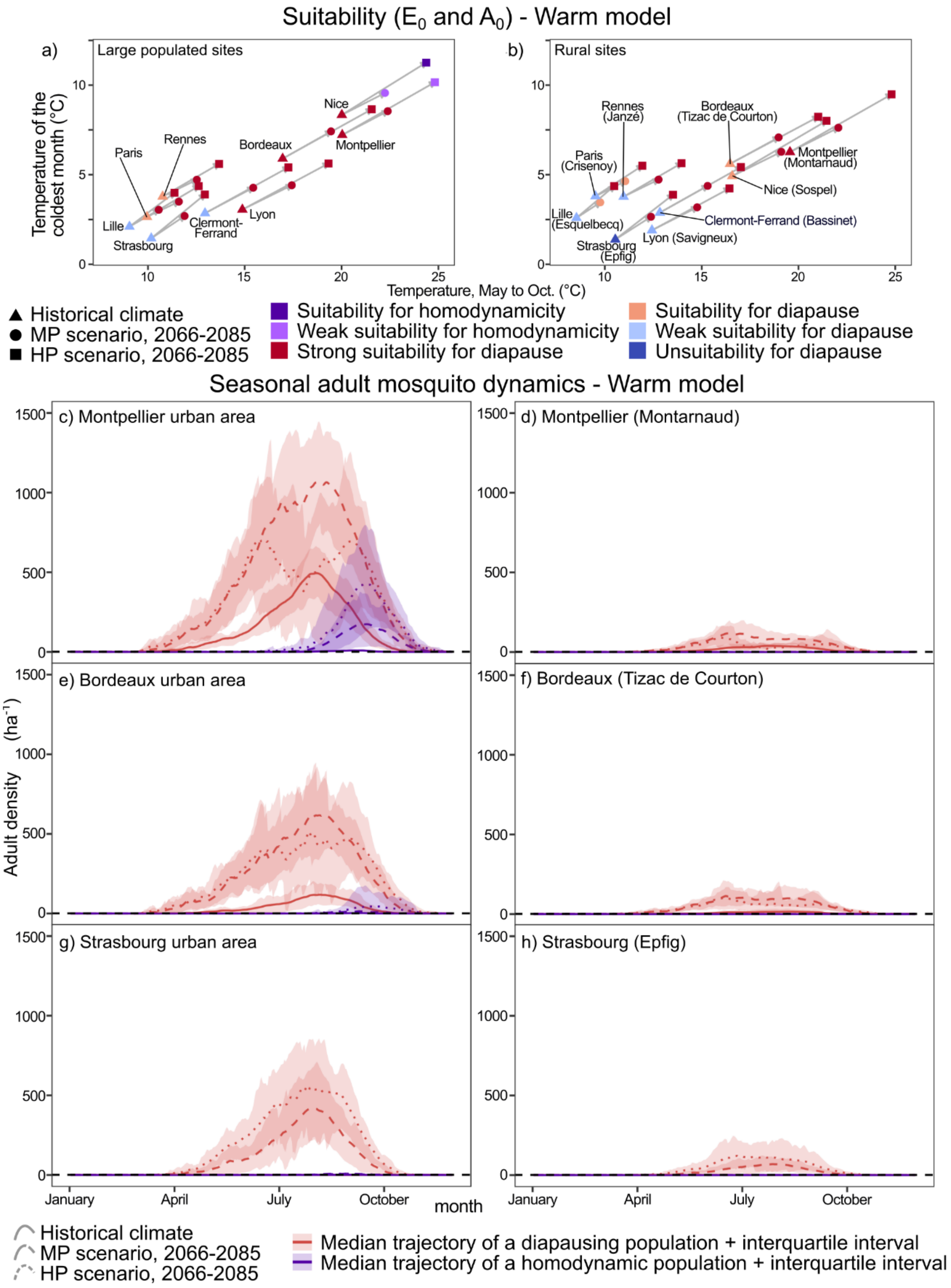


*Figure 31 - Projected change in suitability of the establishment of diapausing homodynamic Asian tiger mosquitoes in urban and rural areas and their seasonal dynamics depending on the time horizon and scenario, for the warm model.*

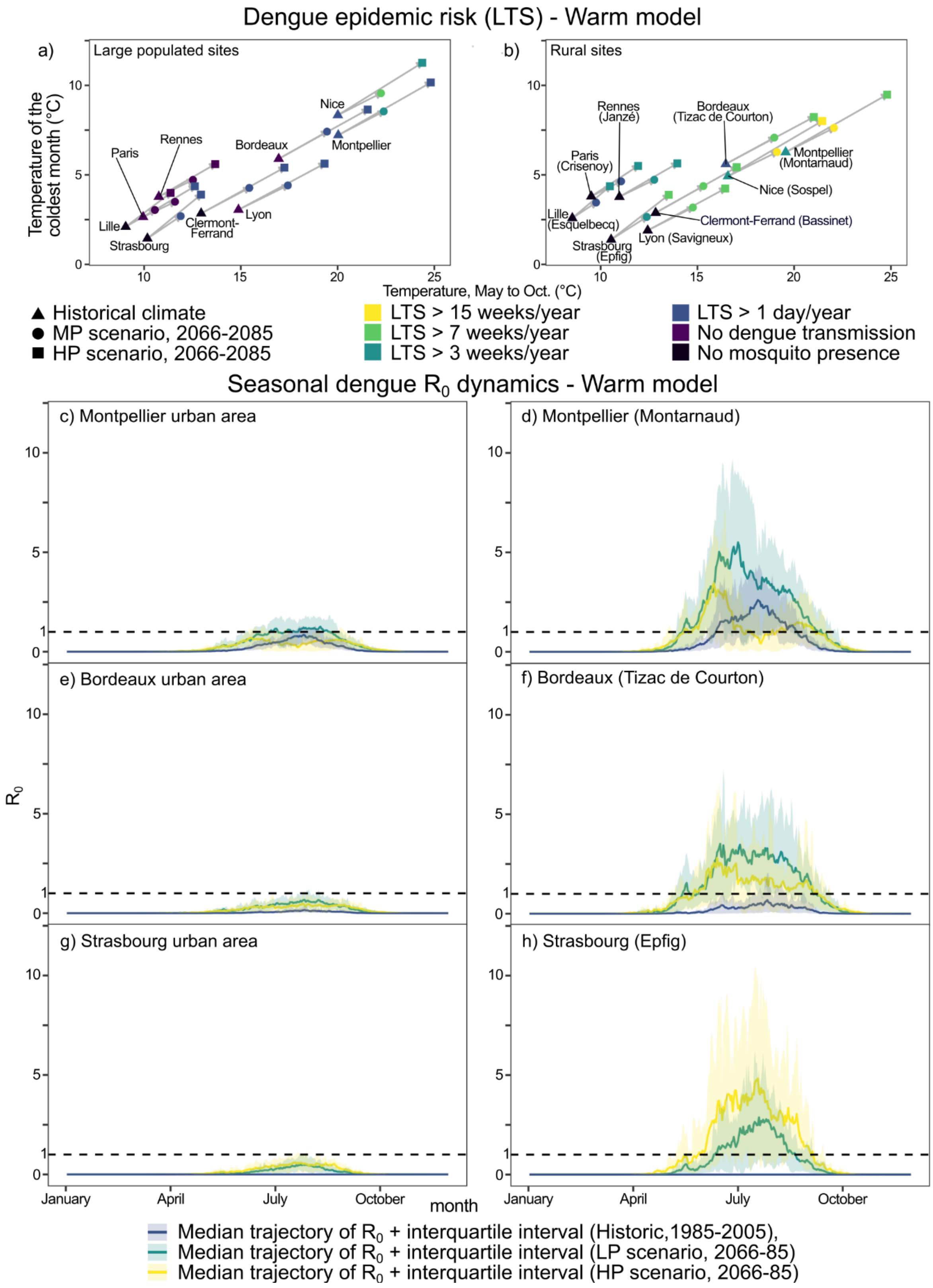


*Figure 32 - Projected change in the duration of the dengue transmission risk period in urban and rural areas and associated seasonal dynamics depending on the time horizon and scenario (warm model).*

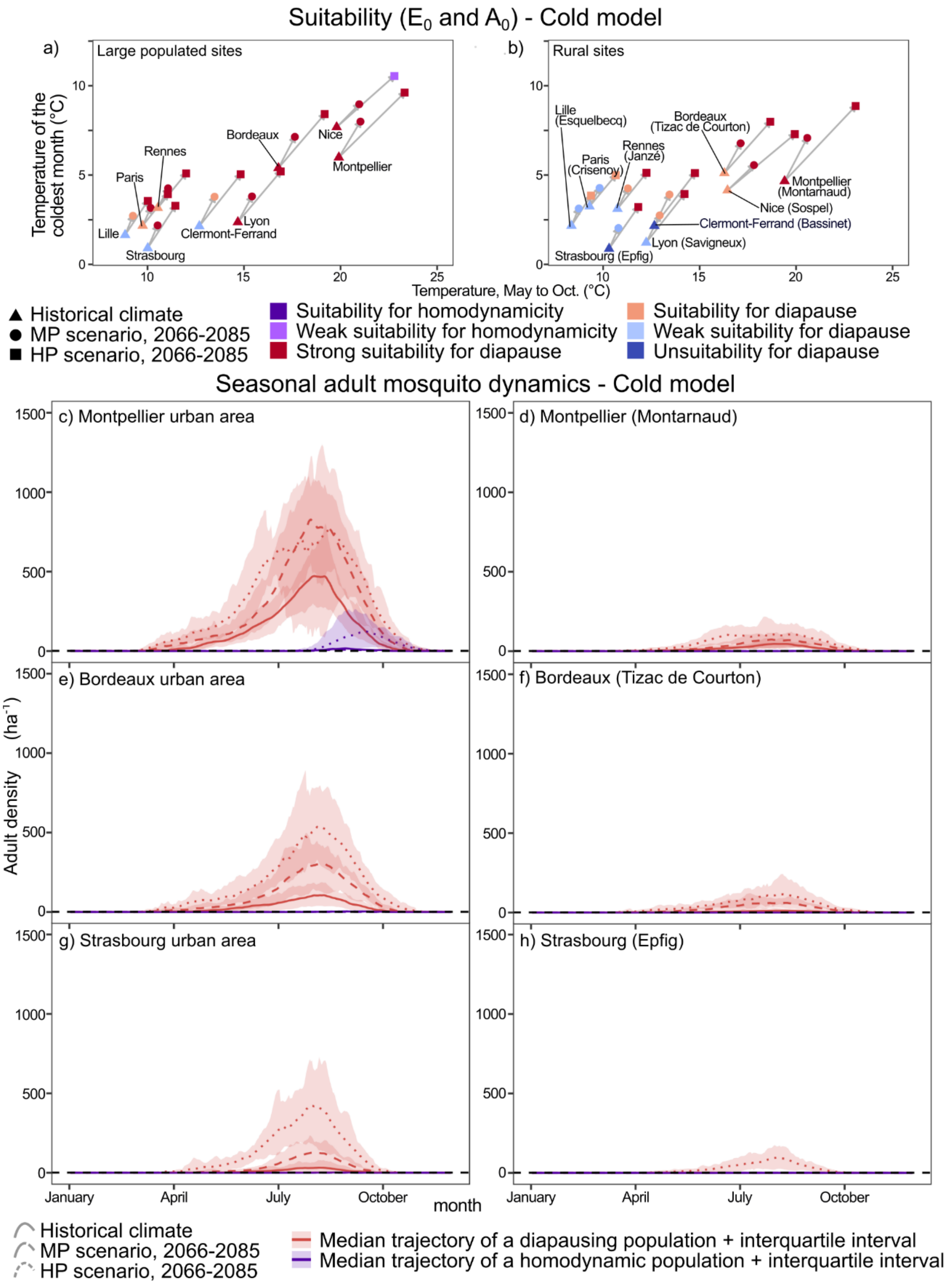


*Figure 33 - Projected change in suitability of the establishment of diapausing homodynamic Asian tiger mosquitoes in urban and rural areas and their seasonal dynamics depending on the time horizon and scenario, for the cold model.*

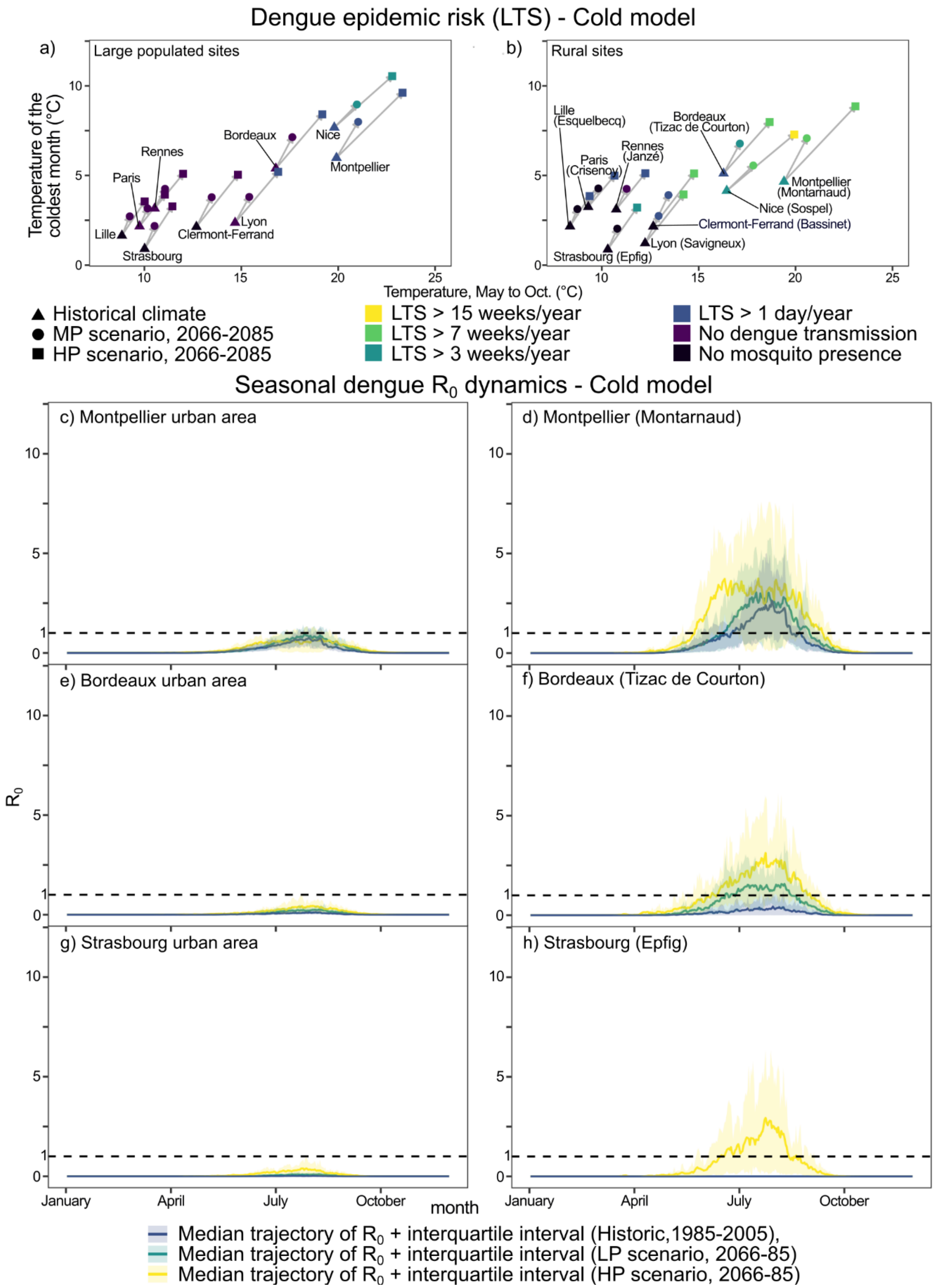


*Figure 34 - Projected change in the duration of the dengue transmission risk period in urban and rural areas and associated seasonal dynamics depending on the time horizon and scenario (cold model).*

| Site | Scenario | T Jan. (°C) | T May to Oct.(°C) | $E_0$ | $A_0$ | LTS (days) | Model |
|---|---|---|---|---|---|---|---|
| Bordeaux | 1986-2005 | 5.4 | 16.8 | 3.27e+01 | 1.42e-10 | 0.0 | cold |
| Bordeaux | 1986-2005 | 5.6 | 16.5 | 2.94e+01 | 1.08e-10 | 0.0 | inter. |
| Bordeaux | 1986-2005 | 5.9 | 17.0 | 3.91e+01 | 8.84e-11 | 0.0 | warm |
| Bordeaux | HP, 2066-85 | 8.4 | 19.2 | 3.84e+03 | 1.83e-05 | 1.2 | cold |
| Bordeaux | HP, 2066-85 | 8.1 | 19.5 | 6.50e+03 | 1.72e-04 | 2.1 | inter. |
| Bordeaux | HP, 2066-85 | 8.7 | 21.6 | 1.60e+04 | 1.39e-03 | 2.9 | warm |
| Bordeaux | MP, 2066-85 | 7.1 | 17.6 | 4.74e+02 | 8.48e-08 | 0.0 | cold |
| Bordeaux | MP, 2066-85 | 6.5 | 18.2 | 9.72e+02 | 1.39e-07 | 0.0 | inter. |
| Bordeaux | MP, 2066-85 | 7.4 | 19.5 | 8.39e+03 | 3.29e-05 | 4.2 | warm |
| Bordeaux (Tizac de Courton) | 1986-2005 | 5.1 | 16.3 | 2.80e+00 | 3.30e-10 | 5.0 | cold |
| Bordeaux (Tizac de Courton) | 1986-2005 | 5.4 | 16.0 | 3.73e+00 | 9.53e-10 | 13.8 | inter. |
| Bordeaux (Tizac de Courton) | 1986-2005 | 5.6 | 16.5 | 3.13e+00 | 2.20e-10 | 8.8 | warm |
| Bordeaux (Tizac de Courton) | HP, 2066-85 | 8.0 | 18.7 | 1.37e+02 | 9.79e-06 | 75.3 | cold |
| Bordeaux (Tizac de Courton) | HP, 2066-85 | 7.7 | 19.0 | 2.33e+02 | 7.04e-05 | 94.8 | inter. |
| Bordeaux (Tizac de Courton) | HP, 2066-85 | 8.2 | 21.0 | 2.39e+02 | 1.27e-04 | 74.8 | warm |
| Bordeaux (Tizac de Courton) | MP, 2066-85 | 6.8 | 17.1 | 2.80e+01 | 1.11e-07 | 43.0 | cold |
| Bordeaux (Tizac de Courton) | MP, 2066-85 | 6.2 | 17.8 | 5.14e+01 | 4.29e-07 | 67.2 | inter. |
| Bordeaux (Tizac de Courton) | MP, 2066-85 | 7.1 | 19.0 | 2.28e+02 | 1.39e-05 | 102.8 | warm |
| Clermont-Ferrand | 1986-2005 | 2.1 | 12.7 | 1.95e-01 | 2.84e-16 | 0.0 | cold |
| Clermont-Ferrand | 1986-2005 | 2.5 | 12.7 | 5.41e-01 | 3.74e-16 | 0.0 | inter. |
| Clermont-Ferrand | 1986-2005 | 2.8 | 12.9 | 4.11e-01 | 5.51e-16 | 0.0 | warm |
| Clermont-Ferrand | HP, 2066-85 | 5.0 | 14.8 | 2.55e+01 | 1.80e-10 | 0.2 | cold |
| Clermont-Ferrand | HP, 2066-85 | 4.8 | 15.3 | 4.27e+02 | 3.40e-09 | 3.4 | inter. |
| Clermont-Ferrand | HP, 2066-85 | 5.4 | 17.3 | 3.05e+03 | 7.08e-08 | 4.8 | warm |
| Clermont-Ferrand | MP, 2066-85 | 3.8 | 13.5 | 9.50e+00 | 8.11e-15 | 0.0 | cold |
| Clermont-Ferrand | MP, 2066-85 | 3.6 | 14.2 | 2.35e+01 | 9.24e-13 | 0.0 | inter. |
| Clermont-Ferrand | MP, 2066-85 | 4.3 | 15.4 | 4.07e+02 | 2.74e-10 | 3.1 | warm |
| Clermont-Ferrand (Bassinet) | 1986-2005 | 2.2 | 12.7 | 4.68e-02 | 2.92e-15 | 0.0 | cold |
| Clermont-Ferrand (Bassinet) | 1986-2005 | 2.7 | 12.6 | 2.20e-01 | 7.50e-15 | 0.8 | inter. |
| Clermont-Ferrand (Bassinet) | 1986-2005 | 2.9 | 12.8 | 1.35e-01 | 2.89e-15 | 0.0 | warm |
| Clermont-Ferrand (Bassinet) | HP, 2066-85 | 5.1 | 14.8 | 2.38e+01 | 2.98e-09 | 58.5 | cold |
| Clermont-Ferrand (Bassinet) | HP, 2066-85 | 4.6 | 15.1 | 4.60e+01 | 1.91e-08 | 90.7 | inter. |
| Clermont-Ferrand (Bassinet) | HP, 2066-85 | 5.4 | 17.0 | 7.67e+01 | 2.23e-07 | 78.2 | warm |
| Clermont-Ferrand (Bassinet) | MP, 2066-85 | 3.9 | 13.4 | 1.62e+00 | 9.82e-13 | 6.3 | cold |
| Clermont-Ferrand (Bassinet) | MP, 2066-85 | 3.5 | 14.0 | 3.41e+00 | 4.42e-11 | 19.4 | inter. |
| Clermont-Ferrand (Bassinet) | MP, 2066-85 | 4.4 | 15.3 | 3.10e+01 | 1.84e-09 | 75.4 | warm |
| Grenoble | 1986-2005 | 1.6 | 14.3 | 6.48e+00 | 3.34e-16 | 0.0 | cold |
| Grenoble | 1986-2005 | 1.6 | 14.3 | 4.19e+00 | 3.27e-16 | 0.0 | inter. |
| Grenoble | 1986-2005 | 2.2 | 14.5 | 6.33e+00 | 2.82e-15 | 0.0 | warm |
| Grenoble | HP, 2066-85 | 4.4 | 16.5 | 8.10e+02 | 3.80e-10 | 0.5 | cold |
| Grenoble | HP, 2066-85 | 4.1 | 17.0 | 2.22e+03 | 3.90e-08 | 7.0 | inter. |
| Grenoble | HP, 2066-85 | 4.9 | 18.8 | 9.03e+03 | 3.18e-07 | 6.6 | warm |
| Grenoble | MP, 2066-85 | 3.1 | 15.2 | 2.63e+01 | 3.65e-14 | 0.0 | cold |
| Grenoble | MP, 2066-85 | 3.5 | 15.7 | 3.30e+02 | 3.53e-11 | 0.8 | inter. |
| Grenoble | MP, 2066-85 | 3.7 | 16.9 | 1.47e+03 | 6.49e-10 | 4.1 | warm |
| Grenoble (La Motte d'Aveillans) | 1986-2005 | -0.7 | 11.9 | 1.52e-04 | 1.00e-10 | 0.0 | cold |
| Grenoble (La Motte d'Aveillans) | 1986-2005 | -0.3 | 11.8 | 2.30e-04 | 1.00e-10 | 0.0 | inter. |
| Grenoble (La Motte d'Aveillans) | 1986-2005 | 0.4 | 12.0 | 1.34e-04 | 2.22e-17 | 0.0 | warm |
| Grenoble (La Motte d'Aveillans) | HP, 2066-85 | 2.6 | 14.7 | 3.46e-01 | 4.53e-15 | 0.0 | cold |
| Grenoble (La Motte d'Aveillans) | HP, 2066-85 | 2.2 | 14.8 | 2.83e-01 | 6.65e-15 | 0.0 | inter. |
| Grenoble (La Motte d'Aveillans) | HP, 2066-85 | 3.3 | 16.7 | 2.15e+00 | 4.19e-13 | 6.7 | warm |
| Grenoble (La Motte d'Aveillans) | MP, 2066-85 | 1.2 | 13.0 | 4.11e-04 | 1.00e-10 | 0.0 | cold |
| Grenoble (La Motte d'Aveillans) | MP, 2066-85 | 1.5 | 13.4 | 5.74e-02 | 1.54e-16 | 0.0 | inter. |
| Grenoble (La Motte d'Aveillans) | MP, 2066-85 | 2.1 | 14.6 | 4.38e-02 | 4.54e-16 | 0.0 | warm |
| Lille | 1986-2005 | 1.6 | 8.8 | 5.69e-01 | 2.97e-15 | 0.0 | cold |
| Lille | 1986-2005 | 2.2 | 8.8 | 1.09e+00 | 2.15e-15 | 0.0 | inter. |
| Lille | 1986-2005 | 2.1 | 9.0 | 2.07e-01 | 4.19e-15 | 0.0 | warm |
| Lille | HP, 2066-85 | 3.6 | 10.0 | 7.23e+01 | 9.20e-11 | 0.0 | cold |
| Lille | HP, 2066-85 | 3.5 | 10.3 | 9.77e+01 | 2.30e-09 | 0.0 | inter. |
| Lille | HP, 2066-85 | 4.0 | 11.4 | 1.97e+03 | 6.06e-08 | 0.2 | warm |
| Lille | MP, 2066-85 | 2.7 | 9.2 | 2.79e+00 | 2.04e-14 | 0.0 | cold |
| Lille | MP, 2066-85 | 2.6 | 9.7 | 1.84e+01 | 1.27e-12 | 0.0 | inter. |
| Lille | MP, 2066-85 | 3.0 | 10.6 | 2.44e+02 | 4.65e-10 | 0.0 | warm |
| Lille (Esquelbecq) | 1986-2005 | 2.2 | 8.4 | 2.82e-01 | 3.79e-13 | 0.0 | cold |
| Lille (Esquelbecq) | 1986-2005 | 2.7 | 8.4 | 6.32e-01 | 1.64e-12 | 0.0 | inter. |
| Lille (Esquelbecq) | 1986-2005 | 2.6 | 8.5 | 1.47e-01 | 1.50e-13 | 0.0 | warm |
| Lille (Esquelbecq) | HP, 2066-85 | 3.8 | 9.4 | 5.17e+00 | 4.11e-09 | 3.4 | cold |
| Lille (Esquelbecq) | HP, 2066-85 | 3.9 | 9.6 | 6.19e+00 | 2.74e-08 | 6.7 | inter. |
| Lille (Esquelbecq) | HP, 2066-85 | 4.4 | 10.5 | 2.80e+01 | 2.17e-07 | 42.1 | warm |
| Lille (Esquelbecq) | MP, 2066-85 | 3.1 | 8.7 | 8.61e-01 | 1.36e-11 | 0.0 | cold |
| Lille (Esquelbecq) | MP, 2066-85 | 3.1 | 9.1 | 2.21e+00 | 2.76e-10 | 0.0 | inter. |

| Site | Scenario | T Jan. (°C) | T May to Oct.(°C) | $E_0$ | $A_0$ | LTS (days) | Model |
|---|---|---|---|---|---|---|---|
| Lille (Esquelbecq) | MP, 2066-85 | 3.5 | 9.8 | 8.51e+00 | 7.27e-09 | 12.9 | warm |
| Lyon | 1986-2005 | 2.4 | 14.7 | 6.51e+01 | 9.39e-14 | 0.0 | cold |
| Lyon | 1986-2005 | 2.7 | 14.6 | 5.77e+01 | 1.08e-13 | 0.1 | inter. |
| Lyon | 1986-2005 | 3.1 | 14.9 | 5.49e+01 | 1.34e-13 | 0.0 | warm |
| Lyon | HP, 2066-85 | 5.2 | 16.9 | 4.93e+03 | 1.51e-07 | 2.1 | cold |
| Lyon | HP, 2066-85 | 4.7 | 17.2 | 7.16e+03 | 7.07e-07 | 7.0 | inter. |
| Lyon | HP, 2066-85 | 5.6 | 19.3 | 9.04e+03 | 5.66e-06 | 5.0 | warm |
| Lyon | MP, 2066-85 | 3.8 | 15.4 | 4.23e+02 | 2.74e-11 | 0.1 | cold |
| Lyon | MP, 2066-85 | 3.9 | 16.1 | 1.78e+03 | 1.27e-09 | 2.3 | inter. |
| Lyon | MP, 2066-85 | 4.4 | 17.4 | 6.22e+03 | 7.03e-08 | 5.7 | warm |
| Lyon (Savigneux) | 1986-2005 | 1.2 | 12.2 | 1.86e-01 | 3.94e-16 | 0.0 | cold |
| Lyon (Savigneux) | 1986-2005 | 1.5 | 12.2 | 2.45e-01 | 4.68e-16 | 0.0 | inter. |
| Lyon (Savigneux) | 1986-2005 | 1.9 | 12.4 | 1.91e-01 | 4.13e-16 | 0.0 | warm |
| Lyon (Savigneux) | HP, 2066-85 | 3.9 | 14.2 | 3.84e+01 | 3.44e-10 | 58.3 | cold |
| Lyon (Savigneux) | HP, 2066-85 | 3.6 | 14.6 | 4.65e+01 | 9.23e-10 | 80.4 | inter. |
| Lyon (Savigneux) | HP, 2066-85 | 4.2 | 16.4 | 1.08e+02 | 1.52e-08 | 76.7 | warm |
| Lyon (Savigneux) | MP, 2066-85 | 2.7 | 12.9 | 1.85e+00 | 2.32e-14 | 4.8 | cold |
| Lyon (Savigneux) | MP, 2066-85 | 2.7 | 13.6 | 9.99e+00 | 2.52e-12 | 40.6 | inter. |
| Lyon (Savigneux) | MP, 2066-85 | 3.2 | 14.8 | 2.31e+01 | 1.16e-10 | 62.6 | warm |
| Marseille | 1986-2005 | 6.4 | 20.9 | 2.13e+03 | 1.81e-05 | 1.1 | cold |
| Marseille | 1986-2005 | 7.3 | 19.4 | 1.75e+03 | 5.28e-06 | 0.5 | inter. |
| Marseille | 1986-2005 | 9.7 | 21.7 | 2.12e+03 | 1.71e-05 | 0.8 | warm |
| Marseille | HP, 2066-85 | 12.5 | 24.6 | 3.31e+04 | 5.36e-02 | 16.6 | cold |
| Marseille | HP, 2066-85 | 12.2 | 23.0 | 4.55e+04 | 1.61e-01 | 37.5 | inter. |
| Marseille | HP, 2066-85 | 11.7 | 25.2 | 4.88e+04 | 3.38e-01 | 19.0 | warm |
| Marseille | MP, 2066-85 | 9.2 | 21.2 | 9.02e+03 | 7.73e-04 | 8.9 | cold |
| Marseille | MP, 2066-85 | 10.1 | 21.6 | 1.95e+04 | 1.58e-03 | 20.4 | inter. |
| Marseille | MP, 2066-85 | 9.7 | 21.2 | 3.52e+04 | 2.21e-02 | 43.5 | warm |
| Marseille (Le Tholonet) | 1986-2005 | 3.8 | 18.6 | 2.36e+00 | 1.10e-10 | 5.9 | cold |
| Marseille (Le Tholonet) | 1986-2005 | 5.0 | 18.3 | 1.77e+00 | 8.09e-11 | 0.8 | inter. |
| Marseille (Le Tholonet) | 1986-2005 | 5.7 | 18.9 | 2.77e+00 | 5.94e-11 | 1.9 | warm |
| Marseille (Le Tholonet) | HP, 2066-85 | 7.9 | 22.5 | 9.34e+01 | 5.26e-06 | 74.5 | cold |
| Marseille (Le Tholonet) | HP, 2066-85 | 7.6 | 22.1 | 1.08e+02 | 1.55e-05 | 90.8 | inter. |
| Marseille (Le Tholonet) | HP, 2066-85 | 8.4 | 24.3 | 1.49e+02 | 3.06e-05 | 70.0 | warm |
| Marseille (Le Tholonet) | MP, 2066-85 | 5.8 | 19.8 | 6.60e+00 | 1.99e-08 | 30.6 | cold |
| Marseille (Le Tholonet) | MP, 2066-85 | 6.8 | 20.6 | 3.35e+01 | 2.48e-07 | 67.8 | inter. |
| Marseille (Le Tholonet) | MP, 2066-85 | 6.5 | 21.2 | 8.42e+01 | 1.94e-06 | 85.8 | warm |
| Montpellier | 1986-2005 | 6.0 | 19.9 | 2.52e+03 | 1.93e-06 | 5.6 | cold |
| Montpellier | 1986-2005 | 6.8 | 19.8 | 2.20e+03 | 1.06e-06 | 8.8 | inter. |
| Montpellier | 1986-2005 | 7.2 | 20.0 | 2.96e+03 | 2.03e-06 | 8.4 | warm |
| Montpellier | HP, 2066-85 | 9.6 | 23.3 | 2.67e+04 | 1.54e-02 | 10.2 | cold |
| Montpellier | HP, 2066-85 | 9.1 | 23.1 | 4.85e+04 | 7.84e-02 | 30.6 | inter. |
| Montpellier | HP, 2066-85 | 10.2 | 24.8 | 4.24e+04 | 2.21e-01 | 12.9 | warm |
| Montpellier | MP, 2066-85 | 8.0 | 21.0 | 9.48e+03 | 1.61e-04 | 12.8 | cold |
| Montpellier | MP, 2066-85 | 8.3 | 21.6 | 2.30e+04 | 5.08e-04 | 25.2 | inter. |
| Montpellier | MP, 2066-85 | 8.5 | 22.4 | 3.88e+04 | 8.69e-03 | 42.0 | warm |
| Montpellier (Montarnaud) | 1986-2005 | 4.7 | 19.4 | 1.09e+01 | 6.58e-09 | 49.5 | cold |
| Montpellier (Montarnaud) | 1986-2005 | 5.8 | 19.3 | 1.07e+01 | 1.39e-08 | 55.6 | inter. |
| Montpellier (Montarnaud) | 1986-2005 | 6.3 | 19.6 | 1.49e+01 | 4.60e-09 | 56.0 | warm |
| Montpellier (Montarnaud) | HP, 2066-85 | 8.9 | 23.1 | 2.36e+02 | 8.39e-05 | 96.0 | cold |
| Montpellier (Montarnaud) | HP, 2066-85 | 8.5 | 23.0 | 4.26e+02 | 4.48e-04 | 123.2 | inter. |
| Montpellier (Montarnaud) | HP, 2066-85 | 9.5 | 24.8 | 1.77e+02 | 3.51e-04 | 72.8 | warm |
| Montpellier (Montarnaud) | MP, 2066-85 | 7.1 | 20.6 | 3.53e+01 | 1.03e-06 | 67.6 | cold |
| Montpellier (Montarnaud) | MP, 2066-85 | 7.4 | 21.3 | 1.61e+02 | 8.50e-06 | 112.7 | inter. |
| Montpellier (Montarnaud) | MP, 2066-85 | 7.6 | 22.1 | 2.24e+02 | 4.57e-05 | 111.0 | warm |
| Nantes | 1986-2005 | 3.5 | 11.5 | 4.20e+00 | 5.30e-13 | 0.0 | cold |
| Nantes | 1986-2005 | 4.0 | 11.3 | 4.35e+00 | 6.04e-13 | 0.0 | inter. |
| Nantes | 1986-2005 | 4.1 | 11.7 | 6.34e+00 | 1.47e-12 | 0.0 | warm |
| Nantes | HP, 2066-85 | 5.6 | 13.1 | 3.74e+02 | 5.30e-08 | 0.0 | cold |
| Nantes | HP, 2066-85 | 5.5 | 13.5 | 5.76e+02 | 8.24e-07 | 0.0 | inter. |
| Nantes | HP, 2066-85 | 5.9 | 14.9 | 4.67e+03 | 7.57e-06 | 0.0 | warm |
| Nantes | MP, 2066-85 | 4.7 | 12.1 | 2.63e+01 | 1.90e-10 | 0.0 | cold |
| Nantes | MP, 2066-85 | 4.4 | 12.7 | 9.12e+01 | 6.16e-10 | 0.0 | inter. |
| Nantes | MP, 2066-85 | 5.1 | 13.6 | 9.47e+02 | 1.69e-07 | 0.0 | warm |
| Nantes (Saint-Philbert-de-Grand-Lieu) | 1986-2005 | 3.8 | 12.1 | 1.01e+00 | 1.39e-11 | 0.0 | cold |
| Nantes (Saint-Philbert-de-Grand-Lieu) | 1986-2005 | 4.3 | 11.9 | 1.47e+00 | 7.19e-11 | 0.6 | inter. |
| Nantes (Saint-Philbert-de-Grand-Lieu) | 1986-2005 | 4.5 | 12.3 | 1.55e+00 | 3.36e-11 | 0.1 | warm |
| Nantes (Saint-Philbert-de-Grand-Lieu) | HP, 2066-85 | 5.9 | 13.7 | 2.35e+01 | 3.26e-07 | 35.2 | cold |
| Nantes (Saint-Philbert-de-Grand-Lieu) | HP, 2066-85 | 5.8 | 14.2 | 4.58e+01 | 3.15e-06 | 42.8 | inter. |
| Nantes (Saint-Philbert-de-Grand-Lieu) | HP, 2066-85 | 6.3 | 15.7 | 1.15e+02 | 1.00e-05 | 66.0 | warm |
| Nantes (Saint-Philbert-de-Grand-Lieu) | MP, 2066-85 | 5.0 | 12.7 | 4.84e+00 | 3.55e-09 | 6.8 | cold |

| Site | Scenario | T Jan. (°C) | T May to Oct.(°C) | $E_0$ | $A_0$ | LTS (days) | Model |
|---|---|---|---|---|---|---|---|
| Nantes (Saint-Philbert-de-Grand-Lieu) | MP, 2066-85 | 4.7 | 13.3 | 1.11e+01 | 1.67e-08 | 17.2 | inter. |
| Nantes (Saint-Philbert-de-Grand-Lieu) | MP, 2066-85 | 5.4 | 14.3 | 5.12e+01 | 8.27e-07 | 60.8 | warm |
| Nice | 1986-2005 | 7.7 | 19.8 | 6.54e+03 | 5.95e-04 | 2.5 | cold |
| Nice | 1986-2005 | 8.0 | 19.8 | 7.26e+03 | 1.60e-04 | 5.0 | inter. |
| Nice | 1986-2005 | 8.3 | 20.0 | 7.91e+03 | 6.41e-04 | 5.4 | warm |
| Nice | HP, 2066-85 | 10.5 | 22.8 | 7.03e+04 | 7.44e-01 | 34.2 | cold |
| Nice | HP, 2066-85 | 9.9 | 22.9 | 1.05e+05 | 2.67e+00 | 52.4 | inter. |
| Nice | HP, 2066-85 | 11.3 | 24.4 | 1.16e+05 | 9.80e+00 | 49.0 | warm |
| Nice | MP, 2066-85 | 9.0 | 21.0 | 2.68e+04 | 1.79e-02 | 23.6 | cold |
| Nice | MP, 2066-85 | 9.6 | 21.5 | 5.32e+04 | 3.80e-02 | 43.9 | inter. |
| Nice | MP, 2066-85 | 9.6 | 22.2 | 7.90e+04 | 5.00e-01 | 70.2 | warm |
| Nice (Sospel) | 1986-2005 | 4.1 | 16.4 | 8.17e+00 | 5.28e-09 | 38.8 | cold |
| Nice (Sospel) | 1986-2005 | 4.6 | 16.4 | 8.54e+00 | 5.63e-09 | 43.8 | inter. |
| Nice (Sospel) | 1986-2005 | 4.9 | 16.6 | 8.67e+00 | 4.50e-09 | 36.2 | warm |
| Nice (Sospel) | HP, 2066-85 | 7.3 | 19.9 | 4.00e+02 | 4.02e-05 | 129.8 | cold |
| Nice (Sospel) | HP, 2066-85 | 6.7 | 19.8 | 2.70e+02 | 8.06e-05 | 130.2 | inter. |
| Nice (Sospel) | HP, 2066-85 | 8.0 | 21.4 | 6.52e+02 | 4.31e-04 | 143.3 | warm |
| Nice (Sospel) | MP, 2066-85 | 5.6 | 17.8 | 6.10e+01 | 6.38e-07 | 97.0 | cold |
| Nice (Sospel) | MP, 2066-85 | 6.3 | 18.3 | 9.56e+01 | 3.23e-06 | 114.3 | inter. |
| Nice (Sospel) | MP, 2066-85 | 6.3 | 19.1 | 1.78e+02 | 1.45e-05 | 127.1 | warm |
| Paris-est | 1986-2005 | 2.2 | 9.8 | 3.00e+00 | 5.42e-13 | 0.0 | cold |
| Paris-est | 1986-2005 | 2.6 | 9.7 | 8.69e+00 | 2.45e-13 | 0.0 | inter. |
| Paris-est | 1986-2005 | 2.6 | 9.9 | 6.34e+00 | 1.91e-13 | 0.0 | warm |
| Paris-est | HP, 2066-85 | 3.9 | 11.1 | 5.22e+02 | 2.00e-08 | 0.0 | cold |
| Paris-est | HP, 2066-85 | 3.9 | 11.5 | 1.13e+03 | 3.43e-07 | 0.4 | inter. |
| Paris-est | HP, 2066-85 | 4.4 | 12.6 | 6.20e+03 | 2.79e-06 | 2.0 | warm |
| Paris-est | MP, 2066-85 | 3.2 | 10.2 | 3.71e+01 | 1.87e-11 | 0.0 | cold |
| Paris-est | MP, 2066-85 | 3.1 | 10.8 | 1.48e+02 | 1.82e-10 | 0.0 | inter. |
| Paris-est | MP, 2066-85 | 3.5 | 11.6 | 1.33e+03 | 5.60e-08 | 0.7 | warm |
| Paris-est (Crisenoy) | 1986-2005 | 3.2 | 9.3 | 1.52e-01 | 1.50e-12 | 0.0 | cold |
| Paris-est (Crisenoy) | 1986-2005 | 3.8 | 9.2 | 2.87e-01 | 3.82e-12 | 0.0 | inter. |
| Paris-est (Crisenoy) | 1986-2005 | 3.8 | 9.5 | 1.62e-01 | 1.75e-12 | 0.0 | warm |
| Paris-est (Crisenoy) | HP, 2066-85 | 5.0 | 10.7 | 4.10e+00 | 1.77e-08 | 2.5 | cold |
| Paris-est (Crisenoy) | HP, 2066-85 | 5.1 | 10.9 | 5.16e+00 | 1.03e-07 | 6.3 | inter. |
| Paris-est (Crisenoy) | HP, 2066-85 | 5.5 | 11.9 | 2.50e+01 | 7.66e-07 | 45.6 | warm |
| Paris-est (Crisenoy) | MP, 2066-85 | 4.3 | 9.8 | 5.99e-01 | 1.74e-10 | 0.0 | cold |
| Paris-est (Crisenoy) | MP, 2066-85 | 4.2 | 10.3 | 1.57e+00 | 8.52e-10 | 0.0 | inter. |
| Paris-est (Crisenoy) | MP, 2066-85 | 4.6 | 11.1 | 5.43e+00 | 4.20e-08 | 12.2 | warm |
| Rennes | 1986-2005 | 3.2 | 10.5 | 1.66e+00 | 3.59e-13 | 0.0 | cold |
| Rennes | 1986-2005 | 3.7 | 10.4 | 2.25e+00 | 2.23e-13 | 0.0 | inter. |
| Rennes | 1986-2005 | 3.8 | 10.8 | 2.82e+00 | 6.21e-13 | 0.0 | warm |
| Rennes | HP, 2066-85 | 5.1 | 12.0 | 1.52e+02 | 2.17e-08 | 0.0 | cold |
| Rennes | HP, 2066-85 | 5.1 | 12.4 | 2.48e+02 | 2.65e-07 | 0.0 | inter. |
| Rennes | HP, 2066-85 | 5.6 | 13.7 | 2.87e+03 | 3.65e-06 | 0.0 | warm |
| Rennes | MP, 2066-85 | 4.3 | 11.1 | 1.17e+01 | 5.67e-11 | 0.0 | cold |
| Rennes | MP, 2066-85 | 4.1 | 11.7 | 3.92e+01 | 2.11e-10 | 0.0 | inter. |
| Rennes | MP, 2066-85 | 4.7 | 12.5 | 4.49e+02 | 8.69e-08 | 0.0 | warm |
| Rennes (Janzé) | 1986-2005 | 3.1 | 10.7 | 2.69e-01 | 8.25e-13 | 0.0 | cold |
| Rennes (Janzé) | 1986-2005 | 3.7 | 10.6 | 5.79e-01 | 3.15e-12 | 0.0 | inter. |
| Rennes (Janzé) | 1986-2005 | 3.8 | 11.0 | 4.49e-01 | 1.50e-12 | 0.0 | warm |
| Rennes (Janzé) | HP, 2066-85 | 5.1 | 12.2 | 1.00e+01 | 3.15e-08 | 7.5 | cold |
| Rennes (Janzé) | HP, 2066-85 | 5.1 | 12.7 | 1.45e+01 | 2.21e-07 | 12.8 | inter. |
| Rennes (Janzé) | HP, 2066-85 | 5.6 | 14.0 | 5.87e+01 | 1.33e-06 | 45.9 | warm |
| Rennes (Janzé) | MP, 2066-85 | 4.3 | 11.3 | 1.62e+00 | 1.59e-10 | 0.0 | cold |
| Rennes (Janzé) | MP, 2066-85 | 4.1 | 11.9 | 3.66e+00 | 9.59e-10 | 1.2 | inter. |
| Rennes (Janzé) | MP, 2066-85 | 4.7 | 12.8 | 1.48e+01 | 6.39e-08 | 21.6 | warm |
| Strasbourg | 1986-2005 | 0.9 | 10.0 | 7.88e-01 | 4.39e-15 | 0.0 | cold |
| Strasbourg | 1986-2005 | 1.5 | 10.1 | 2.75e+00 | 1.13e-15 | 0.0 | inter. |
| Strasbourg | 1986-2005 | 1.4 | 10.2 | 3.90e-01 | 8.29e-16 | 0.0 | warm |
| Strasbourg | HP, 2066-85 | 3.3 | 11.4 | 7.73e+02 | 9.99e-10 | 0.3 | cold |
| Strasbourg | HP, 2066-85 | 3.1 | 11.8 | 1.43e+03 | 9.28e-09 | 3.5 | inter. |
| Strasbourg | HP, 2066-85 | 3.9 | 12.9 | 6.10e+03 | 7.53e-08 | 4.6 | warm |
| Strasbourg | MP, 2066-85 | 2.2 | 10.5 | 2.11e+01 | 1.35e-14 | 0.0 | cold |
| Strasbourg | MP, 2066-85 | 2.4 | 11.2 | 1.87e+02 | 1.15e-11 | 0.1 | inter. |
| Strasbourg | MP, 2066-85 | 2.7 | 11.9 | 4.49e+02 | 1.17e-10 | 2.0 | warm |
| Strasbourg (Epfig) | 1986-2005 | 0.9 | 10.3 | 1.98e-02 | 3.43e-15 | 0.0 | cold |
| Strasbourg (Epfig) | 1986-2005 | 1.5 | 10.5 | 4.21e-01 | 1.34e-15 | 0.0 | inter. |
| Strasbourg (Epfig) | 1986-2005 | 1.4 | 10.5 | 8.27e-02 | 6.64e-16 | 0.0 | warm |
| Strasbourg (Epfig) | HP, 2066-85 | 3.2 | 11.8 | 1.98e+01 | 6.60e-11 | 47.6 | cold |
| Strasbourg (Epfig) | HP, 2066-85 | 3.0 | 12.3 | 8.58e+01 | 4.39e-09 | 88.1 | inter. |
| Strasbourg (Epfig) | HP, 2066-85 | 3.9 | 13.5 | 1.49e+02 | 1.96e-08 | 90.6 | warm |

| Site | Scenario | T Jan. (°C) | T May to Oct.(°C) | $E_0$ | $A_0$ | LTS (days) | Model |
|---|---|---|---|---|---|---|---|
| Strasbourg (Epfig) | MP, 2066-85 | 2.0 | 10.8 | 3.09e-01 | 6.04e-15 | 0.0 | cold |
| Strasbourg (Epfig) | MP, 2066-85 | 2.3 | 11.6 | 1.06e+01 | 9.45e-12 | 43.2 | inter. |
| Strasbourg (Epfig) | MP, 2066-85 | 2.7 | 12.4 | 1.77e+01 | 5.79e-11 | 55.7 | warm |
| Toulouse | 1986-2005 | 4.5 | 19.3 | 7.22e+01 | 2.19e-10 | 0.0 | cold |
| Toulouse | 1986-2005 | 5.9 | 18.2 | 8.75e+01 | 1.70e-10 | 0.1 | inter. |
| Toulouse | 1986-2005 | 7.1 | 19.9 | 1.64e+02 | 1.58e-10 | 0.1 | warm |
| Toulouse | HP, 2066-85 | 9.7 | 22.8 | 7.20e+03 | 5.13e-05 | 1.9 | cold |
| Toulouse | HP, 2066-85 | 9.4 | 22.3 | 1.07e+04 | 1.96e-04 | 5.0 | inter. |
| Toulouse | HP, 2066-85 | 9.5 | 25.2 | 1.40e+04 | 1.22e-03 | 1.9 | warm |
| Toulouse | MP, 2066-85 | 7.1 | 20.1 | 1.28e+03 | 2.86e-07 | 0.7 | cold |
| Toulouse | MP, 2066-85 | 6.3 | 20.8 | 2.28e+03 | 2.71e-07 | 0.8 | inter. |
| Toulouse | MP, 2066-85 | 7.7 | 21.7 | 1.26e+04 | 4.64e-05 | 6.3 | warm |
| Toulouse (Lavalette) | 1986-2005 | 4.8 | 18.7 | 5.08e+00 | 1.61e-10 | 9.6 | cold |
| Toulouse (Lavalette) | 1986-2005 | 5.7 | 18.1 | 6.65e+00 | 3.66e-10 | 32.6 | inter. |
| Toulouse (Lavalette) | 1986-2005 | 6.3 | 18.9 | 7.39e+00 | 1.09e-10 | 31.4 | warm |
| Toulouse (Lavalette) | HP, 2066-85 | 8.9 | 22.0 | 1.70e+02 | 8.45e-06 | 77.3 | cold |
| Toulouse (Lavalette) | HP, 2066-85 | 8.7 | 21.9 | 3.31e+02 | 3.83e-05 | 110.0 | inter. |
| Toulouse (Lavalette) | HP, 2066-85 | 9.0 | 24.7 | 1.76e+02 | 3.92e-05 | 66.8 | warm |
| Toulouse (Lavalette) | MP, 2066-85 | 7.2 | 19.7 | 3.79e+01 | 1.01e-07 | 54.0 | cold |
| Toulouse (Lavalette) | MP, 2066-85 | 6.5 | 20.3 | 5.88e+01 | 2.55e-07 | 76.2 | inter. |
| Toulouse (Lavalette) | MP, 2066-85 | 7.5 | 21.8 | 2.61e+02 | 5.95e-06 | 102.3 | warm |

*Table SI3 - Detail upon variation of climate suitability for the establishment of the mosquito (E0 and A0, to compare with the threshold of 1) and for the duration of the transmission season of dengue (LTS) per scenario and climate model.*

#### Relative importance of climate and population change

In these sensitivity experiments, the effect of climate change alone primarily explain both suitability for the establishment of the vector (Figure 35; compare with Figure 4 in the main text) and for dengue transmission risk (Figure 36; compare with Figure 5 in the main text), since the estimates of suitable areas in % are extremely close.

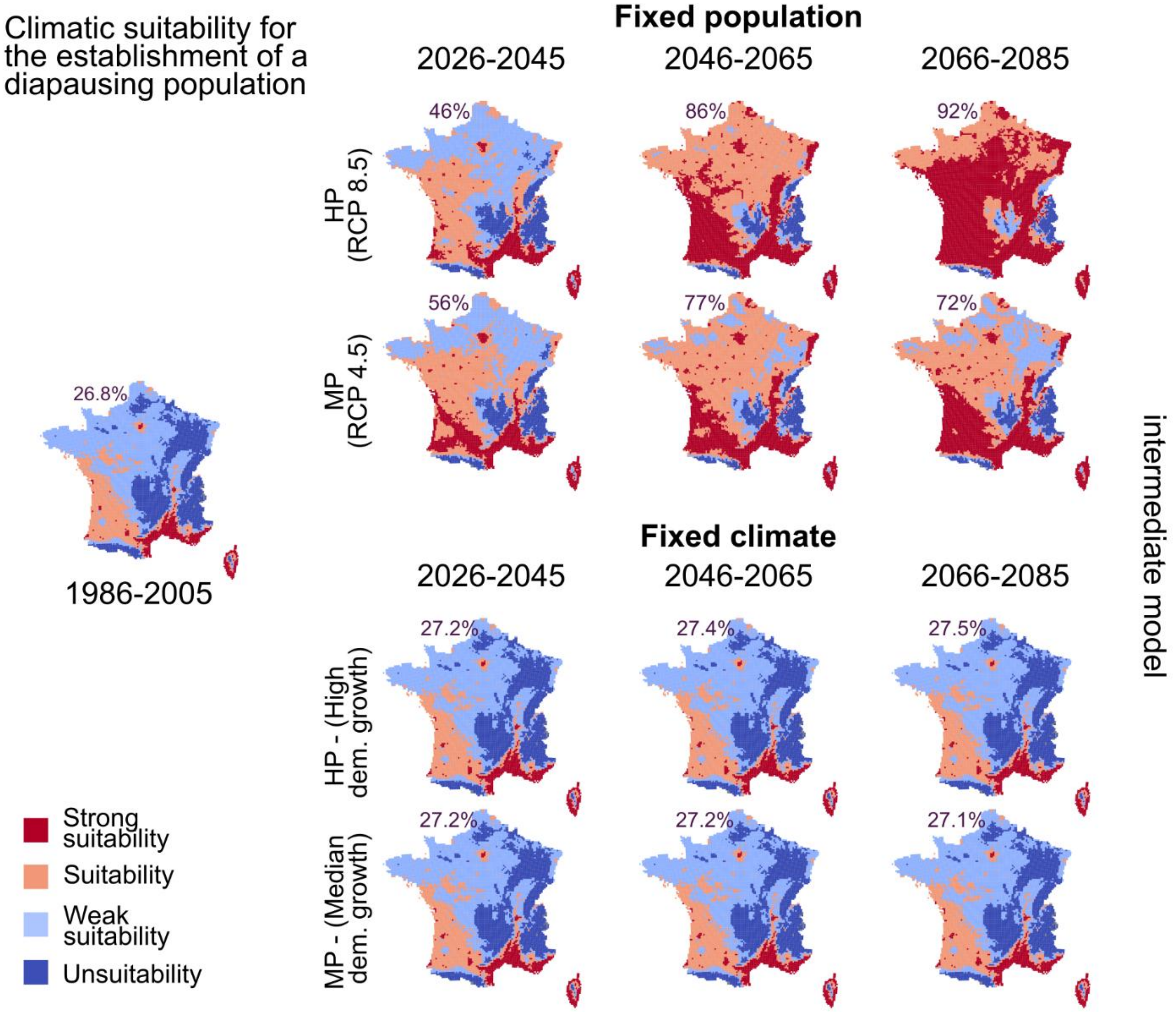


*Figure 35 - Relative importance of climate (upper line) and population changes (below) on spatial suitability for the establishment of Ae. albopictus.*

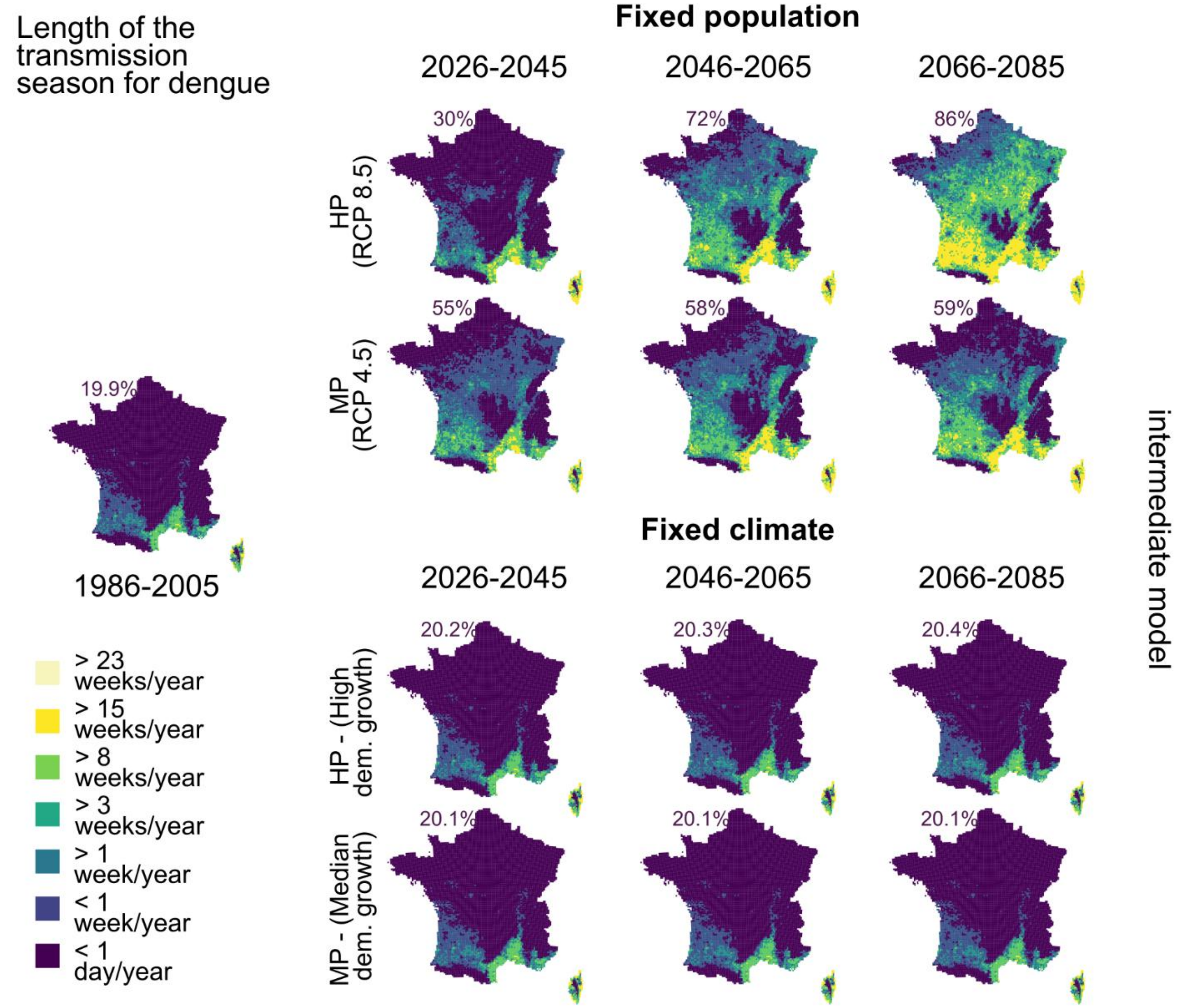


*Figure 36 - Relative importance of climate (upper line) and population changes (below) on the length of the dengue transmission season.*
